# Modified meiosis in the tardigrade *Hypsibius exemplaris* maintains heterozygosity across the genome

**DOI:** 10.64898/2026.03.11.711151

**Authors:** Adriana N. Coke, Siqin Aurora Zhang, Lillian D. Papell, Christina L. Burch, Bob Goldstein

## Abstract

In asexual reproduction, meiosis must be bypassed or altered to maintain ploidy from mother to daughter without fertilization. Most of the ways meiosis can be modified to this end are expected to reduce heterozygosity within individuals; however, many asexual species are highly heterozygous. Asexual reproduction is especially common among species of microscopic, desiccation-tolerant animals such as rotifers, nematodes, and tardigrades, but the cellular and genetic mechanisms underlying asexual reproduction have not been definitively documented in any species of tardigrade. Here, we show that the asexual tardigrade *Hypsibius exemplaris* fails to complete the cell division of meiosis I, followed by a complete meiosis II-like division, and reproduction proceeds without detectable loss of heterozygosity. We used a combined cytological and genomic approach to characterize the mechanism of reproduction and pattern of allele inheritance in this species. Furthermore, we identified heterozygous variants in a subset of transcriptionally active genes consistent with loss of function in one allele, suggesting that maintained heterozygosity in this species allowed divergence between alleles over time. This work establishes the meiotic mechanism and inheritance pattern of reproduction in *H. exemplaris*, which provides a framework for interpreting genetic variation in this organism as a laboratory model. Additionally, our finding that meiosis is modified in *H. exemplaris* via a mechanism that maintains heterozygosity across the genome adds to a growing list of asexual animals that have evolved to reproduce clonally despite the expected long-term costs.

**Article Summary:** Asexual animals must alter meiosis, a highly conserved process of two cell divisions normally used to make eggs and sperm. This study represents the first combined cytological and genetic characterization of how meiosis is modified in a tardigrade. The authors found that the model tardigrade *Hypsibius exemplaris* modifies meiosis by skipping cytokinesis of the first division, followed by a complete second division. They also found that this species preserves heterozygosity across the genome and from generation to generation. Finally, some genes show evidence of divergence between alleles, supporting a broader conclusion that maintaining heterozygosity influences how asexual species evolve.

## Introduction

Meiosis is a highly conserved process, ancestral to eukaryotes, that determines how genes are passed from parent to offspring (Speijer et al. 2015). A diploid cell undergoes two divisions during meiosis to produce a haploid gamete, and in sexual reproduction, diploidy is restored through fertilization with another haploid gamete. These canonical processes are the fundamental basis for Mendelian genetics, and nondisjunction or other deviations from typical meiosis can have dire consequences. However, to reproduce asexually, canonical meiosis must be bypassed or altered such that ploidy can be maintained from mother to daughter without fertilization. Therefore, understanding how meiosis is modified is central to understanding asexual reproduction in animals, or parthenogenesis. Most parthenogens modify meiosis rather than eliminating it (Blanc et al. 2025). Meiosis can be modified in several ways to maintain ploidy: one of the two meiotic divisions may be incomplete, there may be fusion of two gametes from a single meiosis, or there may be an extra round of DNA replication before, during, or after meiosis. Most of these mechanisms of asexual reproduction are predicted to result in immediate or progressive loss of heterozygosity within individuals, if meiosis still involves recombination (Blanc et al. 2025). Indeed, many meiotic parthenogens of non-hybrid origins have low levels of heterozygosity (Jaron et al. 2021).

However, observed heterozygosity in some asexual species has contradicted theoretical expectations. For example, the rotifer *Adineta vaga* (Simion et al. 2021; Terwagne et al. 2022), the nematode *Mesorhabditis belari* (Blanc et al. 2023), and the ant *Ooceraea biroi* (Lacy et al. 2024) all reproduce by abortive meiosis I parthenogenesis, where meiosis I begins but does not result in the formation of a polar body, followed by a complete meiosis II-like division. This meiosis modification results in assortment of non-sister chromatids to produce a diploid offspring and should, if meiotic crossing over occurs, result in loss of heterozygosity distal to crossover sites (Blanc et al. 2025); however, all three species maintain heterozygosity across the genome (Simion et al. 2021; Blanc et al. 2023; Lacy et al. 2024). In *A. vaga*, the underlying basis for heterozygosity maintenance remains unclear (Blanc et al. 2025), but in *M. belari* and *O. biroi*, this paradox has been resolved through the discovery of an additional meiosis modification: either both recombined or both non-recombined chromatids are retained in the egg, and hence heterozygosity is maintained at a genome scale (Blanc et al. 2023; Lacy et al. 2024). These examples demonstrate that substantial heterozygosity is stably maintained in some asexual animals despite meiosis modifications to maintain ploidy that are expected to cause loss of heterozygosity. Careful cytological and genetic characterization of meiosis modifications in more species is required to understand whether these examples are indicative of a larger evolutionary pattern.

When heterozygosity is maintained in asexual reproduction over sufficient evolutionary time, alleles of a single gene will accumulate mutations independently (Welch and Meselson 2000). Independent evolution between haplotypes has been demonstrated genome-wide for asexual oribatid mites (Brandt et al. 2021; Öztoprak et al. 2025). Haplotype divergence may lead to one allele becoming nonfunctional over evolutionary time due to relaxed selection on that allele, as seen in the oribatid mite *Platynothrus peltifer* (Öztoprak et al. 2025). Alternatively, it has been suggested that haplotype divergence may lead to a divergence in function between alleles, potentially with adaptive benefits (Pouchkina-Stantcheva et al. 2007). Whether independent evolution of alleles is a widespread phenomenon among diploid clonal parthenogens remains an open question in the field.

Clades in which asexual reproduction has evolved repeatedly are especially useful for investigating how meiosis has been modified and how the modifications affect inheritance patterns and evolution. Parthenogenesis is particularly common in three phyla of desiccation-tolerant, microscopic animals – rotifers (Welch and Meselson 2000), nematodes (Denver et al. 2011), and tardigrades (Bertolani 2001) – likely because these animals can undergo passive, long-distance dispersal by wind (Bertolani et al. 1990; Incagnone et al. 2015; Ptatscheck et al. 2018; Fontaneto 2019), which makes uniparental reproduction evolutionarily advantageous by allowing a single individual to found a new population (Baker 1955; Cheptou 2012). In tardigrades, many species are thought to exist all around the globe (Bertolani et al. 1990; Gąsiorek 2024), consistent with the theory that individuals have been spread by long-distance passive dispersal. There is also some evidence that parthenogenetic tardigrades occupy larger geographic ranges than sexual relatives (Guidetti et al. 2019). However, despite the growing use of these animals as laboratory models (Goldstein 2022; Arakawa 2022), the cellular and genetic mechanisms underlying parthenogenetic reproduction have not been definitively documented in any species of tardigrade (Blanc et al. 2025).

The tardigrade *Hypsibius exemplaris* is an emerging laboratory model for studies of extremo-tolerance and, thanks in part to its transparent embryos, evolutionary developmental biology (Goldstein 2018a), but its mode of reproduction has not been definitively resolved. Although parthenogenesis is the most common mode of reproduction among tardigrade species studied to date (Bertolani 2001), the mechanism of asexual reproduction in this clade has only been observed cytologically in one species: Prior to the taxonomic separation of *Hypsibius exemplaris* from *Hypsibius dujardini* (Gąsiorek et al. 2018), Ammermann observed tardigrade embryos under a microscope and proposed that parthenogenesis in *H. dujardini* involved an additional round of DNA replication between meiosis I and meiosis II (Ammermann 1967). This mechanism is predicted to result in essentially complete loss of heterozygosity each generation, though Ammerman did not test this expectation. Additionally, no subsequent studies have examined the meiotic mechanism of parthenogenesis in *H. exemplaris*, and its pattern of allele inheritance remains entirely unknown. Clarifying the cytological basis of parthenogenesis and its genetic consequences in this species is therefore essential for understanding how and how long ago asexuality evolved in this tardigrade lineage, as well as for interpreting genetic variation in an increasingly widely used model organism.

Here, we investigate how meiosis was modified for parthenogenesis in *Hypsibius exemplaris* and explore its genomic consequences. First, we cytologically characterize meiosis in this species. We then test whether heterozygosity is stably inherited from generation to generation using locus-specific assays and quantify genome-wide heterozygosity using existing sequencing data from individual tardigrades. Finally, we examine whether long-term maintenance of heterozygosity is associated with the accumulation of allele-specific mutations predicted to disrupt gene function, and we assess the transcriptional activity of genes carrying such variants. Together, these analyses link modified meiosis to patterns of allele inheritance and provide insight into the evolutionary consequences of maintained heterozygosity in asexual reproduction.

## Materials and Methods

### Maintenance of tardigrade cultures

*Hypsibius exemplaris* cultures (Z151) were maintained as previously described (McNuff 2018). Briefly, cultures were kept in 35mm vented petri dishes (Fisher FB0875711YZ) with approximately 2mL of Deer Park spring water and 0.5mL algae (*Chloroccocum sp.* from Carolina Biological Supply). Cultures were kept at room temperature and split approximately once every three weeks. For inheritance tracking, individual tardigrades were maintained in round-bottom 96-well plates (Greiner CELLSTAR 650185) with approximately 100uL of spring water plus 5uL algae per well.

### Fixed DNA staining and imaging of embryos

To precisely track the timing of meiosis, gravid adults were collected and observed until embryos were laid, upon which the time was recorded. Embryos were removed from the cuticle using a 25G needle (BD 305122) and then fixed for imaging as previously described (Smith 2018). Briefly, embryos were added to a 4% paraformaldehyde fixative solution and left for 30 minutes to rock at room temperature, followed by five washes. Embryos were added to the fixative solution in 5-minute intervals post-laying to complete a comprehensive timeline of post-laying meiosis. When multiple embryos were laid by the same mother, pairs of embryos were fixed together at the same time-point, followed by the next pair 10 minutes later (Figure S1e). To image embryos before laying, gravid adults were added to the same fixative solution and embryos were dissected out directly into the solution using a needle, after which the same protocol was followed. After fixation and washes, the eggshells of embryos were nicked with a needle to increase permeability. Embryos were then mounted onto slides with 28.41µm glass beads (Whitehouse Scientific MS0028) and approximately 2uL of DAPI fluoromount-G mounting media (Southern Biotech 0100-20). Slides were sealed with clear nail polish after the mounting media was allowed to dry overnight. Imaging was performed using a Zeiss 710 laser scanning confocal microscope with a 100x objective.

### Live DIC imaging of embryos

Embryos were imaged by Differential Interference Contrast (DIC) light microscopy as previously described (Heikes and Goldstein 2018). Briefly, gravid adults were collected and mounted onto slides with 28.41µm glass beads (Whitehouse Scientific’s Monodisperse Standards) and Deer park spring water. Slides were sealed immediately with Valap, a 1:1:1 mixture of Vaseline, lanolin, and paraffin maintained at 70°C and applied with a paintbrush. Images were taken in 5-minute intervals for timelapses using a E800 upright microscope with a 60x oil objective and pco.panda sCMOS or Nikon DS-Fi3 camera.

### DNA isolation from single tardigrades or pooled samples

DNA was isolated from tardigrades through modification of a single-worm lysis protocol developed for *Caenorhabditis elegans* (Williams et al. 1992). Single tardigrades were added to 5uL worm lysis buffer with Proteinase K, or 15-20 animals were added to 15uL of the same buffer, and then tubes were centrifuged briefly. Samples were twice frozen at −80°C and then thawed, the first time at room temperature and the second in a thermocycler block preheated to 65°C. Samples were then denatured at 65°C for 1 hour followed by 80°C for 15 minutes. Before use in PCR, water was added to double the volume of the initial buffer solution and samples were vortexed repeatedly. DNA samples were stored at −20°C. We were able to perform approximately 10-12 PCR reactions per individual using this method.

### Selection of loci for genotyping individual tardigrades

Conserved single-copy genes were identified in *Hypsibius exemplaris* for tracking the inheritance of alleles across generations. From the OrthoDB catalog of orthologs (Kuznetsov et al. 2023), we generated a list of genes that were present in >90% and single-copy in >90% of all metazoans in the database. We output the protein sequences for these genes for the arthropod *Drosophila melanogaster*, randomly selected 10 genes from the list using a custom python script, and then searched for homologs of these 10 genes in *Hypsibius exemplaris* using reciprocal BLASTp. We identified homologs for 9/10 genes using this method, attempted to amplify each locus, and successfully amplified three loci by nested PCR (see Table S1 and Figure S2). These three successful loci were used for subsequent experiments. Similarly, for genotyping individuals at loci near the ends of chromosomes, we attempted to amplify loci that were 1) near the ends of chromosomes 1 and 2 according to a putative chromosome-level assembly generated using Hi-C data (Dudchenko et al. 2017; Dudchenko et al. 2018; Hoencamp et al. 2021; DNA Zoo Consortium dnazoo.org)) and 2) on the same NCBI reference assembly scaffolds as predicted telomeric sequences (Yoshida et al. 2017). We were successful in amplifying two loci (see Table S1 and Figure S4) which were then used for subsequent experiments.

### Primer design and PCR

Primers were designed using NCBI Primer BLAST optimized for PCR with an annealing temperature of 60°C. For most loci, nested PCR was performed in two steps whereby the first step amplified a region that encompassed the entire final target sequence for 20 cycles, followed by amplification of the final target sequence for 30 cycles. PCR reactions utilized either the Q5 High-Fidelity 2X Master Mix (NEB M0492) or the LongAmp Taq 2x Master Mix (NEB M0287) with reaction and cycle parameters determined by the manufacturer’s protocol. For downstream cloning and allele sequencing, when applicable, the second pair of primers included a 20bp overhang for cloning into a pDEST17 destination vector. This overhang was omitted for genotyping assays. Primers are listed in Table S1.

### Allele-specific amplicon sequencing

Final products from PCR performed on pooled tardigrade samples (15-20 animals each) were purified using the Zymoclean Gel DNA Recovery Kit (Zymo D4008) and quantified using Nanodrop. For conserved single-copy gene loci, amplicons were cloned into a pDEST17 Gateway destination vector (Invitrogen 11803012) using the NEBuilder HiFi DNA Assembly Master Mix kit (NEB E2621), and plasmids were transformed into chemically-competent NEB 5-alpha *E. coli* (NEB C2987), both according to the manufacturer’s protocols. The resulting plasmids each contained a single allele from the heterozygous mixture of PCR products. Multiple plasmids per locus were sequenced via Whole Plasmid Sequencing performed by Plasmidsaurus using Oxford Nanopore Technology with custom analysis and annotation. For loci near telomeric sequences, mixed-allele PCR products were directly sequenced via Premium PCR Sequencing by Plasmidsaurus using Oxford Nanopore Technology with custom analysis and annotation. Data was analyzed using custom scripts in Python version 3.14.5 (Python 3.14 Documentation 2001) and R version 4.4.0 (R Core Team 2025).

### Genotyping

Sequencing data for both alleles of each locus was analyzed using Benchling to count heterozygous sites and identify restriction enzymes that differentially cut the two alleles. Enzymes used for each locus are listed in Table S1. Final products from nested PCR performed on single tardigrades (or pooled samples for testing purposes) were directly digested with the corresponding enzyme. Resulting products were analyzed using gel electrophoresis (1% agarose with Ethidium Bromide). A second technical replicate of genotyping was attempted for cases when one or both alleles was not detected in the original experiment, with some cases not undergoing technical replication due to limited DNA per individual.

### Variant calling, heterozygosity calculations, and variant impact annotations

Reads from the genomic sequencing datasets described in Table S2 were mapped against a reference genome using the BWA-MEM algorithm from the Burroughs-Wheeler Alignment Tool version 0.7.17 (Li 2013). Variants were called and filtered using BCFtools version 1.22 (Danecek et al. 2021). Sites were filtered for depth (minimum 40 reads) and quality (minimum 30 quality score). The choice of quality score cut-off did not impact the pattern of heterozygosity across chromosomes, and implementing this filter excludes highly repetitive regions from analysis. Post-filtering, sites were called as heterozygous if a minor allele was present at a frequency above 0.25, keeping the expected false-positive rate below 0.0022 (the binomial probability of sampling the minor allele in fewer than 10 of 40 reads). This method biases against false positive heterozygosity calls. The same method of heterozygosity calling was used for both individual (Arakawa et al. 2016) and bulk (Boothby et al. 2015) tardigrade sequencing datasets. Heterozygosity was calculated by mapping against two reference genomes: 1) the reference assembly for *Hypsibius exemplaris* currently stored in NCBI (Yoshida et al. 2017), which is annotated but is not assembled to a chromosome level, and 2) a putative chromosome-level assembly generated using Hi-C data along with many other species (Dudchenko et al. 2017; Dudchenko et al. 2018; Hoencamp et al. 2021; DNA Zoo Consortium dnazoo.org)). We used the NCBI reference genome for calculating genome-wide levels of heterozygosity, as well as for assessing the predicted functional consequences of heterozygous variants, because it is the more complete genome and is annotated. We only used the Hi-C genome for analyzing the pattern of heterozygosity across putative chromosomes. In this case, the percentage of heterozygous sites per 100kb bin was calculated as the percentage of sites post-filtering that fit the described metric for heterozygosity. To assess the predicted functional consequences of variants, we made a list of heterozygous variants that were detected in all three bulk tardigrade sequencing replicates (Boothby et al. 2015) and at least one individual tardigrade sequencing replicate (Arakawa et al. 2016), all mapped against the annotated NCBI reference assembly (Yoshida et al. 2017). These heterozygous variants were analyzed for their predicted functional impact on genes using SnpEff version 5.2f (Cingolani et al. 2012). Data was compiled and analyzed using custom scripts in R version 4.4.0 (R Core Team 2025). OpenAI ChatGPT-4 and ChatGPT-5 LLMs were used for code syntax and troubleshooting. All sequencing datasets and reference genomes used in this study are listed in Table S2.

### mRNA expression analysis

Relative expression levels were quantified from three replicate RNA-seq datasets of approximately 10k active, adult tardigrades each (Yoshida et al. 2017) mapped against the NCBI reference genome (Yoshida et al. 2017) using Salmon version 1.10.1 (Patro et al. 2017) and tximport version 1.3.9 (Soneson et al. 2016). Allele-specific expression was calculated using ASEQ (Romanel et al. 2015) on heterozygous sites, requiring minimum depth coverage of 10, minimum base quality of 10, and minimum read quality of 20. For each heterozygous site that met these quality filters and occurred within a transcribed gene, we obtained from the ASEQ output the number of reads in each replicate that mapped to the variant allele (i.e., the ALT allele) and to the REF allele. We assessed the known bias that favors alignment of reads containing reference over non-reference (ALT) alleles by calculating the mean frequency of the ALT alleles across all heterozygous sites and samples. The resulting mean of 0.471 was used as the null expectation for ALT allele frequency under equal expression of the reference and ALT alleles. For heterozygous sites that met the ASEQ quality filters in all 3 replicates, we determined the binomial probability (p-value) of obtaining an ALT allele frequency as low or lower than the observed allele frequency, given the total number of mapped reads and a true variant allele frequency of 0.471. Variants were considered to exhibit allele-specific reductions in expression if the ALT allele was the minor allele in all three samples and if the median p-value of that allele across the three samples was less than 5.78 × 10^−6^ (0.05/595; the sequential Bonferroni-corrected threshold for multiple tests in this case of 607 out of 9258 tests achieving significance). Data was compiled and analyzed using custom scripts in R version 4.4.0 (R Core Team 2025). OpenAI ChatGPT-4 and ChatGPT-5 LLMs were used for code syntax and troubleshooting. All sequencing datasets and reference genomes used in this study are listed in Table S2.

## Results

### *Hypsibius exemplaris* reproduces asexually by abortive meiosis I

To investigate how meiosis is modified for asexual reproduction in this species, we first asked how chromosomes are partitioned during meiotic divisions. Many attempts to stain DNA in living embryos without also disrupting development were unsuccessful (Papell et al. 2026). Therefore, we imaged embryos fixed at finely-timed stages immediately after laying to generate a timeline of meiosis. *Hypsibius exemplaris* is diploid with 10 chromosomes (Gabriel et al. 2007), so if meiosis were occurring canonically, we would expect two divisions of 5 pairs: First, a division of homologous chromosomes, followed by a division of sister chromatids. Analysis of our timeline of meiosis revealed a meiosis-like anaphase of 5 pairs of chromosomes followed by a second anaphase of 10 pairs of chromosomes (Figure 1a). In metaphase and anaphase of both divisions, chromosomes were arranged in rings and roughly equally spaced (Figure 1a and Figure S1). Interestingly, embryos appeared to be paused in anaphase I for a significant duration of meiosis, during which the configuration of chromosomes remained intact (Figure S1b). Later, a single polar body was observed following the second anaphase (Figure 1a). From this result alone, there are two possibilities for meiosis modification that could produce this pattern. The first possibility is abortive meiosis I, where meiosis I metaphase and anaphase occur but the division does not progress fully to cytokinesis producing a polar body. The second possibility is that even though we failed to detect a polar body following meiosis I with this method, a polar body is formed following each division and an additional replication of DNA occurs between meiosis I and II (as proposed by (Ammermann 1967)).

**Figure 1:**
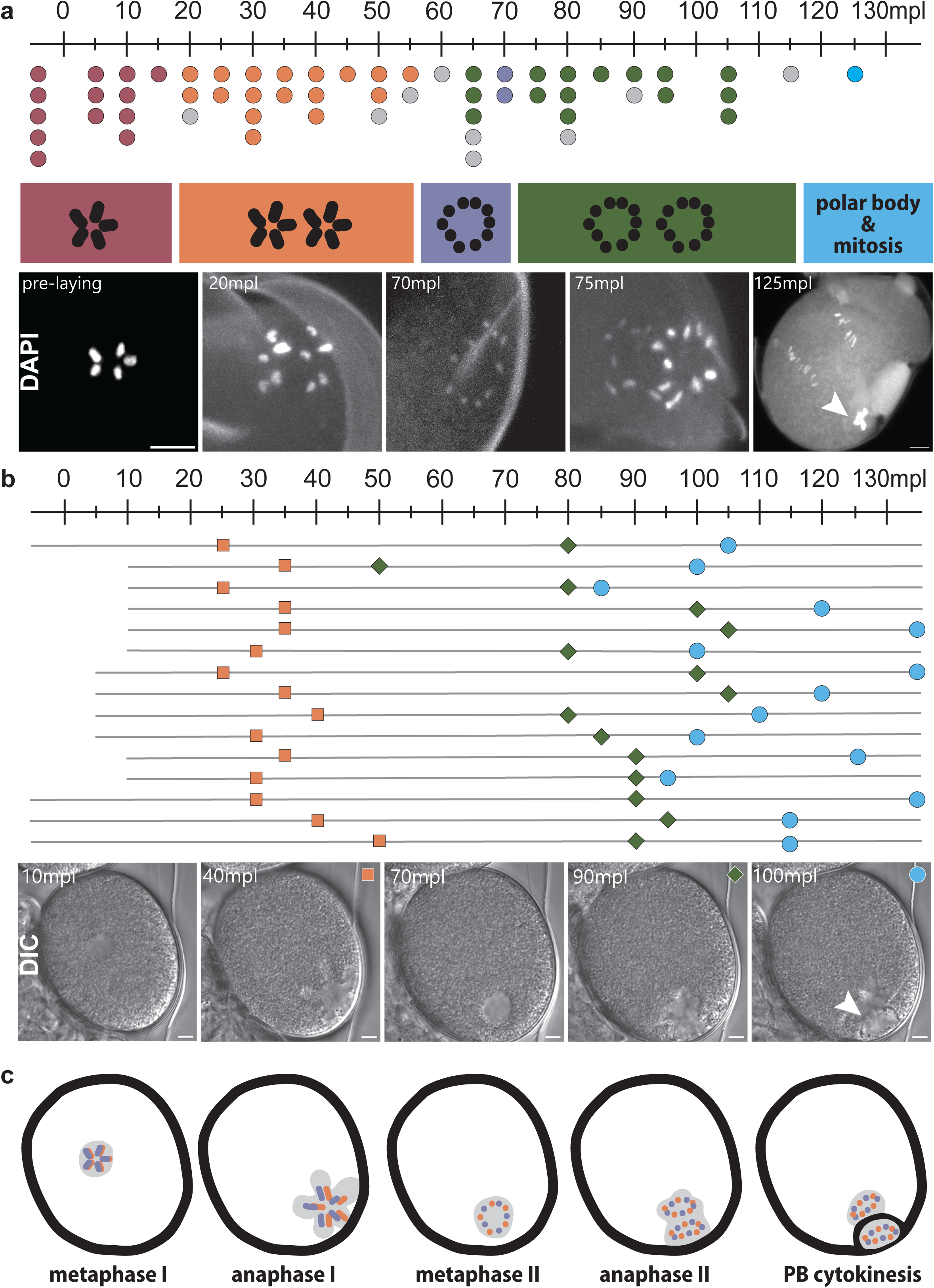
The tardigrade *Hypsibius exemplaris* reproduces asexually by abortive meiosis I. **(a)** Each dot represents a tardigrade embryo that was fixed along a timeline from 0 to 125 minutes post-laying (mpl). A total of 57 embryos laid by 22 adults were fixed in pairs and imaged (see Figure S1e for relationships between embryos). Dots to the left of zero on the timeline represent those that were cut out of gravid adults directly into fixative solution. Embryos were then stained with DAPI and imaged using a scanning confocal microscope. During analysis, embryos were categorized into one of five groups depending on the number and orientation of visible chromosomes (red: five chromosomes arranged in a ring, orange: five pairs of chromosomes separating, purple: ten chromosomes arranged in a ring, green: ten pairs of chromosomes separating, blue: visible polar body & mitotic division). Uncategorized embryos are shown in gray. Representative images of DAPI staining within embryos, one for each category, are shown. The arrowhead points to a polar body. Scale bars are 5µm. **(b)** Live embryos were imaged every 5min with Differential Interference Contrast (DIC), and the relative timing of when they were laid was recorded. Each line indicates the timeline of a single embryo progressing through meiosis, starting from the time it was laid. Only embryos where a polar body was ultimately visible were included, for a total of 15 embryos laid by 11 adults (maximum of two embryos per adult). Orange squares indicate the first time DNA migrated to the periphery, and green diamonds indicate the second time DNA migrated to the periphery. Blue circles indicate the first time a polar body was visible (characterized by a membrane surrounding DNA at the periphery). Finally, images of a single representative embryo are shown over time as it proceeds through meiosis, with symbols marking the events corresponding to the above timelines. The arrowhead points to a polar body. Scale bars are 5µm. **(c)** Model of modified meiosis cytology in *H. exemplaris*. Inferred homologous chromosomes are differentially colored, purple and orange. Light gray surrounding the chromosomes represents the area visible by DIC imaging. Meiosis I metaphase and anaphase occur but do not conclude with polar body cytokinesis. Meiosis II then proceeds as normal, resulting in a single polar body. Alt text: Timelines ranging from 0 to 130 minutes post-laying, symbols depicting replicates of imaged *H. exemplaris* embryos, and representative images for different phases of modified meiosis are shown for each of two microscopy experiments (panels a and b).

To distinguish between these hypotheses, we turned to live imaging by Differential Interference Contrast (DIC) to observe meiosis and polar body formation (Video S1). Immediately upon laying, embryos appeared to have a rounded nucleus (Figure S1a-b). The chromosomes then migrated to the cell periphery, where a shape reminiscent of a five-pointed star became visible (Figure S1b). Combined with our fixed-embryo results, this shape corresponds to the five pairs of chromosomes held in configuration during anaphase I. The star shape migrated along the cell periphery before moving slightly inward and forming a second rounded shape (Figure S1b-c). Finally, the chromosomes returned to the cell periphery and a division with cytokinesis occurred, producing a polar body (Figure 1b and Figure S1d). To robustly test whether polar body formation only ever follows the second anaphase, we generated a second timeline from DIC films. In 100% of 15 filmed embryos, we observed a single polar body, which was only visible following the second trip of chromosomes to the cell periphery (Figure 1b). Taken together with the previous result, we conclude that *H. exemplaris* reproduces via abortive meiosis I in which metaphase and anaphase of meiosis I occur but do not result in the formation of a polar body, and the chromosomes that are canonically released into the first polar body are instead retained (Figure 1c).

### Heterozygosity is inherited from mother to daughter

All mechanisms of meiotic parthenogenesis cause non-Mendelian allele inheritance patterns, but the resulting inheritance pattern varies depending on how meiosis is modified. With abortive meiosis I, whether and to what extent heterozygosity is lost depends on two characteristics of canonical meiosis that cannot be assessed by microscopy: independent assortment of chromatids during meiosis II and crossing over. If these characteristics of canonical meiosis are maintained in *H. exemplaris*, reproduction should result in loss of heterozygosity distal to crossovers (Figure 2). We set out to investigate whether heterozygosity is lost in *H. exemplaris* by tracking the inheritance of alleles from parents to offspring.

**Figure 2:**
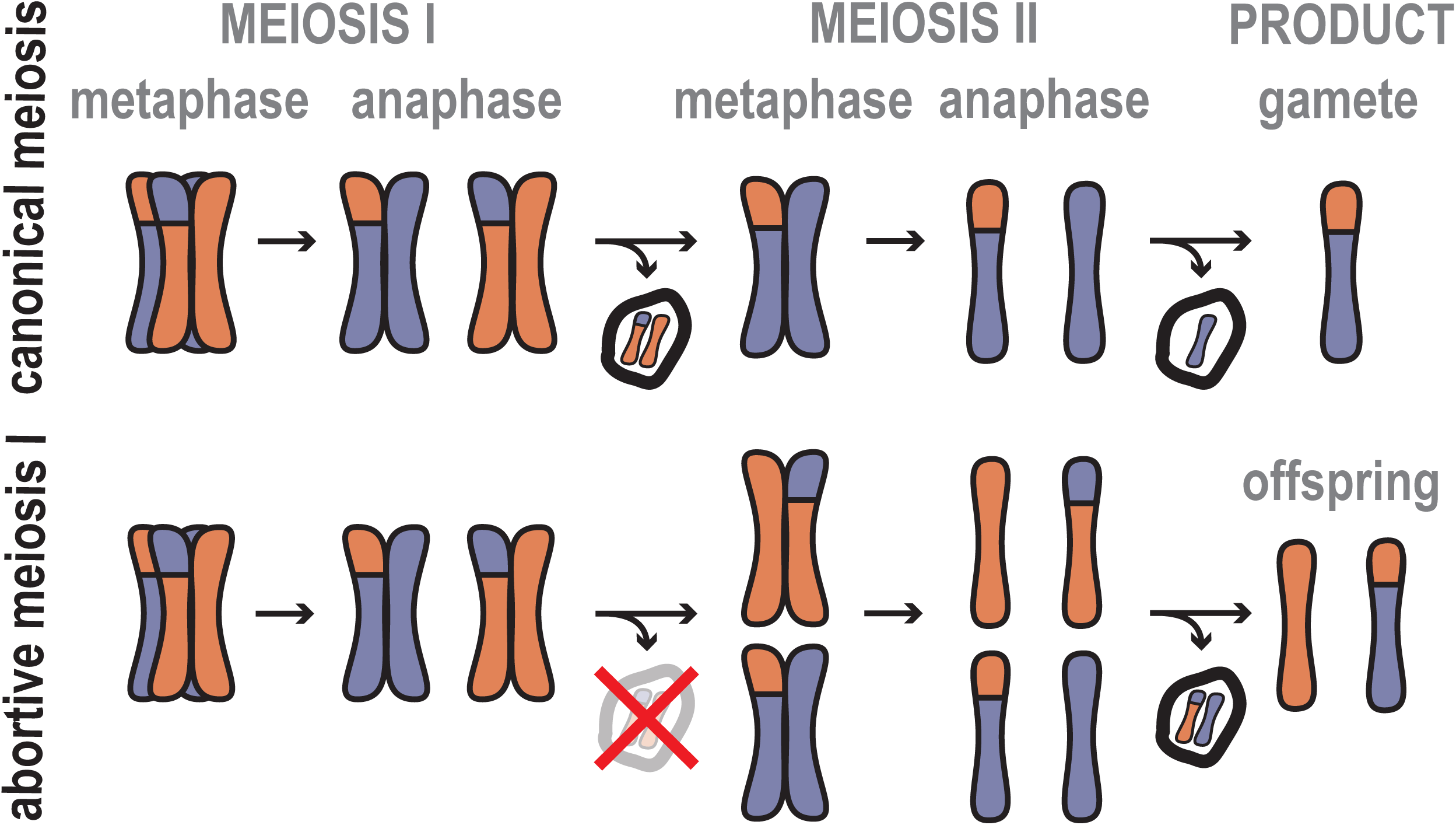
Predicted genetic consequences of abortive meiosis I parthenogenesis with crossing over. One pair of homologous chromosomes, purple and orange, is shown proceeding through canonical and modified meiotic divisions. A single crossover has occurred between a pair of non-sister chromatids. Meiosis I is a division of homologous chromosomes, canonically producing a polar body. Meiosis II is a division of sister chromatids, canonically producing a second polar body. In the case of abortive meiosis I, the first polar body is not formed, and an assortment of non-sister chromatids in meiosis II results in a polar body and a diploid offspring. Assuming a single crossover, independent assortment during meiosis II makes it so the probability of loss of heterozygosity distal to the crossover site is 50%. Alt text: Labeled diagram showing one pair of homologous chromosomes proceeding through either canonical meiosis or modified meiosis.

First, we set out to identify loci at which individuals are heterozygous, assuming any exist. To avoid misinterpreting duplicated genes as heterozygous alleles, we identified *H. exemplaris* homologs of genes that are largely conserved as single-copy across metazoans (Kuznetsov et al. 2023) (see Methods). We used nested PCR on pooled samples of 15-20 tardigrades to amplify the entire coding regions of three genes from this list, each located on a different chromosome of a putative chromosome-level assembly (Dudchenko et al. 2017; Dudchenko et al. 2018; Hoencamp et al. 2021; DNA Zoo Consortium dnazoo.org)) (Figure S2a and Table S1). To reliably sequence all potential alleles at these loci, we cloned PCR products into plasmids and sequenced at least 6 plasmids per gene. We found that all three gene loci, including flanking regions, had two major alleles. Differences between the two alleles made up 1.42%, 1.11%, and 3.22% of total sequenced nucleotides for the three loci (Figure S2b). To genotype individual tardigrades, we used these allele sequences to identify restriction enzymes that selectively cut one of the two alleles at each locus. Applying this genotyping assay to an individual tardigrade, we found that the individual contained both alleles, and thus was heterozygous, at all three loci (Figure 3a).

**Figure 3:**
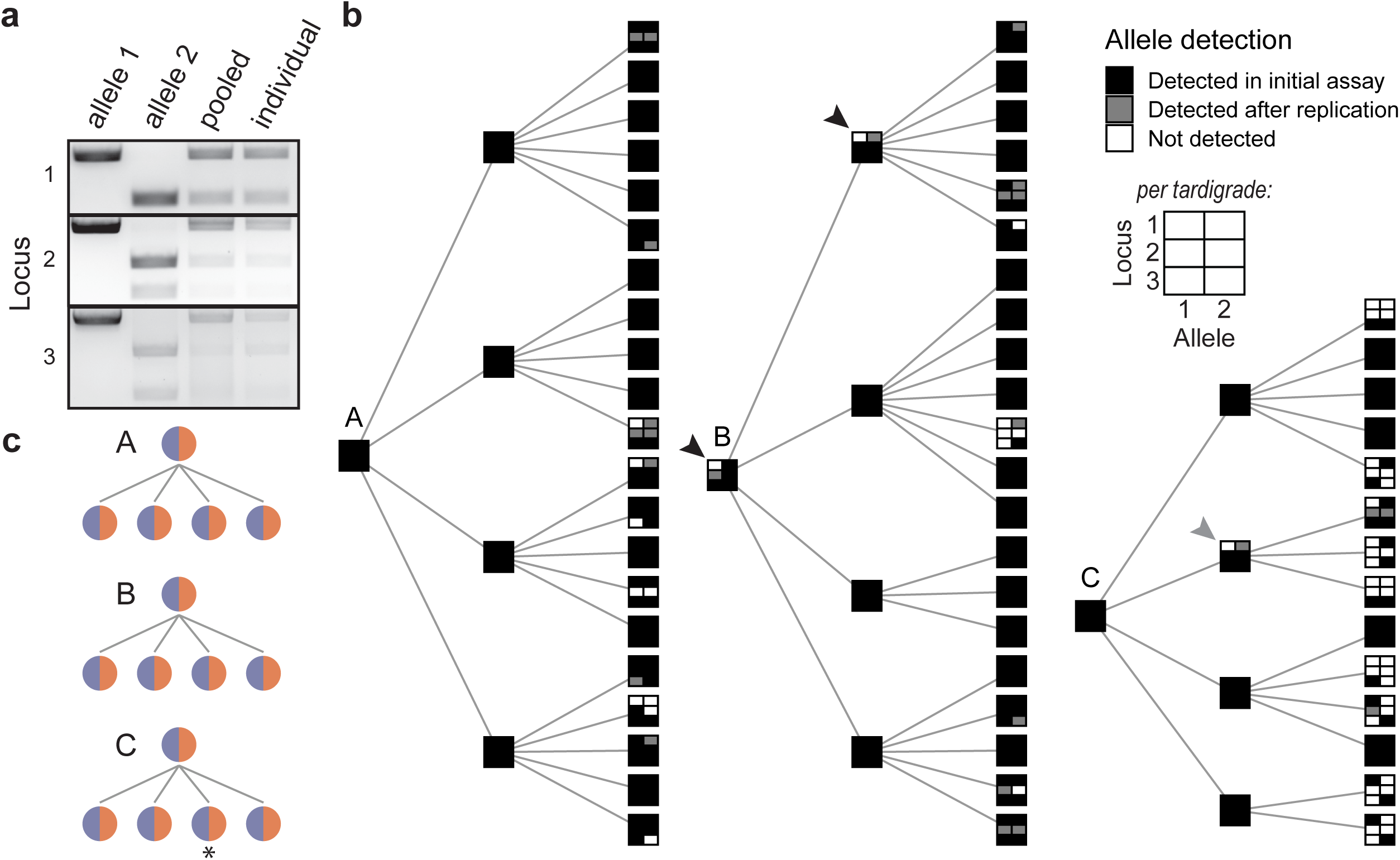
Heterozygosity is inherited through the germline at three independent loci. **(a)** PCR followed by allele-specific restriction enzyme digestion was used to detect alleles at three loci. PCR products obtained from plasmids containing each allele and pooled tardigrade samples were included as controls. Allele detection in an individual tardigrade reveals that this individual is heterozygous at all three loci. **(b)** Allele inheritance was tracked in three asexual pedigrees over three generations. Each individual is represented as a square divided into six quadrants, with rows corresponding to the three loci and columns corresponding to the two alleles at each locus. Quadrants are filled when the corresponding allele was detected by the restriction-enzyme assay. In a subset of individuals, alleles not detected in the initial assay were detected in a second technical replicate (gray). Black arrowheads point to individuals for which detection of both alleles in their offspring implies parental heterozygosity despite incomplete allele detection, and a gray arrowhead points to the single instance of first and second generation individuals where heterozygosity was not able to be confirmed from allele detection assays in the parent or its offspring (locus 1). **(c)** Inferred germline genotype pedigrees for all tardigrade families based on allele detection results in each parent and/or their offspring. Only one locus is shown because the results for all three loci are the same, except for the single starred individual, in which heterozygosity could not be confirmed for locus 1. Alt text: A labeled image of gel electrophoresis for multiple samples (a) and diagrams showing pedigrees of allele inheritance, where multiple individuals in a family are shown with lines connecting parent to offspring. Pedigrees in panel b show three generations, and pedigrees in panel c show two.

To investigate the rate of loss of heterozygosity in *H. exemplaris*, we set out to track the inheritance of alleles from generation to generation. To accomplish this, we established three tardigrade families by raising mothers and daughters in individual wells of 96-well plates for three successive generations. We then genotyped all tardigrades in these families using restriction enzyme genotyping assays and found that both alleles were detected in most individuals for all three loci (Figure 3b; for each locus, we detected both alleles in 54, 59, and 62 individuals out of 71 tested). In a subset of individuals, a second technical replicate detected alleles that were not observed in the initial assay (Figure 3b, gray boxes), indicating that allele non-detection generally reflected technical limitations rather than true allele loss. In further support of this conclusion, in rare cases where we detected only one allele in the first or second generation, offspring were often found to carry both alleles (Figure 3b, black arrows). In only one instance, inheritance of both alleles could not be confirmed by either technical replication or offspring genotypes (Figure 3b, gray arrow). These results may reveal a very low rate of loss of heterozygosity: a maximum of one case out of 45 that we examined (15 first or second generation individuals times 3 loci). Otherwise, heterozygosity was stably inherited through the germline between generations (Figure 3c).

### *Hypsibius exemplaris* is heterozygous across its genome

Our previous results revealed that many individuals are heterozygous at three specific loci, and that heterozygosity is generally inherited, but we cannot distinguish whether loss of heterozygosity due to crossovers is rare or absent entirely. Additionally, it remains possible that the three loci we studied harbored rare examples of heterozygous sites. Over sufficient evolutionary time with abortive meiosis I parthenogenesis causing loss of heterozygosity, we would expect *H. exemplaris* to have very low levels of heterozygosity across its genome. Therefore, we next characterized heterozygosity of *H. exemplaris* individuals at a genome scale. We calculated the percentage of heterozygous sites across the entire genome for four individual tardigrades (see Methods; sequencing data generated by (Arakawa et al. 2016); see Table S2 for details of all published datasets used in this study). We performed this calculation by mapping reads against the current scaffold-level NCBI reference assembly (generated by (Yoshida et al. 2017); see Table S2). Across the four samples, the mean percentage of heterozygous bases genome-wide was 1.40% +/− 0.17% standard error (as expected, heterozygosity was elevated at four-fold degenerate sites: 1.82% +/− 0.22% standard error; Figure S3a). For context, 1-2% heterozygosity is consistent with several clonally-reproducing parthenogenetic animals (Brandt et al. 2021; Blanc et al. 2023; Lacy et al. 2024).

Next, we set out to investigate the pattern of heterozygosity across chromosomes in *H. exemplaris*. Because abortive meiosis I parthenogenesis predicts crossover-dependent loss of heterozygosity (Figure 2), we reasoned that even with overall retained heterozygosity, rare crossovers may leave genomic signatures visible on a chromosome-wide scale. For example, for monocentric species that reproduce by abortive meiosis I, loss of heterozygosity is expected distal to crossover sites (as shown in Figure 2). For these species, even very occasional crossover events over evolutionary time may result in the general pattern of lower heterozygosity near the ends of chromosomes compared to the middle. Chromosomes in *H. exemplaris* appear to be pulled by a single point during early mitotic divisions (Figure 1a, rightmost DAPI panel), making this species likely monocentric. If there are defined centromeres in mitosis, it is likely these are also used in meiosis, but regardless of centromere type, we would expect large stretches of homozygosity from recent but rare crossover events, as seen in the asexual rotifer *A. vaga* (Simion et al. 2021) and asexual nematode *Halicephalobus mephisto* (Amini and Bracht 2025). To assess the pattern of heterozygosity across the genome, we used a putative chromosome-level assembly generated using Hi-C by the DNA Zoo Consortium (Dudchenko et al. 2017; Dudchenko et al. 2018; Hoencamp et al. 2021; DNA Zoo Consortium dnazoo.org)) (Table S2) because the NCBI reference genome is not assembled to the chromosome level. As a preliminary validation of this assembly, we used BLAST to align predicted *H. exemplaris* telomeric repeats (Yoshida et al. 2017) against the putative chromosomes and found that in at least three out of five chromosomes, telomeres appeared to be highly enriched at both ends (Figure 4a). Additionally, genome-wide heterozygosity calculated using the Hi-C assembly as a reference genome was similar to what we found using the NCBI reference assembly (Figure S3a). When we plotted the pattern of heterozygosity within individuals across the length of each chromosome, we found that heterozygosity was fairly evenly distributed without any distinctive regions of loss of heterozygosity (Figure 4b). This observation was true for each of the four single-tardigrade replicates (Figure S3b), indicating that differences in heterozygosity levels between these samples were not due to recent crossover events. Because this chromosome-level assembly has not been rigorously verified, we also tested whether heterozygosity has been maintained very near chromosome ends by genotyping individuals at two loci on telomere-containing NCBI scaffolds (Yoshida et al. 2017) (Table S1). We found that 3 out of 3 tested individual tardigrades were heterozygous at both loci (Figure S4), indicating that individuals are heterozygous very near the ends of chromosomes. We conclude that *H. exemplaris* individuals show no genomic signs of loss of heterozygosity due to meiotic crossovers.

**Figure 4:**
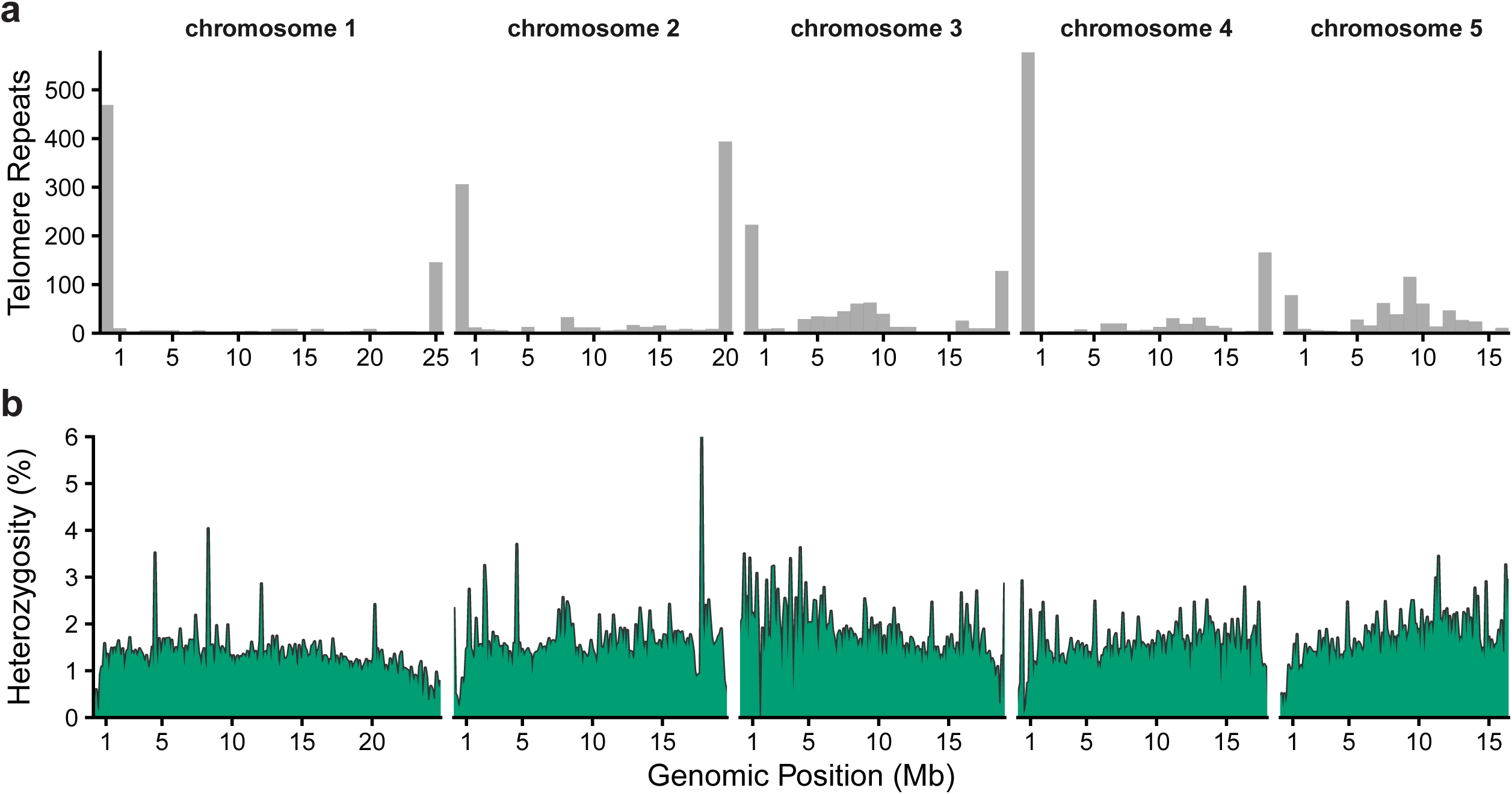
Heterozygosity is relatively stable across the genome. **(a)** Using BLAST+, the predicted *Hypsibius exemplaris* telomeric repeat sequence (GATGGGTTTT) from Yoshida et al. 2017 was aligned to the putative five-chromosome genome assembly generated using Hi-C data by the DNA Zoo Consortium (see Table S2). Shown are the total counts of alignments of this sequence per 1Mb bin. **(b)** Heterozygosity across the five chromosomes of *Hypsibius exemplaris* was calculated from single-tardigrade sequencing data generated by Arakawa et al. 2016, and using the chromosome-level genome assembly above. Heterozygosity was calculated in 100kb bins as the percentage of heterozygous bases out of all sites that passed filtering cut-offs. Heterozygosity was calculated for each of the four individuals sequenced, and the mean across the four replicates (per bin) is shown here. Alt text: Plots showing how alignment of telomeric repeats (a) and heterozygosity (b) varies across the length of five chromosomes.

### Maintained heterozygosity may permit variants that render one allele non-functional

It is not known how long ago parthenogenetic *H. exemplaris* diverged from sexual ancestors, but maintenance of heterozygosity is expected to permit divergence between alleles over time (Welch and Meselson 2000). To explore whether such divergence is detectable in *H. exemplaris*, we examined evidence for genes with highly divergent alleles, focusing on cases where one allele has accumulated mutations predicted to disrupt gene function. Specifically, we sought genes carrying heterozygous variants consistent with one allele of an otherwise transcriptionally expressed gene harboring predicted loss-of-function mutations.

We identified heterozygous variants and annotated them based on their predicted effects on protein function. First, we made a list of heterozygous variants that were detected in three bulk tardigrade sequencing replicates from (Boothby et al. 2015) and at least one individual tardigrade sequencing replicate from (Arakawa et al. 2016) (Table S2). This conservative filtering was used to increase confidence that the variants analyzed are heterozygous within individuals, as well as recurrent within laboratory populations rather than arising from technical noise or rare individuals. Next, we annotated these variants by their predicted functional consequences (see Methods) and focused on genes containing variants classified as high impact (Figure 5a). Several genes carried stop-gained variants that truncate ≥25% of the predicted protein, while several others contained frameshift variants that alter ≥25% of the protein sequence, both of which likely impact function (Figure 5b-c). Full details for all genes with predicted high-impact variants are provided in Table S3.

**Figure 5:**
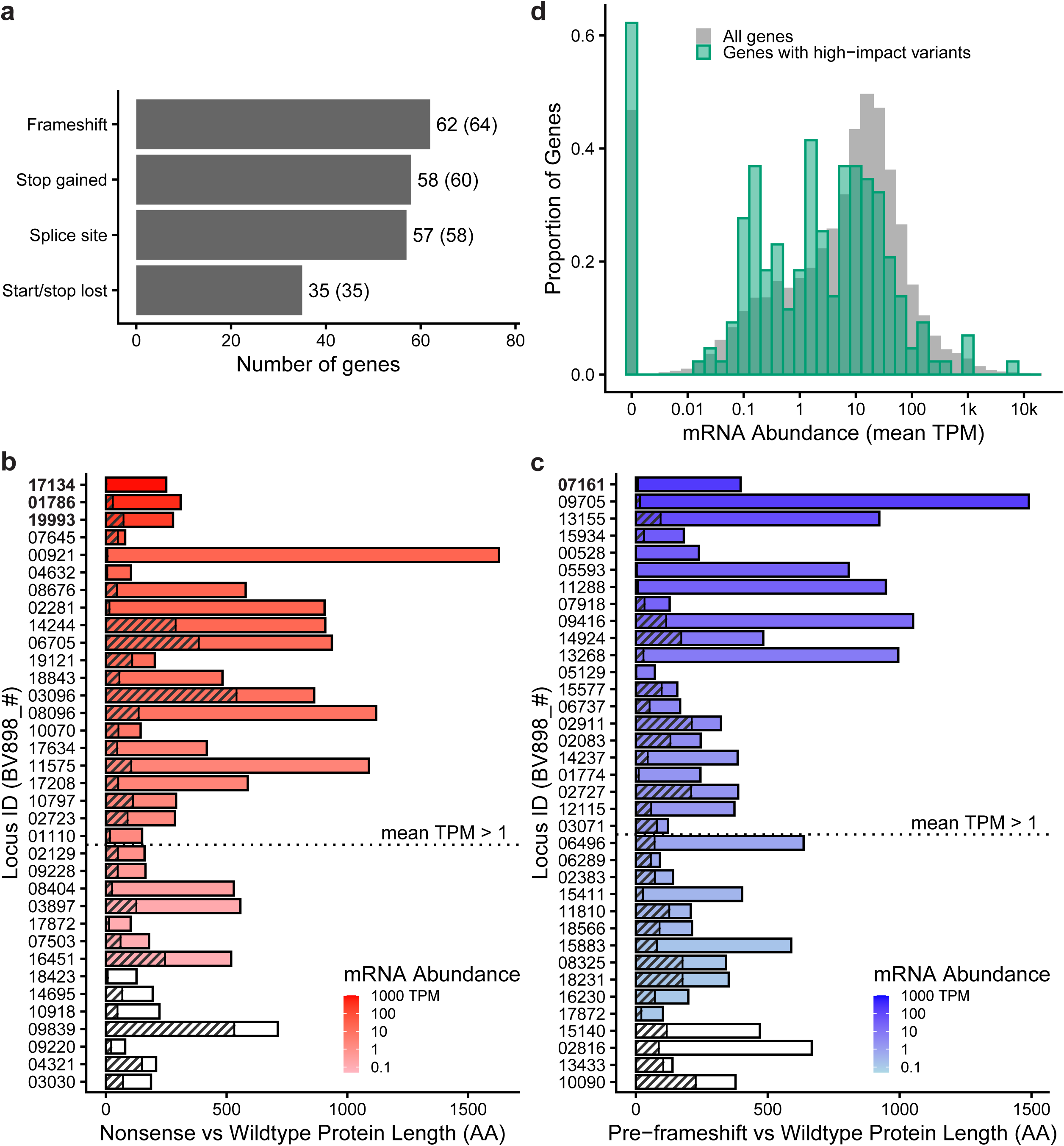
Evidence of variants rendering one allele non-functional in *Hypsibius exemplaris*. **(a)** Genes carrying predicted high-impact heterozygous variants in *H. exemplaris*. Only variants detected in all three bulk tardigrade sequencing datasets and at least one individual tardigrade sequencing dataset are shown. Bars indicate the number of genes per variant consequence class. Numbers in parentheses indicate the total number of variants in each class, since in a few cases a single gene contains two high-impact variants. **(b-c)** Genes with nonsense or frameshift variants that affect ≥25% of the predicted protein sequence. Red, blue, or white bars indicate the predicted functional protein length, while overlaid hashed bars indicate the proportion of that predicted protein sequence that comes before the early STOP or frameshift mutation. Importantly, frameshift variants are likely to impact total protein length by producing downstream stop codons, so the hashed bars do not represent the predicted length of the variant allele protein product. Genes are colored and ordered by mean mRNA transcript abundance (TPM), with white bars representing genes with no detected expression. Bolded Gene IDs are examples with especially high mRNA expression mentioned in the text. **(d)** Distribution of mRNA transcript abundance levels for genes with high-impact heterozygous variants compared to all genes in the genome. Transcript abundance levels represent the total transcript abundance of each gene (both alleles). Histograms show the fraction of genes at each expression level following log10 transformation of mean transcript abundance (TPM) across three RNA-seq replicates from Yoshida et al. 2017. Alt text: Graphs showing gene and variant counts in different categories (a), predicted protein product lengths for select genes (b-c), and a histogram of gene expression levels (d).

The presence of high-impact variants in one allele raises the question of whether these genes remain transcriptionally expressed or if, instead, they represent fully non-functional and non-expressed pseudogenes. To address this question, we examined mRNA expression levels of genes containing high-impact heterozygous variants using previously published RNA-seq datasets (Yoshida et al. 2017) (Table S2). Although genes with high-impact variants were modestly enriched for low or undetectable expression, the distribution of gene expression levels overlapped substantially with that of all annotated genes in the genome (Figure 5d). Notably, several genes with substantial stop-gained or frameshift variants affecting ≥25% of the predicted protein sequence remained highly expressed (Figure 5b-c). For example, the ribosomal subunit protein uL6 (BV898_17134) carries a premature stop codon truncating one allele by 99.2%; an endoplasmic reticulum transmembrane protein (BV898_01786) harbors a stop variant truncating one allele by 90.6%; and the lipid droplet-binding protein 1 gene (BV898_07161) carries a frameshift variant affecting 98% of one allele (all bolded in Figure 5b-c). All three of these genes fall into the top 5% of genome-wide mRNA expression levels (Table S3). These examples illustrate that even genes with severe predicted loss-of-function variants in one allele can remain robustly expressed.

Next, we asked whether the predicted non-functional allele of the genes harboring high-impact variants is less transcribed than the functional allele. The RNA-seq data were of sufficient depth and quality to enable an assessment of allele-specific expression at 48 of the high-impact variant positions (Table S4; see Methods). The predicted non-functional alleles were significantly less transcribed than the reference alleles in 6.25% (3/48) of cases, including 2 intron variants (BV898_07915 and BV898_09527) and the 3^rd^ highest expressed among the nonsense variants (BV898_19993; bolded in Figure 5b). However, that percentage is nearly identical to the 6.59% (593/9001) of heterozygous sites in other genes that also showed significantly less transcription of the non-reference allele. Taken together, the RNA-seq data provide no conclusive evidence that reduced transcription either enabled or followed the predicted functional divergence that resulted from these high-impact variants.

## Discussion

In this study, we found that the tardigrade *Hypsibius exemplaris* reproduces by abortive meiosis I parthenogenesis and maintains heterozygosity across its genome from generation to generation (Figure 6). Additionally, we identified multiple transcriptionally expressed genes that harbor predicted loss-of-function mutations in one allele, suggesting that maintained heterozygosity resulted in divergence between alleles (Figure 6). This work represents the first combined cytological and genetic characterization of parthenogenesis in a tardigrade, situating *H. exemplaris* within a broader but largely underexplored diversity of reproductive mechanisms found among asexual animals. As an increasing number of researchers utilize tardigrades in their laboratories (Goldstein 2022; Arakawa 2022), a clear understanding of allele inheritance patterns in this and other species is increasingly important for the development of genetic tools. Our finding that *H. exemplaris* maintains stable heterozygosity will affect interpretation of, and the approaches needed to generate, genetically modified lineages in this species. Additionally, this finding will affect interpretations of population-level heterozygosity.

**Figure 6:**
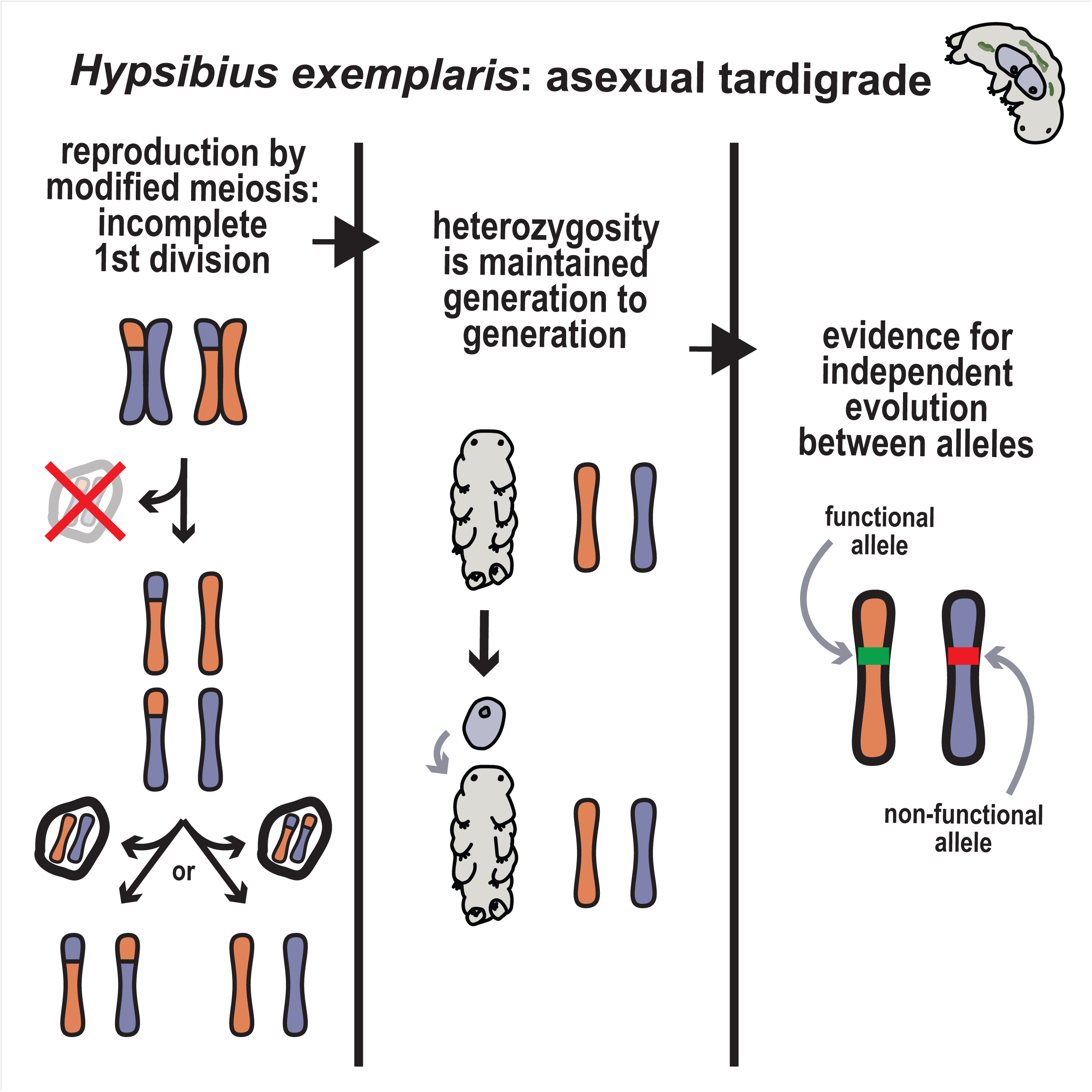
Summary of conclusions. Heterozygosity is maintained either by co-segregation of recombinant chromatids (as shown) or by complete inhibition of meiotic crossovers. Alt text: Summary figure titled “*Hypsibius exemplaris*: asexual tardigrade” with three columns, each with corresponding diagrams. The columns are labeled: “reproduction by modified meiosis: incomplete first division”, “heterozygosity is maintained generation to generation”, and “evidence for independent evolution between alleles”.

Abortive meiosis I parthenogenesis with independent assortment and crossing over will result in loss of heterozygosity distal to crossovers, with near-complete loss over time (Omilian et al. 2006; Xu et al. 2011; Blanc et al. 2025). However, we found no evidence of loss of heterozygosity in *H. exemplaris*, neither at specific loci across generations nor in the pattern of heterozygosity across putative chromosomes. Our experiment to track allele inheritance over generations did reveal a single instance where inheritance of both alleles at one locus could not be confirmed; however, the technical limitations of this assay combined with our later finding that heterozygosity is consistent across the genome strongly support the conclusion that this single failure to detect inheritance of both alleles was a technical artifact rather than a biological reality. Of course, we cannot rule out the possibility that rare LOH events remain undetected, including transient events rapidly removed by purifying selection, small tracts of LOH occuring throughout the genome due to gene conversion, or even cryptic sex. Nonetheless, we conclude from our findings that *Hypsibius exemplaris* largely maintains heterozygosity through parthenogenetic reproduction. Currently, an average of 1.40% of bases are heterozygous in *H. exemplaris* individuals, which is similar to the levels reported in other clonal parthenogens (Brandt et al. 2021; Blanc et al. 2023; Lacy et al. 2024). Importantly, it is unknown for how long *H. exemplaris* has been reproducing asexually, so it remains an open question whether heterozygosity in this species is continuing to rise or has already reached an equilibrium between mutations, selection, and potential rare LOH events.

A few possibilities may explain the mechanism behind how heterozygosity is maintained in *H. exemplaris*. On the one hand, it is unlikely that selection against loss of heterozygosity events, alone, could explain our findings, particularly given consistently high offspring survival rates in this species (Goldstein 2018b). Instead, a more likely explanation is that heterozygosity is preserved by an additional modification of meiosis. For example, it is possible that meiotic crossovers do not occur at all. We attempted multiple experiments to address this question, including allele-specific oligo-FISH and EdU labeling of nascent DNA (Blanc et al. 2023), but we have yet to find a method that produces informative results. However, structures that have been interpreted as synaptonemal complexes have been observed before (Jezierska et al. 2021), *H. exemplaris* does have the genetic potential to undergo meiotic crossovers: In most species, crossovers are initiated by Spo11-induced double-strand breaks (Keeney et al. 1997) and promoted by the conserved MutS homologs Msh4 and Msh5 (Hollingsworth et al. 1995; Hunter 2015), and we found by reciprocal BLASTp against *Caenorhabditis elegans* that *Hypsibius exemplaris* contains homologs for all three proteins (Spo11: BV898_09712, Msh4: BV898_19257, and Msh5: BV898_08111). If *H. exemplaris* does undergo regular meiotic crossovers, heterozygosity may still be maintained by co-segregation of recombinant chromatids during meiosis II, as observed in two other abortive meiosis I parthenogens (Figure 6) (Blanc et al. 2023; Lacy et al. 2024).

If heterozygosity is maintained over sufficient time, we expect to see sequence divergence between alleles that leads, in some instances, to a divergence in function (Welch and Meselson 2000; Pouchkina-Stantcheva et al. 2007; Brandt et al. 2021; Öztoprak et al. 2025). In line with this idea, we have identified multiple transcriptionally expressed genes that harbor predicted loss-of-function mutations in only one allele in *H. exemplaris*. Although preliminary, our findings are consistent with the idea that *H. exemplaris* has been reproducing clonally for long enough for the predicted functional divergence to occur in a subset of genes. We found no evidence in an existing RNA-seq dataset that allele-specific gene expression has also evolved at these sites. We might not expect reductions in the expression of non-functional alleles if the observed functional divergence is relatively new, but the dearth of evidence for allele-specific expression could also have resulted from the low statistical power associated with use of an RNA-seq dataset that was not originally collected for this purpose. Future work to compare heterozygosity levels between tardigrade lineages may shed additional light on the question of how long this and other tardigrade species have been reproducing asexually.

Maintenance of heterozygosity in asexual species is expected to have complex evolutionary consequences that differ between the short- and long-term. Loss of heterozygosity typically has an immediate fitness cost for newly non-clonal parthenogenetic or inbreeding populations due to the phenotypic expression of recessive deleterious alleles previously masked by heterozygosity (Engelstädter 2008; Charlesworth and Willis 2009). Therefore, in the short-term, maintaining heterozygosity avoids this inbreeding depression. On the other hand, long-term clonal reproduction is predicted to have drastic negative consequences on finite populations due to Hill-Robertson interference, whereby linkage between loci without recombination reduces the efficiency of selection (Hill and Robertson 1966; Agrawal 2006). Hill-Robertson interference results in a reduced ability to adapt to changing environments and an irreversible accumulation of deleterious mutations over time (Muller’s Ratchet) (Muller 1964; Lynch et al. 1993). This expectation has contributed to the long-standing view that clonal asexual lineages represent evolutionary dead ends compared to both non-clonal asexual lineages and sexually reproducing populations (Smith 1978; Neiman et al. 2017). However, *H. exemplaris* now adds to a growing list of well-characterized clonal parthenogenetic animals (Blanc et al. 2025). Whether these species are doomed in the long run, represent rare survivors against the usual fate of clonal reproduction, occasionally reproduce in a non-clonal fashion, or call into question the dead-end rule itself, remains to be determined.

## Data Availability Statement

All code generated to analyze data for this study is provided in File S1. All previously published sequencing datasets that were analyzed are listed in Table S2. The authors declare that all other data supporting the results of this study are available within the manuscript or Supplementary Materials.

## Acknowledgements

We thank the members of the Goldstein and Burch labs for support and feedback on this project, including Kira Heikes and Courtney Clark-Hachtel for training in tardigrade-specific techniques. We thank Nat Prunet for assistance in the UNC Biology microscopy core, Corbin Jones at the UNC Bioinformatics and Analytics Research Collaborative (BARC) for feedback on data analysis methodology, and Ata Kalirad and Jeff Sekelsky for comments on the manuscript.

## Author Contributions

A.C. conceived and designed the experiments with contributions from B.G.; A.C., S.Z., and L.P. performed the experiments; A.C., C.B., and S.Z. analyzed the data; A.C. and S.Z. prepared the digital images; A.C., B.G., and C.B. drafted the article.

