## Supplementary figures and images for "Modified meiosis in the tardigrade *Hypsibius exemplaris* maintains heterozygosity across the genome"

### Figure S1

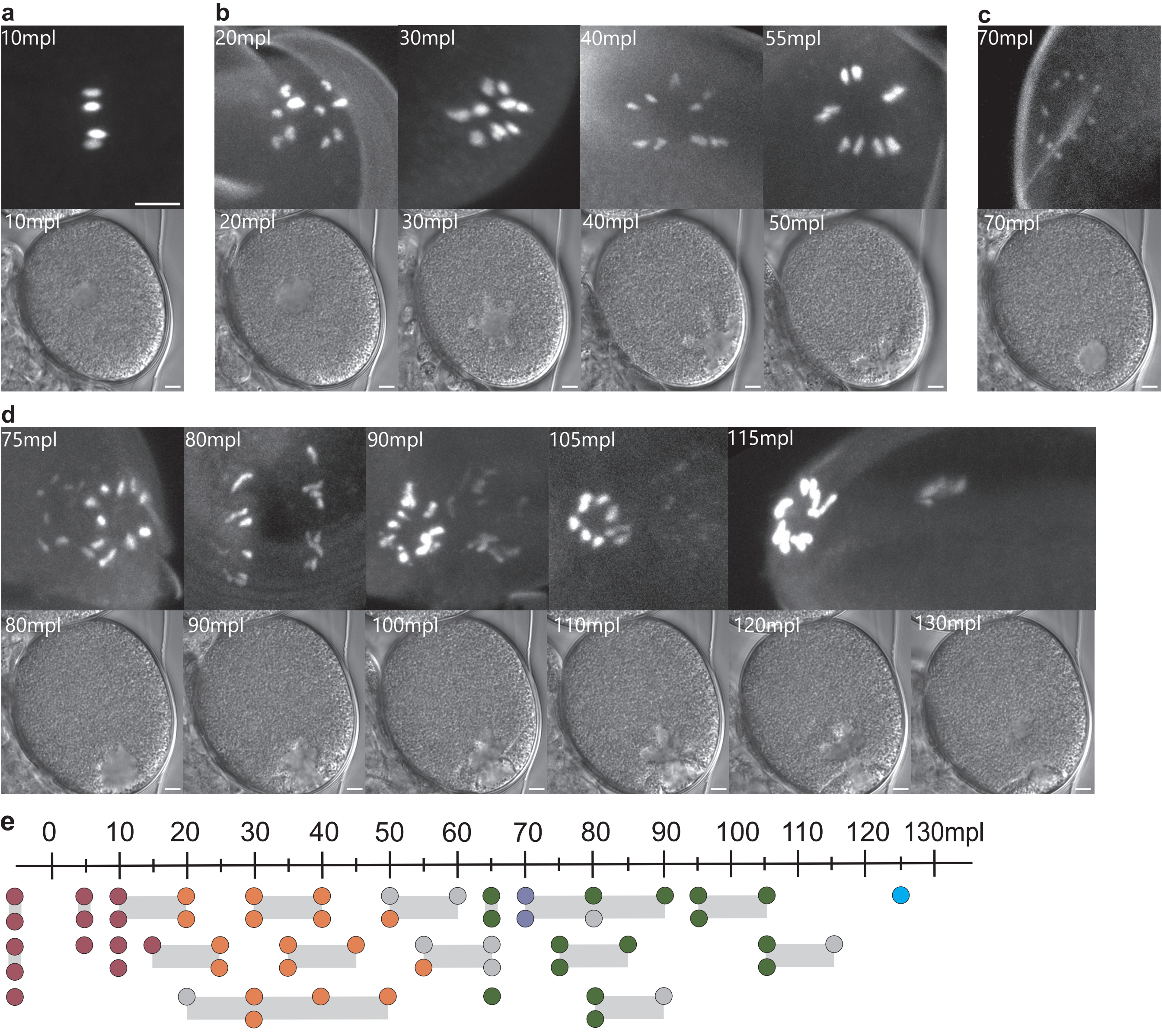

### Figure S2

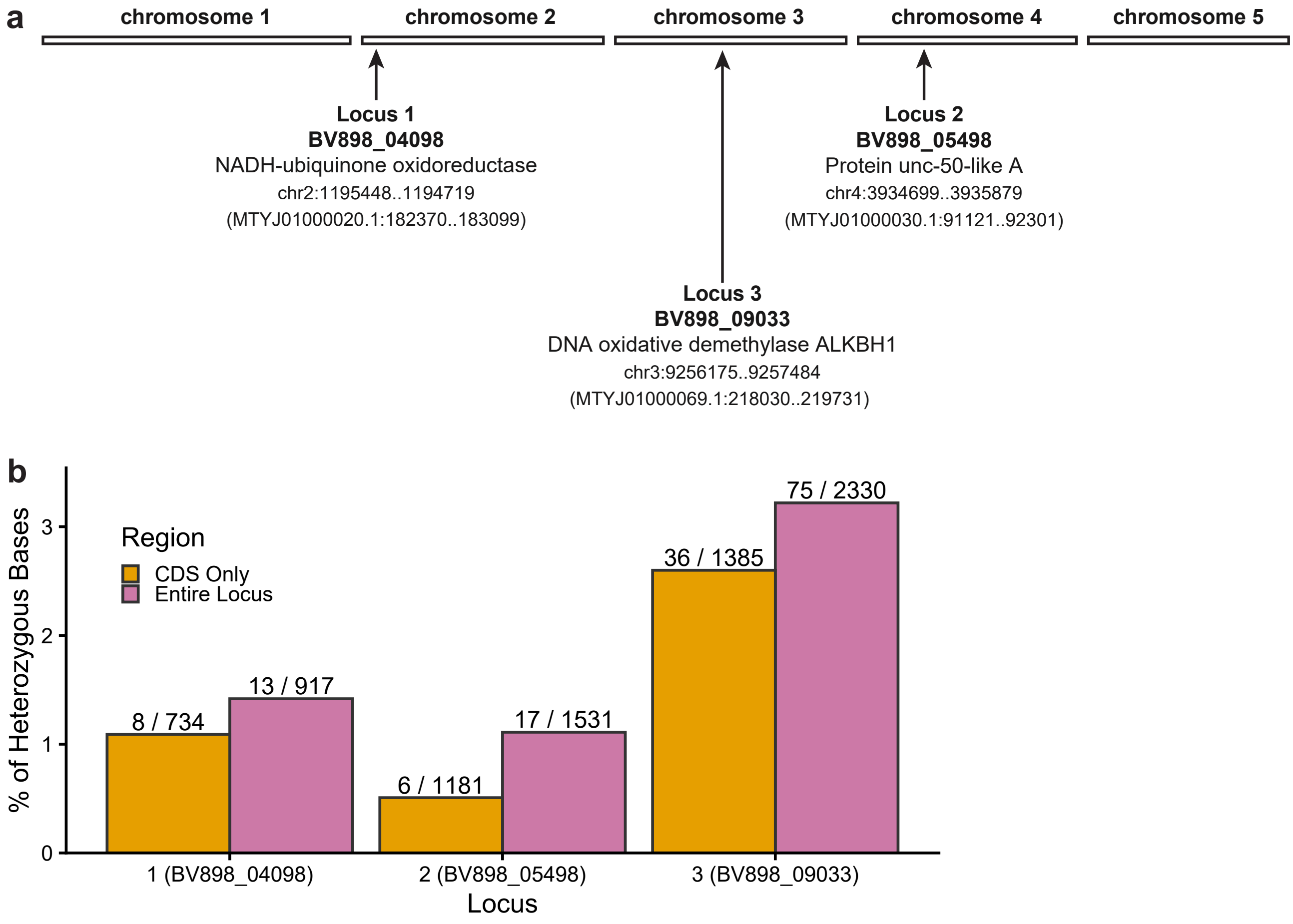

### Figure S3

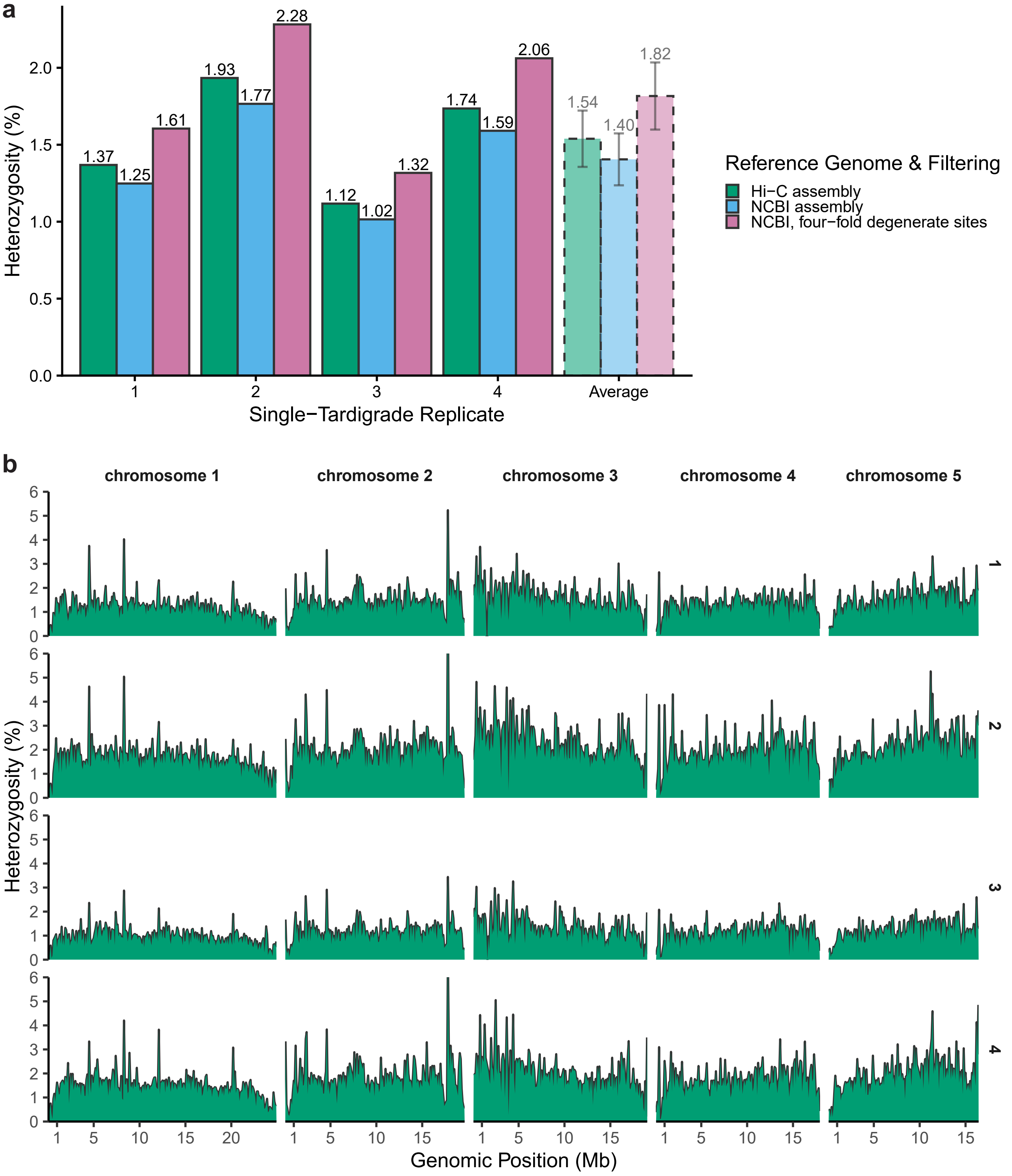

### Figure S4

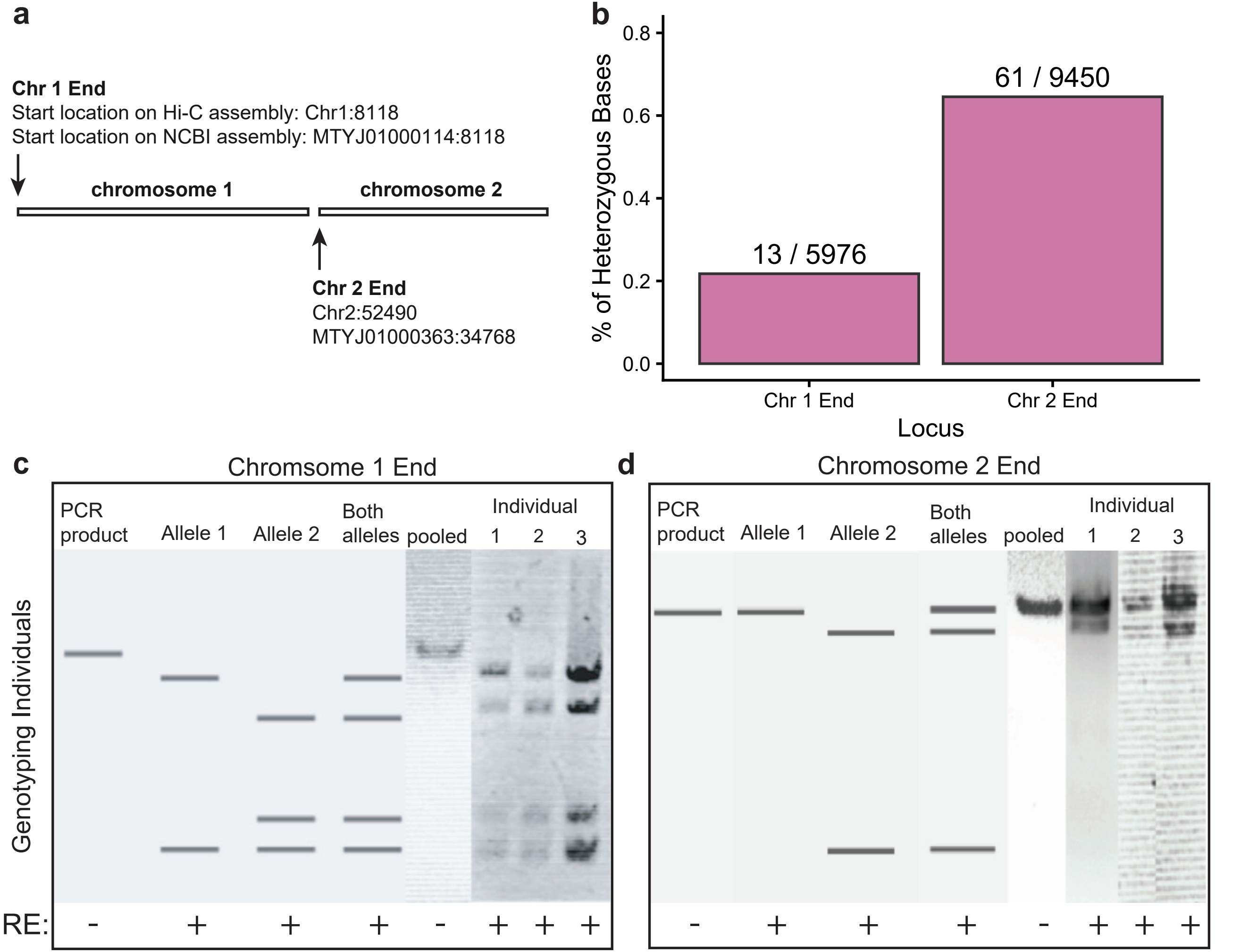
