## Supplementary material for "Modified meiosis in the tardigrade *Hypsibius exemplaris* maintains heterozygosity across the genome": File S1: allele_specific_expression_analysis_pipeline.sh

```
#!/bin/bash
set -euo pipefail
trap 'echo "Error on line $LINENO: $BASH_COMMAND"; exit 1' ERR

# Load modules
module load bwa
module load samtools
module load gcc
module load bcftools

#SBATCH --job-name=aseq
#SBATCH --output=aseq.out
#SBATCH --cpus-per-task=8
#SBATCH --mem=64g
#SBATCH --time=6:00:00

###############################################
# USER SETTINGS
###############################################
# Usage:
#   This script:    1) maps RNA-seq reads to generate BAM files and
#                   2) runs ASEQ for allele-specific expression analysis (ASE).
#
#   Revise user inputs to match your file paths and sample names.
#
#   Requires previously generated VCF files of heterozygous variants from WGS analysis.
###############################################

# Set working directory (where ASE results will be saved)
cd /work/users/a/d/adricoke/single_tardigrade_seq_analysis
if [ ! -d allele_specific_expression_analysis ]; then
    mkdir allele_specific_expression_analysis
fi
cd allele_specific_expression_analysis
aseq_workdir=$(pwd)

# ASEQ program location
aseq_dir=/nas/longleaf/home/adricoke/aseq/binaries/linux64

# Reference genome and gene annotation file (GFF) are the same for all samples.
ref=/work/users/a/d/adricoke/single_tardigrade_seq_analysis/refs/Hypsibius_exemplaris/NCBI/GCA_002082055.1/GCA_002082055.1_nHd_3.1_genomic.fna
genes=/work/users/a/d/adricoke/single_tardigrade_seq_analysis/refs/Hypsibius_exemplaris/NCBI/GCA_002082055.1/genomic.gff

### RNA-seq Raw Data ###
# Directory containing raw RNA-seq data files (FASTQ)
rnaseq_data_dir=/work/users/a/d/adricoke/single_tardigrade_seq_analysis/data/Yoshida2017_RNAseq_reads_10K_Active
# Sample names (prefixes of FASTQ files) for RNA-seq data
RNASEQ_SAMPLES=("SRR5218239" "SRR5218240" "SRR5218241")
# Note: Pipeline assumes paired-end RNA-seq with filenames like SRR5218239_1.fastq and SRR5218239_2.fastq

### Previously Generated Variant VCF Files ###
# Directory containing variant VCF files from WGS analysis
variants_dir=/work/users/a/d/adricoke/single_tardigrade_seq_analysis/NCBI_genome_results
### Heterozygous variants identified from bulk datasets
VARIANTS_LIST=("Hypsibius_exemplaris_bulk1/heterozygous.vcf.gz" \
               "Hypsibius_exemplaris_bulk2/heterozygous.vcf.gz" \
               "Hypsibius_exemplaris_bulk3/heterozygous.vcf.gz")
### Heterozygous variants identified from individual tardigrade sequencing replicates
REPLICATE_LIST=("Hypsibius_exemplaris_rep1/heterozygous.vcf.gz" \
                "Hypsibius_exemplaris_rep2/heterozygous.vcf.gz" \
                "Hypsibius_exemplaris_rep3/heterozygous.vcf.gz" \
                "Hypsibius_exemplaris_rep4/heterozygous.vcf.gz")

###############################################
# MERGE VCF FILES TO ONLY KEEP SHARED HETEROZYGOUS VARIANTS
###############################################
# KEEP ONLY:
#   1) sites shared across all 3 bulk VCFs
#   2) sites present in at least 1 individual replicate VCF
###############################################
echo
echo "Finding shared heterozygous sites..."
echo

# Step 1:
# Keep only variants present in all 3 bulk VCFs
bcftools isec \
  -n=3 \
  -Oz \
  -p bulk_isec \
  "${variants_dir}/${VARIANTS_LIST[0]}" \
  "${variants_dir}/${VARIANTS_LIST[1]}" \
  "${variants_dir}/${VARIANTS_LIST[2]}"

# Step 2:
# Merge replicate VCFs into a union of replicate sites
bcftools merge \
  --force-samples \
  -Oz -o merged_replicates.vcf.gz \
  "${variants_dir}/${REPLICATE_LIST[0]}" \
  "${variants_dir}/${REPLICATE_LIST[1]}" \
  "${variants_dir}/${REPLICATE_LIST[2]}" \
  "${variants_dir}/${REPLICATE_LIST[3]}"

bcftools index merged_replicates.vcf.gz

# Step 3:
# Keep only shared bulk variants also found in >=1 replicate
bcftools isec \
  -n=2 \
  -w1 \
  -Oz \
  -p final_isec \
  bulk_isec/0000.vcf.gz \
  merged_replicates.vcf.gz

# Step 4:
# Flatten to positions only and keep ASEQ-safe SNPs
bcftools view \
  -m2 -M2 \
  -v snps \
  -G \
  -Ov -o shared_heterozygous_variants.vcf \
  final_isec/0000.vcf.gz

###############################################
# (ONCE PER REFERENCE) GENERATE GENES.BED FROM GENE ANNOTATION GFF
###############################################

if [ ! -f genes.bed ]; then
    echo
    echo "Generating genes.bed from genes.gff..."
    echo

    awk 'BEGIN{OFS="\t"}
     !/^#/ && $3=="gene" {
         # GFF: 1-based inclusive; BED: 0-based, start inclusive, end exclusive
         # extract a clean gene ID, e.g. BV898_00001 from ID=gene-BV898_00001
         id = "."
         split($9, a, ";")
         for (i in a) {
             if (a[i] ~ /^ID=/) {
                 tmp = a[i]
                 sub(/^ID=/, "", tmp)
                 # strip "gene-" prefix if present
                 sub(/^gene-/, "", tmp)
                 id = tmp
             }
         }
         print $1, $4-1, $5, id, 0, $7
        }' $genes > genes.bed

else
    echo
    echo "genes.bed already exists. Skipping generation."
    echo
fi

###############################################
# ALIGN RNA-SEQ READS TO REFERENCE GENOME
###############################################
echo
echo "Aligning RNA-seq reads to reference genome and creating bamlist.txt and vcflist.txt..."
echo
> bamlist.txt
> vcflist.txt

cd $rnaseq_data_dir

for S in "${RNASEQ_SAMPLES[@]}"; do
    reads1="${S}_1.fastq"
    reads2="${S}_2.fastq"

    echo "Aligning ${S}..."
    bwa mem -t 8 "${ref}" "${reads1}" "${reads2}" \
        | samtools sort -o "${S}.sorted.bam"

    samtools index "${S}.sorted.bam"

    # append mapped RNAseq data to bamlist.txt
    echo "${rnaseq_data_dir}/${S}.sorted.bam" >> "${aseq_workdir}/bamlist.txt"

    # append merged VCF to vcflist.txt (same for all samples - append once per RNA-seq sample)
    echo "${aseq_workdir}/shared_heterozygous_variants.vcf" >> "${aseq_workdir}/vcflist.txt"
done

cd $aseq_workdir

###############################################
# ALLELE-SPECIFIC EXPRESSION ANALYSIS WITH ASEQ
###############################################
echo
echo "Running allele-specific expression analysis with ASEQ..."
echo

# ASEQ parameters:
# mbq=10 minimum base quality
# mrq=20 minimum read quality
# mdc=10 minimum depth coverage
# pht=0.05 p-value heterozygosity test
# pft=0.05 p-value frequency test

$aseq_dir/ASEQ \
  vcflist=vcflist.txt \
  bamlist=bamlist.txt \
  genes=genes.bed \
  mode=ASE \
  threads=8 \
  mbq=10 \
  mrq=20 \
  mdc=10 \
  pht=0.05 \
  pft=0.05 \
  out=aseq_results

###############################################
# DONE
###############################################
echo
echo "Pipeline complete."
echo "Results can be found in ./aseq_results"
echo
```
