## Supplementary material for "Modified meiosis in the tardigrade *Hypsibius exemplaris* maintains heterozygosity across the genome": File S1: Analyze_Heterozygosity_at_Chr_Ends.ipynb

```
# %% [code]
"run this function if needed: for fasta files with two consensus sequences combined in one"

#split one fasta file with multiple sequences into separate fasta files 
def split_fasta(input_file):
    sequences = fasta_to_list(input_file)
    
    for name, seq in sequences:
        # use the sequence header name as the filename
        output_file = f"{name}.fasta"
        with open(output_file, "w") as f:
            f.write(f">{name}\n")
            f.write("".join(seq) + "\n")
        print(f"Written: {output_file}")
consensus_sequences = "chrom2left_consensus_sequences.fasta"

split_fasta(consensus_sequences)

# %% [code]
import csv
import os

###user inputs###

"""requires separate fasta files and use split_fasta function if necessary"""
#read in fasta file

loci = [
    {
        "name": "chrom1left",
        "top": "chrom1left_consensus_consensus2.fasta",
        "bottom": "chrom1left_consensus_consensus1.fasta",
    },
    
    # separate files (one sequence per file)
    {
        "name": "chrom2mid",
        "bottom": "Chr-2M_results/chrom2mid_primer2_bottomaligned.fasta",
        "top":    "Chr-2M_results/Chrom2mid_primer2_topaligned.fasta",
    },
    {
        "name": "chrom2left",
        "top": "chrom2left_consensus_consensus2.fasta",
        "bottom": "chrom2left_consensus_consensus1.fasta",
    },
]

###Python script from all loci  

#convert fasta file to list in [[name,list]]
def fasta_to_list(fasta_file_name): 
    consensus = [] 
    name = None 
    seq = []

    with open(fasta_file_name,"r") as f: 
        for line in f:
            line = line.strip()

            if line.startswith(">"):
                if name is not None: 
                    consensus.append([name, seq])

                name = line[1:]
                seq = []
            else: 
                seq.extend(line)
        if name is not None: 
            consensus.append([name,seq])
        return consensus  

# --- extract locus name from filename ---
locus_name = os.path.splitext(os.path.basename(bottom_fasta_file_name))[0]

#create key dictionary for different counts 
def count_heterozygosity(seq1, seq2):
    counts = {"heterozygous": 0, "indels": 0, "homozygous": 0}
    for i in range(len(seq1)):
        if seq1[i] != seq2[i]:
            counts["heterozygous"] += 1
            if seq1[i] == "-" or seq2[i] == "-":
                counts["indels"] += 1
        else:
            counts["homozygous"] += 1
    return counts

# --- write to CSV ---
output_csv = "results.csv"


with open(output_csv, "a", newline="") as csvfile:
    fieldnames = ["locus", "heterozygous", "indels", "homozygous"]
    writer = csv.DictWriter(csvfile, fieldnames=fieldnames)
    writer.writeheader()

    for locus in loci:
        seq1 = fasta_to_list(locus["bottom"])[0][1]
        seq2 = fasta_to_list(locus["top"])[0][1]

        counts = count_heterozygosity(seq1, seq2)

        writer.writerow({
            "locus": locus["name"],
            "heterozygous": counts["heterozygous"],
            "indels": counts["indels"],
            "homozygous": counts["homozygous"]
        })

print(f"Processed: {locus['name']} → {counts}")

# %% [code]
```
