## Supplementary material for "Modified meiosis in the tardigrade *Hypsibius exemplaris* maintains heterozygosity across the genome": File S1: call_filter_and_annotate_variants_pipeline.sh

```
#!/bin/bash
set -euo pipefail
trap 'echo "Error on line $LINENO: $BASH_COMMAND"; exit 1' ERR

###############################################
# USER SETTINGS
###############################################
# Usage:
#   Single-end: ./script.sh <reference.fa> <reads.fastq[.gz]> [snpeff_database snpeff_dir]
#   Paired-end: ./script.sh <reference.fa> <reads_1.fastq[.gz]> <reads_2.fastq[.gz]> [snpeff_database snpeff_dir]

if [ "$#" -lt 2 ]; then
    echo "Usage:"
    echo "  $0 <reference.fa> <reads.fastq[.gz]> [snpeff_database snpeff_dir]"
    echo "  $0 <reference.fa> <reads_1.fastq[.gz]> <reads_2.fastq[.gz]> [snpeff_database snpeff_dir]"
    exit 1
fi

ref="$1"
reads1="$2"

# Detect paired-end vs single-end
if [[ "$#" -eq 3 || "$#" -eq 5 ]]; then
    # Paired-end mode
    reads2="$3"
    shift 3
else
    # Single-end mode
    reads2=""
    shift 2
fi

# Optional snpEff settings
if [ "$#" -eq 2 ]; then
    snpeff_database="$1"
    snpeff_dir="$2"
else
    snpeff_database="NA"
    snpeff_dir="NA"
fi

echo "Reference genome: $ref"
if [ -n "$reads2" ]; then
    echo "Reads (paired-end):"
    echo "  R1: $reads1"
    echo "  R2: $reads2"
else
    echo "Reads (single-end): $reads1"
fi
echo "SnpEff database: $snpeff_database"
echo "SnpEff directory: $snpeff_dir"
echo

###############################################
# MAP READS TO REF
###############################################
echo "Mapping reads to reference genome with BWA..."
echo

if [ -n "$reads2" ]; then
    # Paired-end
    bwa mem "$ref" "$reads1" "$reads2" > reads_to_ref.sam
else
    # Single-end
    bwa mem "$ref" "$reads1" > reads_to_ref.sam
fi

samtools view -bShu reads_to_ref.sam | samtools sort -o sorted.bam
samtools index sorted.bam

###############################################
# CALL VARIANTS
###############################################

# -Ou and -Ob arguments maintain binary format to speed computation
# -a FORMAT/DP,FORMAT/AD adds depth and allele depth to vcf file
# -f identifies reference fasta file
# -m option calls multiple alleles (versus -c whch only generates consensus sequences)
# -o specifies output file

echo "Calling variants with bcftools..."
echo

bcftools mpileup -Ou \
-a FORMAT/DP,FORMAT/AD \
-f $ref \
sorted.bam | \
bcftools call -m -Ob -o variants.vcf.gz

bcftools index variants.vcf.gz

###############################################
# FILTER VARIANTS
###############################################

# remove low quality (QUAL<30) & low coverage (DP<40) variants
# and keep only heterozygous variants
    # heterozygous variants have minor allele frequency between 0.25 and 0.5; minor allele may be ALT or REF
    # minor allele = min between AD[0] (REF) and AD[1] (ALT)
    # Use FORMAT/DP for total depth at position, FORMAT/AD for allele depths

echo "Filtering variants..."
echo

# filter by quality and depth
bcftools view -i 'QUAL>=30 && FORMAT/DP[0]>=40' variants.vcf.gz -Oz -o filtered.vcf.gz

bcftools index filtered.vcf.gz

# filter for heterozygous variants
bcftools view -i \
'((FORMAT/AD[0:0]/FORMAT/DP[0]>=0.25 && FORMAT/AD[0:0]/FORMAT/DP[0]<=0.5) || (FORMAT/AD[0:1]/FORMAT/DP[0]>=0.25 && FORMAT/AD[0:1]/FORMAT/DP[0]<=0.5))' \
filtered.vcf.gz -Oz -o heterozygous.vcf.gz

bcftools index heterozygous.vcf.gz

###############################################
# ANNOTATE VARIANTS WITH SnpEff
###############################################

if [ "$snpeff_database" = "NA" ] || [ "$snpeff_dir" = "NA" ] || [ ! -d "$snpeff_dir/data/$snpeff_database" ]; then
    echo "Skipping SnpEff annotation as database or directory not provided or not found."
    echo
    echo "To identify and download a SnpEff database (example shown for Hypsibius exemplaris):"
    echo "  # Once per user, download SnpEff to user's home directory and unzip"
    echo "      $ wget https://snpeff.odsp.astrazeneca.com/versions/snpEff_latest_core.zip"
    echo "      $ unzip snpEff_latest_core.zip"
    echo "  # Once per species..."
    echo "  # Move into user's SnpEff directory"
    echo "      $ cd snpEff"
    echo "  # Check for database and find correct name"
    echo "      $ java -jar snpEff.jar databases |grep -i Hypsibius"
    echo "  # Download database"
    echo "      $ java -jar snpEff.jar download Hypsibius_exemplaris_gca002082055v1"
    echo
else
    echo "Annotating variants with SnpEff..."
    echo

    ## Rename chromosomes if rename_chromosomes.txt exists
        # Format of rename_chromosomes.txt: (tab-separated; need to generate manually in advance)
        # old_name1    new_name1
        # old_name2    new_name2
        # ...
    if [ -f rename_chromosomes.txt ]; then
        # save backup of original filtered VCF
        cp heterozygous.vcf.gz heterozygous.vcf.gz.orig
        # rename chromosomes
        bcftools annotate --rename-chrs rename_chromosomes.txt -o heterozygous.tmp.vcf.gz -Oz heterozygous.vcf.gz
        # replace original filtered VCF with renamed version & re-index
        mv heterozygous.tmp.vcf.gz heterozygous.vcf.gz
        bcftools index -f heterozygous.vcf.gz
    fi

    java -Xmx4g -jar $snpeff_dir/snpEff.jar \
      -v \
      -dataDir $snpeff_dir/data \
      $snpeff_database \
      heterozygous.vcf.gz > annotated.vcf
fi

###############################################
# OUTPUT VARIANT INFO AS CSV FILES
###############################################

# QUAL is quality score
# CHROM/POS specify position on reference
# FORMAT/DP is depth
# FORMAT/AD{0} is allele depth for reference allele, FORMAT/AD{1} is allele depth for alternate allele
# %ANN field contains multiple subfields separated by | character; see SnpEff documentation for details

echo "Generating filtered, heterozygous variant CSV..."
echo

if [ -f annotated.vcf ]; then
    bcftools query -f '%CHROM,%POS,%REF,%ALT,%QUAL[,%DP,%AD{0},%AD{1}][,%ANN]' -o heterozygous_variants.csv annotated.vcf
else
    # If no SnpEff annotation was done, output filtered variants without ANN field
    bcftools query -f '%CHROM,%POS,%REF,%ALT,%QUAL[,%DP,%AD{0},%AD{1}]' -o heterozygous_variants.csv heterozygous.vcf.gz
fi

echo "Generating filtered, all-genotype variant CSV..."
echo
bcftools query -f '%CHROM,%POS,%REF,%ALT,%QUAL[,%DP,%AD{0},%AD{1}]' -o all_variants.csv filtered.vcf.gz

###############################################
# DONE
###############################################

echo "Pipeline complete."
echo "Main output written to: heterozygous_variants.csv and all_variants.csv"
echo "Additional outputs: reads_to_ref.sam, sorted.bam, variants.vcf.gz, filtered.vcf.gz, heterozygous.vcf.gz, annotated.vcf, [snpEff_genes.txt, snpEff_summary.html]"
```
