## Supplementary material for "Modified meiosis in the tardigrade *Hypsibius exemplaris* maintains heterozygosity across the genome": File S1: Chr_Ends_Heterozygosity_Plotting.R

```
###############################################################################
# Specify File locations
###############################################################################
library(ggplot2)
library(readr)
library(dplyr)
library(plyr)
library(tidyverse)

# change to match file paths on your computer
main_dir = "C:/Users/addie/OneDrive - University of North Carolina at Chapel Hill/LAB/tardigrade_heterozygosity/Aurora_He_meiosis_paper_results"
setwd(main_dir)
getwd()
plots_dir = "C:/Users/addie/OneDrive - University of North Carolina at Chapel Hill/LAB/tardigrade_heterozygosity/He_FIGURE_PLOTS"

###############################################################################
# Plot Aesthetics
###############################################################################

### Adjusts settings for ALL plots ###

# theme settings
theme_set(theme_classic())
theme_update(text=element_text(size=10), # IS USUALLY 10
             strip.background = element_blank(), # remove box around facet labels
             strip.text = element_text(face = "bold"), # and bold text
             legend.key.size = unit(0.1, "in") # adjust legend sizing
)

cbPalette <- c("#999999", "#56B4E9", "#E69F00", "#CC79A7", "#009E73")

###############################################################################
# Read in Data
###############################################################################
data_files <- "Heterozygosity results.csv"

mydata<- read.csv(data_files)

head(mydata)

###############################################################################
# calculations
###############################################################################

mydata$ratio_het_sites_withindels <- (mydata$heterozygous/(mydata$heterozygous+mydata$homozygous))

mydata$ratio_het_sites_no_indels <- ((mydata$heterozygous-mydata$indels)/(mydata$heterozygous+mydata$homozygous-mydata$indels))

mydata$ratio_indels_in_het_sites <- (mydata$indels/mydata$heterozygous)

###############################################################################
# plots!
###############################################################################

### barplot of % of het sites per locus, EXCLUDING indels

plotting_data <- mydata %>%
  subset(locus %in% c("chrom1left", "chrom2left")) # only plot these 2 loci

# re-label loci
plotting_data$locus <- factor(plotting_data$locus,
                       levels = c("chrom1left", "chrom2left"),
                       labels = c("Chr 1 End", "Chr 2 End"))

ggplot(data=plotting_data) + 
  geom_col(aes(x = locus, y = 100*ratio_het_sites_no_indels),
           fill = "#CC79A7", color = "grey20") + 
  # label bars with het sites count
  geom_text(aes(x = locus, y = 100*ratio_het_sites_no_indels, label = paste0(heterozygous-indels, " / ", heterozygous+homozygous-indels)), vjust = -0.5) +
  # scale_fill_manual(values=c("#E69F00", "#CC79A7")) +
  scale_y_continuous(limits = c(0,0.8))+
  theme(legend.position = "none") +
  labs(x = "Locus", y = "% of Heterozygous Bases")
plot_title <- "Chr_Ends_Loci_Het_Percentage_ignore_indels"
ggsave(filename = paste0(plots_dir,"/",plot_title, ".png" ), width = 3.5, height = 2.5, dpi = 300)
ggsave(filename = paste0(plots_dir,"/",plot_title, ".pdf" ), width = 3.5, height = 2.5)
```
