## Supplementary material for "Modified meiosis in the tardigrade *Hypsibius exemplaris* maintains heterozygosity across the genome": File S1: Compile_Heterozygosity_Data.R

```
library(tidyverse)
library(plyr)

###############################################################################
# User Settings (change if needed)
###############################################################################

## Set working directory
setwd("/work/users/a/d/adricoke/single_tardigrade_seq_analysis")
main_dir = getwd()

## Set directory to save summary data files
summary_files_dir <- "/work/users/a/d/adricoke/single_tardigrade_seq_analysis/all_refs_heterozygosity_results"

# ## Set bin sizes for across-genome analysis
# bin_sizes <- c(10000, 100000)
# 
# message("Bin sizes:")
# for (bin_size in bin_sizes) {
#   message(message(paste0(bin_size/1000, 'kb')))
# }

###############################################################################
# Find File locations and treatments
###############################################################################

## Identify variant files
# all_variants.csv includes only filtered variants (QUAL>=30, depth>=40), both heterozygous and homozygous
data_files <- dir(main_dir, recursive=T, include.dirs=T, pattern="all_variants.csv", full.names=T)
data_files

## Extract info from file path
sample_IDs = NULL
species = NULL
methods = NULL
replicates = NULL
references = NULL
for (file in data_files) {
  folder_name <- basename(dirname(file))
  sample_IDs <- c(sample_IDs, folder_name)
  # Check if folder contains "_rep" or "_bulk"
  if (grepl("_rep", folder_name)) {
    split_name <- unlist(strsplit(folder_name, "_rep"))
    species <- c(species, split_name[1])
    methods <- c(methods, "individual")
    replicates <- c(replicates, as.integer(split_name[2]))
  } else if (grepl("_bulk", folder_name)) {
    split_name <- unlist(strsplit(folder_name, "_bulk"))
    species <- c(species, split_name[1])
    methods <- c(methods, "bulk")
    replicates <- c(replicates, as.integer(split_name[2]))
  } else {
    species <- c(species, folder_name)
    methods <- c(methods, "individual") # defauly method = individual
    replicates <- c(replicates, 1)  # default replicate = 1
  }
  temp <- temp <- unlist(strsplit(file, "/"))
  temp2 <- ifelse(unlist(strsplit(temp[8], "new_"))[1] == "", unlist(strsplit(temp[8], "new_"))[2], unlist(strsplit(temp[8], "new_"))[1])
  references <- c(references, unlist(strsplit(temp2, "_genome_results"))[1])
}
sample_IDs
species
methods
replicates
references

# Filter file list to only species = Hypsibius_exemplaris
He <- species == "Hypsibius_exemplaris"
data_files <- data_files[He]
data_files[He]
sample_IDs[He]
species <- species[He]
methods <- methods[He]
replicates <- replicates[He]
references <- references[He]


# ## To filter to only CDS regions, read in cds_mask.csv files
# cds_data_files <- dir(main_dir, recursive=T, include.dirs=T, pattern="cds_mask.csv", full.names=T)
# cds_data_files
# 
# # Extract info from file path
# cds_references = NULL
# for (file in cds_data_files) {
#   temp <- unlist(strsplit(file, "/"))
#   cds_references <- c(cds_references, temp[10])
# }
# cds_references


## To filter to only 4-fold degenerate sites, read in all_4_fold_degenerate_sites.bed
four_fold_degenerate_sites_file <- "all_4_fold_degenerate_sites.bed"

###############################################################################
# Read in ALL Data & determine genotype per site
###############################################################################

## Variants data
df <- as.data.frame(NULL)
for (i in 1:length(data_files)) {
# for (i in 4) { # FOR TESTING: NCBI ref, individual rep 1

  # Read in Data
  mydata = readr::read_csv(data_files[i], col_names=FALSE)
  message(data_files[i])
  
  # name columns and remove SnpEff columns (if they exist)
  colnames(mydata) <- c("scaffold","position","REF","ALT","QUAL","depth","REF_depth","ALT_depth")
  mydata <- select(mydata, c("scaffold","position","REF","ALT","QUAL","depth","REF_depth","ALT_depth"))
  # head(mydata)
  
  # mydata <- subset(mydata, position < bin_size*2) # 2 bins only (per scaffold) FOR TESTING
  # summary(mydata)
  
  # Add additional columns for sample ID, species, method, replicate, & reference
  mydata$sample_ID <- sample_IDs[i]
  mydata$species <- species[i]
  mydata$method <- methods[i]
  mydata$replicate <- replicates[i]
  mydata$reference <- references[i]
  
  # when there are no ALT allele reads, the ALT_depth column is a '.' instead of a 0 in this csv
  # replace '.'s with 0s
  mydata$ALT_depth <- ifelse(mydata$ALT_depth=='.', 0, mydata$ALT_depth)
  # ALT_depth and QUAL also need to be converted to numeric
  mydata$ALT_depth <- as.numeric(mydata$ALT_depth)
  mydata$QUAL <- as.numeric(mydata$QUAL)
  
  # ALT allele is not necessarily the minor allele, so calculate minor allele frequency
  mydata$minor_allele_frequency <- ifelse(
    mydata$ALT_depth>mydata$REF_depth, # if ALT_depth is greater than REF_depth
    mydata$REF_depth/mydata$depth, # then minor allele frequency is REF_depth/depth
    mydata$ALT_depth/mydata$depth) # otherwise, minor allele frequency is ALT_depth/depth
  # minor allele frequency will be between 0 and 0.5
  
  # call each position as either heterozygous (1) or homozygous (0)
  # heterozygous if minor allele frequency is between 0.25-0.5
  # if coverage = 40 (current minimum), the chance of a true heterozygous allele showing up at 0.25 frequency
  # (assuming coin-flip probability for each read) is 0.000771, or 0.0771%
  # at higher coverage, the chance of this happening gets smaller & smaller
  # homozygous if minor allele frequency = 0 (error rate should be very low)
  mydata$heterozygosity = ifelse(
    mydata$minor_allele_frequency > 0.25, 1, ifelse(mydata$minor_allele_frequency == 0, 0, '.'))
  
  # remove NULL positions where genotype could not be called by these metrics (high error rate or other issue)
  mydata <- subset(mydata, heterozygosity != '.')
  mydata$heterozygosity <- as.numeric(mydata$heterozygosity)
  
  # Add to combined df
  df <- rbind(df, mydata)
}
# head(df)


# ## CDS masks
# cds_masks <- as.data.frame(NULL)
# for (i in 1:length(cds_data_files)) {
# 
#   # Read in data
#   mydata <- readr::read_csv(cds_data_files[i], col_names=FALSE)
#   message(cds_data_files[i])
#   
#   # Name columns
#   colnames(mydata) <- c("scaffold","position","CDS_coverage")
#   
#   # mydata <- subset(mydata, position < bin_size*2) # 2 bins only (per scaffold) FOR TESTING
#   
#   # Assign reference from file path
#   mydata$reference <- cds_references[i]
#   
#   # Add to combined df
#   cds_masks <- rbind(cds_masks, mydata)
# }
# # head(cds_masks)


## Four-fold degenerate sites
four_fold_degenerate_sites <- read.table(four_fold_degenerate_sites_file, header=FALSE)
colnames(four_fold_degenerate_sites) <- c("scaffold","NA","position","transcript","codon","NA")


###############################################################################
# Flag each site as CDS or non-CDS (both filtered & unfiltered versions will be used for downstream analysis)
###############################################################################

# Create unique site key for entire df
df$site_key <- paste(df$scaffold, df$position, df$reference, sep = ":")

# # Create unique site key for CDS sites
# cds_keys <- unique(paste(cds_masks$scaffold, cds_masks$position, cds_masks$reference, sep = ":"))
# 
# # Flag each site as CDS TRUE/FALSE
# df$CDS <- df$site_key %in% cds_keys
# 
# message("CDS filter applied.")

###############################################################################
# Flag each site as four-fold-degenerate or not (both filtered & unfiltered versions will be used for downstream analysis)
###############################################################################

# Create unique site key for four-fold-degenerate sites
ffd_keys <- unique(paste(four_fold_degenerate_sites$scaffold, four_fold_degenerate_sites$position, "NCBI", sep = ":"))

# Flag each site as four-fold-degenerate TRUE/FALSE
df$four_fold_degenerate <- df$site_key %in% ffd_keys

# Drop the helper column
df$site_key <- NULL

message("Four fold degenerate sites filter applied.")

###############################################################################
# Compile & Summarize Data
###############################################################################

# ## With and without CDS filter (no FFD filter), both references, summary per sample AND per bin across the genome
# summary_df <- as.data.frame(NULL)
# for (CDS_filter in c(TRUE,FALSE)) {
#   
#   # Apply CDS filter (or not)
#   if (CDS_filter == TRUE) {
#     mydata <- subset(df, CDS==TRUE)
#   }
#   else {
#     mydata <- df
#   }
#   mydata$CDS_filter <- CDS_filter
#   
#   ### Summary heterozygosity data ###
#   # Summarize data per sample
#   mysummary <- mydata %>%
#     ddply(.(species, reference, method, replicate, CDS_filter), summarize,
#           heterozygosity_fraction = mean(heterozygosity),
#           heterozygosity_sd = sd(heterozygosity),
#           mean_coverage = mean(depth),
#           sd_coverage = sd(depth),
#           mean_quality = mean(QUAL),
#           sd_quality = sd(QUAL)
#     )
#   # Combine into summary_df (save outside of loop)
#   summary_df <- rbind(summary_df, mysummary)
#   
#   ### Binned across the genome heterozygosity data ###
#   for (bin_size in bin_sizes) {
#     # subset to HiC reference only
#     binned_df <- subset(mydata, reference == "HiC")
#     # split data into bins of specified size by dividing position by bin size and rounding up
#     binned_df$bin <- ceiling(binned_df$position/bin_size)
#     binned_df <- binned_df %>%
#         # calculate heterozygosity and other metrics per bin
#         ddply(.(species, reference, method, replicate, CDS_filter, scaffold, bin), summarize,
#               sites_count = length(bin), # total number of genotyped sites per bin
#               heterozygosity_count = sum(heterozygosity),
#               heterozygosity_fraction = mean(heterozygosity),
#               heterozygosity_sd = sd(heterozygosity),
#               mean_minor_allele_freq = mean(minor_allele_frequency),
#               mean_coverage = mean(depth),
#               sd_coverage = sd(depth),
#               mean_quality = mean(QUAL),
#               sd_quality = sd(QUAL)
#         )
#       # Save binned heterozygosity data as csv (one per CDS filter & bin size combo)
#       fileName <- paste0(summary_files_dir,'/tardigrade_HiC_heterozygosity_all_samples_CDS_filter_',CDS_filter,'_',bin_size/1000,'kb_bins.csv')
#       write.csv(binned_df, fileName, row.names = FALSE)
#       
#       message(paste0("Binned data saved as ", fileName))
#   }
# }
# 
# # Save summary heterozygosity data as csv (CDS filter ON/OFF both included per sample)
# fileName <- paste0(summary_files_dir,"/heterozygosity_summary_data.csv")
# write.csv(summary_df, fileName, row.names = FALSE)
# 
# message(paste0("Summary data saved as ", fileName))


## With four-fold-degenerate filter, no CDS filter, NCBI reference only, summary per sample only
ffd_summary_df <- df %>%
  subset(four_fold_degenerate == TRUE & reference == "NCBI") %>%
  ddply(.(species, reference, method, replicate), summarize,
        heterozygosity_fraction = mean(heterozygosity),
        heterozygosity_sd = sd(heterozygosity),
        mean_coverage = mean(depth),
        sd_coverage = sd(depth),
        mean_quality = mean(QUAL),
        sd_quality = sd(QUAL)
  )

# Save summary heterozygosity data as csv
fileName <- paste0(summary_files_dir,"/four_fold_degenerate_heterozygosity_summary_data.csv")
write.csv(ffd_summary_df, fileName, row.names = FALSE)

message(paste0("Four fold degenerate site summary data saved as ", fileName))
```
