## Supplementary material for "Modified meiosis in the tardigrade *Hypsibius exemplaris* maintains heterozygosity across the genome": File S1: Compile_RNA_Expression_Data.R

```
library(tximport)
library(readr)
library(dplyr)
library(tidyr)
library(tibble)
library(stringr)
library(purrr)

###############################################################################
# User Settings (change if needed)
###############################################################################

## Set working directory
setwd("/work/users/a/d/adricoke/single_tardigrade_seq_analysis/rna_expression_analysis_results")
main_dir = getwd()

###############################################################################
# Find File locations and treatments
###############################################################################

## Identify sample files
data_files <- dir(main_dir, recursive=T, include.dirs=T, pattern="quant.sf", full.names=T)
data_files

## Extract info from sample file paths
sample_names = NULL # all info from below, for pipeline usage
citations = NULL
sample_descriptions = NULL
replicates = NULL
for (file in data_files) {
  temp <- basename(dirname(file))
  sample_names <- c(sample_names, temp)
  citations <- c(citations, unlist(strsplit(temp, "_"))[1])
  sample_descriptions <- c(sample_descriptions, unlist(strsplit(temp, "_"))[2])
  replicates <- c(replicates, unlist(strsplit(temp, "_"))[3])
}
sample_names
citations
sample_descriptions
replicates

## read in transcript->gene map
tx2gene <- read.csv("tx2gene.csv", stringsAsFactors = FALSE)
colnames(tx2gene) <- c("transcript_id","gene_id")
head(tx2gene)

###############################################################################
# Use tximport to analyze data
###############################################################################

## Build files vector for tximport (names = sample_names)
# This mirrors tximport’s expected input: a named vector of quant.sf paths
names(data_files) <- sample_names
files <- data_files

## tximport to aggregate Salmon transcripts -> genes (TPM)
txi <- tximport(files, type = "salmon", tx2gene = tx2gene)

# ---- Strip "gene-" prefix from gene IDs (applies to all outputs) ----
clean_ids <- sub("^gene-", "", rownames(txi$abundance))
rownames(txi$abundance) <- clean_ids

gene_tpm_mat <- txi$abundance  # genes x samples, TPM
# head(gene_tpm_mat)
gene_tpm_df  <- data.frame(gene_id = rownames(gene_tpm_mat), gene_tpm_mat,
                           check.names = FALSE)
# head(gene_tpm_df)

## Per-sample rankings: rank, percentile, z-score (TPM-based)
epsilon <- 1e-6
gene_tpm_long <- gene_tpm_df %>%
  tidyr::pivot_longer(cols = -gene_id, names_to = "sample", values_to = "TPM") %>%
  # attach parsed metadata (same order as sample_names)
  dplyr::left_join(
    tibble(sample = sample_names,
           citation = citations,
           description = sample_descriptions,
           replicate = replicates),
    by = "sample"
  ) %>%
  dplyr::group_by(sample) %>%
  dplyr::mutate(
    rank       = rank(-TPM, ties.method = "min"),             # 1 = highest TPM
    percentile = 100 * (1 - (rank - 1) / dplyr::n()),         # 100 = top
    logTPM     = log10(TPM + epsilon),
    zscore     = (logTPM - mean(logTPM)) / sd(logTPM)
  ) %>%
  dplyr::ungroup()
# head(gene_tpm_long)

## Overall rankings across samples (mean/median/geometric mean TPM)
gene_stats <- gene_tpm_long %>%
  dplyr::group_by(gene_id) %>%
  dplyr::summarize(
    mean_TPM          = mean(TPM),
    median_TPM        = median(TPM),
    geo_mean_TPM      = exp(mean(log(TPM + epsilon))) - epsilon,
    max_TPM           = max(TPM),
    mean_rank         = mean(rank),
    median_rank       = median(rank),
    samples_expressed = sum(TPM > 0),
    n_samples         = dplyr::n_distinct(sample),
    .groups = "drop"
  ) %>%
  dplyr::mutate(
    rank_meanTPM     = rank(-mean_TPM, ties.method = "min"),
    rank_medianTPM   = rank(-median_TPM, ties.method = "min"),
    rank_geoMeanTPM  = rank(-geo_mean_TPM, ties.method = "min"),
    rank_byAvgRank   = rank(mean_rank, ties.method = "min")   # lower mean_rank = higher expression
  )
# head(gene_stats)

###############################################################################
# Save outputs as csv's
###############################################################################

## Write outputs to working directory
write.csv(gene_tpm_long, file.path(main_dir, "gene_tpm_ranks_per_sample.csv"),
          row.names = FALSE)
write.csv(gene_stats, file.path(main_dir, "gene_expression_rank_overall.csv"),
          row.names = FALSE)
write.csv(gene_tpm_df, file.path(main_dir, "gene_tpm_matrix.csv"),
          row.names = FALSE)

# head(gene_tpm_long)
# head(gene_stats)
# head(gene_tpm_df)
```
