## Supplementary material for "Modified meiosis in the tardigrade *Hypsibius exemplaris* maintains heterozygosity across the genome": File S1: Compile_Variant_Summary_Data.R

```
library(tidyverse)

###############################################################################
# User Settings (change if needed)
###############################################################################

## Set working directory
setwd("/work/users/a/d/adricoke/single_tardigrade_seq_analysis/NCBI_genome_results")
main_dir = getwd()

###############################################################################
# Step 1. Find File locations and treatments
###############################################################################

## Identify files
# heterozygous_variants.csv includes only filtered variants (QUAL>=30, depth>=40, heterozygous-only sites)
data_files <- dir(main_dir, recursive=T, include.dirs=T, pattern="heterozygous_variants.csv", full.names=T)
data_files

## Extract info from file path
# assumes file path structure is: .../species_replicate/heterozygous_variants.csv
species = NULL
replicates = NULL
for (file in data_files) {
  temp <- unlist(strsplit(file, "/heterozygous_variants.csv"))
  temp2 <- unlist(strsplit(temp[1], "/"))[9]
  species <- c(species, unlist(strsplit(temp2, "_rep"))[1])
  replicates <- c(replicates, ifelse(is.na(unlist(strsplit(temp2, "_rep"))[2]), 1, unlist(strsplit(temp2, "_rep"))[2]))
}
species
replicates

###############################################################################
# Step 2. Read in, Organize, & Combine Data
###############################################################################

# List of subfields in each SnpEff annotation (16 fields per spec)
snpeff_fields <- c(
  "Allele", "Annotation", "Annotation_Impact", "Gene_Name", "Gene_ID",
  "Feature_Type", "Feature_ID", "Transcript_BioType", "Rank", 
  "HGVS_c", "HGVS_p", "cDNA_pos", "CDS_pos", "AA_pos", 
  "Distance", "Errors_Warnings_Info"
)

df <- as.data.frame(NULL)
for (i in 1:length(data_files)) {
# for (i in 3:5) { # for testing, H.e. bulk samples only
  
  ## Read in Data ##
  mydata = readr::read_csv(data_files[i], col_names=FALSE)
  
  ## Organize Data ##
  
  # Identify number of columns
  ncols <- ncol(mydata)
  
  # Name the fixed columns (first 8 are known)
  fixed_cols <- c("scaffold","position","REF","ALT","QUAL","depth","REF_depth","ALT_depth")
  
  # Name the remaining columns generically (any number of SnpEff columns)
  snpeff_cols_raw <- paste0("SnpEff", seq_len(ncols - length(fixed_cols)))
  
  # Apply names
  colnames(mydata) <- c(fixed_cols, snpeff_cols_raw)
  
  # Reshape from wide to long (one annotation per row)
  mydata <- mydata %>%
    pivot_longer(
      cols = starts_with("SnpEff"), 
      names_to = "SnpEff_num", 
      values_to = "SnpEff_raw",
      values_drop_na = TRUE
    ) %>%
    filter(!is.na(SnpEff_raw) & SnpEff_raw != "") %>%
    # Split each SnpEff_raw into its 16 fields
    separate(SnpEff_raw, into = snpeff_fields, sep = "\\|", fill = "right")
  
  # Add additional columns for species + replicate
  mydata$species <- species[i]
  mydata$replicate <- replicates[i]
  
  # Select and rename desired fields
  mydata <- mydata %>%
    transmute(
    species, replicate,
    scaffold, position, REF, ALT, QUAL, depth, REF_depth, ALT_depth,
    Annotation,
    Annotation_Impact,
    Gene_Name,
    Gene_ID,
    AA_change = HGVS_p,
    AA_pos
  )
  
  ## Add data to df ##
  df <- rbind(df, mydata)
}

head(df)

### (Optional) Save Combined Data ###

# fileName <- "SnpEff_variants.csv"
# write.csv(df, fileName, row.names = FALSE)


###############################################################################
# Step 3. Summarize Data: Count Variants
###############################################################################

### Count variants by category ###
variant_type_summary <- df %>%
  count(species, replicate, Annotation, Annotation_Impact, sort = FALSE)
# head(variant_type_summary)

## Save Summary Data
fileName <- "SnpEff_variant_counts.csv"
write.csv(variant_type_summary, fileName, row.names = FALSE)


### Count variants per gene, per sample ###
per_gene_summary <- df %>%
  filter(Annotation != "intergenic_region") %>%
  count(species, replicate, Gene_ID, sort = FALSE)
# head(per_gene_summary)

## Save Summary Data
fileName <- "SnpEff_per_gene_het_variant_counts.csv"
write.csv(per_gene_summary, fileName, row.names = FALSE)

###############################################################################
# Step 4. Find Tardigrade Variants that are Shared Across Samples
###############################################################################

### 1/3: Filter to only variants shared across at least 2 single-tardigrade replicates ###

shared_variants <- df %>%
  subset(species == "Hypsibius_exemplaris") %>%
  # identify which variant sites are seen in all 4 replicates
  distinct(species, replicate, scaffold, position, REF, ALT) %>%
  group_by(species, scaffold, position, REF, ALT) %>%
  summarise(n_reps = n_distinct(replicate), .groups = "drop") %>%
  filter(n_reps >= 2) %>%
  # join back to original df to recover annotation info, etc.
  left_join(df, by = c("species", "scaffold", "position", "REF", "ALT")) %>%
  # keep one representative row per variant
  distinct(species, scaffold, position, REF, ALT, .keep_all = TRUE) %>%
  select(-replicate)                      # drop replicate info

# head(shared_variants)

### Save Data ###

write.csv(shared_variants, "He_variants_shared_in_>=2_reps.csv", row.names = FALSE)

###############################################################################

### 2/3: Filter to only variants shared across all 3 BULK tardigrade samples ###

bulk_shared_variants <- df %>%
  subset(species == "Hypsibius_exemplaris_bulk1" | species == "Hypsibius_exemplaris_bulk2" | species == "Hypsibius_exemplaris_bulk3") %>%
  # identify which variant sites are seen in all 3 "species" (samples)
  distinct(species, scaffold, position, REF, ALT) %>%
  group_by(scaffold, position, REF, ALT) %>%
  summarise(n_reps = n_distinct(species), .groups = "drop") %>%
  filter(n_reps == 3) %>%
  # join back to original df to recover annotation info, etc.
  left_join(df, by = c("scaffold", "position", "REF", "ALT")) %>%
  # keep one representative row per variant
  distinct(scaffold, position, REF, ALT, .keep_all = TRUE) %>%
  select(-replicate)                      # drop replicate info

# head(shared_variants)

### Save Data ###

write.csv(bulk_shared_variants, "He_variants_shared_in_all_bulk_seq_samples.csv", row.names = FALSE)

###############################################################################

### Filter to only HIGH-IMPACT variants shared across all 3 BULK tardigrade samples AND one single-tardigrade replicate ###

# Make a subset of single-tardigrade data
single <- df %>% 
  filter(species == "Hypsibius_exemplaris")

# Add unique keys to each dataset
bulk_keyed <- bulk_shared_variants %>%
  mutate(var_id = paste(scaffold, position, REF, ALT, sep = "_"))

single_keyed <- single %>%
  mutate(var_id = paste(scaffold, position, REF, ALT, sep = "_"))

# Pull the set of variant IDs found in at least one single replicate
single_unique_ids <- unique(single_keyed$var_id)

bulk_overlap_single <- bulk_keyed %>%
  filter(var_id %in% single_unique_ids)

# subset to only high impact variants (nonsense, frameshift...)
bulk_overlap_single_high_impact <- bulk_overlap_single %>%
  subset(Annotation_Impact == "HIGH")

### Save Data ###

fileName <- "select_high_impact_variants.csv"
write.csv(bulk_overlap_single_high_impact, fileName, row.names = FALSE)
```
