## Supplementary material for "Modified meiosis in the tardigrade *Hypsibius exemplaris* maintains heterozygosity across the genome": File S1: High_Impact_Variants_ASE_Analysis.R

```
library(data.table)

###############################################################################
# Step 1. Specify File Locations and Read in Data
###############################################################################

### File locations ###

# I downloaded files locally, per sample, before plotting

setwd("C:/Users/cburch/Dropbox/addie/tardigrades")

### ASEQ Results ###
ASEQ_variant_files <- dir(getwd(), recursive=T, include.dirs=T, pattern=".ASE.ASEQ", full.names=T)
ASEQ_variant_files

ASEQ_variants <- data.table()
for (f in ASEQ_variant_files) {
  mydata <- fread(f)
  mydata$sample <- sub(".*(SRR[0-9]+)\\..*", "\\1", f)
  ASEQ_variants <- rbind(ASEQ_variants, mydata)
}
head(ASEQ_variants)

### SnpEff-identified variants of interest ###
snpEff_variant_file <- "select_high_impact_variants.csv"
snpEff_variants <- fread(snpEff_variant_file)
head(snpEff_variants)

length(unique(snpEff_variants$Gene_ID))

### GENE INFORMATION ###
genes_stats <- fread("genes_statistics.csv") # for all tested genes, fraction of SNPs with ASE, per sample
head(genes_stats)


#######################################
# COMBINE INFORMATION FROM THE THREE DATA FILES
#######################################

#remove indels
ASEQ_variants <- ASEQ_variants[nchar(ref) == 1 & nchar(alt) == 1]

# add a "gene" column that identifies variants within genes by matching chr and pos between start and end
# note that some positions map to multiple genes.  This code just grabs the first match.
ASEQ_variants[, gene := genes_stats[.SD, on = .(chr, start <= pos, end >= pos), mult = "first", x.gene]]

# create columns to hold counts of the REF and ALT base calls
nuc_mat <- as.matrix(ASEQ_variants[, c("A", "C", "G", "T")])
ASEQ_variants$ref_count <- nuc_mat[cbind(seq_len(nrow(ASEQ_variants)),
                                         match(ASEQ_variants$ref, c("A", "C", "G", "T")))]

ASEQ_variants$alt_count <- nuc_mat[cbind(seq_len(nrow(ASEQ_variants)),
                                         match(ASEQ_variants$alt, c("A", "C", "G", "T")))]

# create columns to hold the minor and ALT allele frequencies
ASEQ_variants[, minor_AF := pmin(ref_count, alt_count) / (ref_count + alt_count)]
ASEQ_variants[, alt_AF := alt_count / (ref_count + alt_count)]

# try to capture the alignment bias that reads with REF alleles are more likely to
# to align than reads with ALT alleles.  Use only variant sites in genes.
mean_alt_AF <- mean(ASEQ_variants[!is.na(gene)]$alt_AF)
mean_alt_AF
# 0.4709173

ASEQ_variants[, `:=`(
  p_value = pbinom(alt_count, cov, mean_alt_AF)
)]

head(ASEQ_variants)

ASEQ_variants_in_genes <- ASEQ_variants[!is.na(gene)]

# Is the ALT allele the minor allele?
ASEQ_variants_in_genes$is_minor <- as.numeric(ASEQ_variants_in_genes$minor_AF==ASEQ_variants_in_genes$alt_AF)

z <- ASEQ_variants_in_genes[, .(
  n_samples = .N,
  median_AF = median(alt_AF),
  median_p = median(p_value),
  n_minor = sum(is_minor)
), by = .(gene,pos)]

nrow(z)
#11705

#use all variants that passed the quality filters in all 3 samples to determine a significance threshold
z2 <- z[n_samples==3]
n_tests <- nrow(z2)
n_tests
#9258

x <- 0
repeat {
  x <- x + 1
  if (nrow(z2[n_minor == 3 & (median_p < 0.05 / (n_tests-x))]) < x) {
    print(x - 1)
    break
  }
}
n_sig_tests <- x-1
n_sig_tests
#607

p_cutoff <- 0.05 / (n_tests - n_sig_tests)
p_cutoff
# 5.779679e-06


# variant sites we can assess = in all 3 samples and in genes
z2 <- z[n_samples==3 & !(gene %in% snpEff_variants$Gene_ID)]
nrow(z2)
#9001


minor_ase <- nrow(z2[n_minor == 3 & (median_p < p_cutoff)])
minor_ase
#593

#fraction of alleles with significantly reduced expression
minor_ase/nrow(z2)
#0.0659


##############################################
high_impact_genes <- ASEQ_variants[gene %in% snpEff_variants$Gene_Name]

high_impact_variants <- high_impact_genes[
  snpEff_variants,
  on = .(chr = scaffold, pos = position),
  nomatch = 0
]

high_impact_variants$is_minor <- as.numeric(high_impact_variants$minor_AF==high_impact_variants$alt_AF)


z <- high_impact_variants[, .(
  n_samples = .N,
  n_minor = sum(is_minor),
  median_coverage = median(cov),
  median_ALT_allele_frequency = median(alt_AF),
  median_p = median(p_value)
), by = .(gene,chr,pos)]

head(z)

z2 <- z[n_samples==3]
nrow(z2)
#48

fwrite(z2, file="high_impact_variants_ASE_data_and_statistics.csv")

minor_ase <- nrow(z2[n_minor == 3 & median_p < p_cutoff])
minor_ase
#3

minor_ase/nrow(z2)
#0.0625

#which genes?
ase_gene_names <- z2[n_minor == 3 & median_p < p_cutoff]$gene
snpEff_variants[Gene_Name %in% ase_gene_names]


dt <- high_impact_genes

pdf("genes_with_ase_minor_variants.pdf", width=9, height=18)
par(mfrow=c(6,3), mar=c(4,4,2,2))
for (g in ase_gene_names) {
  for (s in unique(dt$sample)) {
    geneset <- dt[gene==g & sample==s]
    if (nrow(geneset)>0) {
      
      geneset$p_value <- NA
      geneset$expected_minor_AF <- NA
      for (r in 1:nrow(geneset)) {
        geneset$p_value[r] <- pbinom(geneset$minor_AF[r]*geneset$cov[r],geneset$cov[r], 0.5)
        geneset$expected_minor_AF[r] <- expected_minor_AF(geneset$cov[r])
      }
      geneset$p_value[geneset$p_value < 1e-20] <- 1e-20
      
  

      plot(geneset$p_value ~ geneset$minor_AF, 
           main=paste(g,sub(".*(..)\\..*$", "\\1", s)),
           xlab="minor_AF",
           ylab="binomial p-value",
           log="y",
           ylim=c(min(geneset$p_value),1),
           xlim=c(0,0.5),
           col=as.factor(geneset$pos)
      )
      abline(h = 0.05, lty=2)
      abline(h = 0.05/nrow(geneset), lty=3)
      abline(v = mean(geneset$expected_minor_AF), lty=2, col="red")
      abline(v = mean(geneset$minor_AF), lty=1, col="red")
      legend("topleft", 
             legend=paste0(" mean cov=",round(mean(geneset$cov)),
                           "\n n_variants=",nrow(geneset)
             ),
             bty="n")
      temp <- high_impact_variants[gene==g]
      for (p in unique(temp$pos)) {
        if(geneset$minor_AF[geneset$pos==p]==geneset$alt_AF[geneset$pos==p]) {
          points(geneset$p_value[geneset$pos==p] ~ geneset$minor_AF[geneset$pos==p], pch=16)
        } else {
          points(geneset$p_value[geneset$pos==p] ~ geneset$minor_AF[geneset$pos==p], pch=13)
        }
      }
    } else {
      plot(c(1e-20,1) ~ c(0,0.5), type="n", 
           main=paste(g,sub(".*(..)\\..*$", "\\1", s)),
           xlab="minor_AF",
           ylab="binomial p-value",
           log="y",
           ylim=c(1e-20,1),
           xlim=c(0,0.5)
      )
    }
  }
}
dev.off()
```
