## Supplementary material for "Modified meiosis in the tardigrade *Hypsibius exemplaris* maintains heterozygosity across the genome": File S1: identify_synonymous_sites.R

```
# sbatch -t 02:00:00 --mem=3g --wrap="Rscript identify_synonymous_sites.R --transcript=1"

library(GenomicFeatures)
library(GenomicRanges)
library(Biostrings)
library(Rsamtools)
library(data.table)
library(optparse)

# ---- inputs ----
gff_file <- "/work/users/a/d/adricoke/single_tardigrade_seq_analysis/refs/Hypsibius_exemplaris/NCBI/GCA_002082055.1/genomic.gff"
fasta_file <- "/work/users/a/d/adricoke/single_tardigrade_seq_analysis/refs/Hypsibius_exemplaris/NCBI/GCA_002082055.1/GCA_002082055.1_nHd_3.1_genomic.fna"
output_folder <- "/work/users/c/b/cburch/addie/tardigrades/4_fold_degenerate_sites/"

#parse command line arguments
parser <- OptionParser()
parser <- add_option(parser, c("-t", "--transcript"), type="numeric",
                     default="", help="transcript number")
args= commandArgs(trailingOnly=TRUE)
a <- parse_args(parser, args)
#transcript_number <- 1
transcript_number <- a$transcript

# ---- build transcript database from GFF ----
txdb <- txdbmaker::makeTxDbFromGFF(gff_file, format = "gff3")

# CDS by transcript
cds_by_tx <- cdsBy(txdb, by = "tx", use.names = TRUE)

# ---- load reference genome ----
fa <- FaFile(fasta_file)
open(fa)

# helper: get CDS sequence in transcript orientation
get_tx_seq <- function(gr) {
  seqs <- getSeq(fa, gr)
  if (as.character(unique(strand(gr))) == "-") {
    seqs <- reverseComplement(seqs)
  }
  do.call(xscat, as.list(seqs))
}

# helper: is this codon position 4-fold degenerate?
is_fourfold_site <- function(codon, pos) {
  codon <- toupper(as.character(codon))
  if (nchar(codon) != 3 || grepl("[^ACGT]", codon)) return(FALSE)
  
  aa0 <- as.character(translate(DNAString(codon)))
  bases <- c("A", "C", "G", "T")
  orig <- substr(codon, pos, pos)
  alts <- setdiff(bases, orig)
  
  for (alt in alts) {
    mut <- codon
    substr(mut, pos, pos) <- alt
    aa1 <- as.character(translate(DNAString(mut)))
    if (aa1 != aa0) return(FALSE)
  }
  TRUE
}

# ---- collect 4-fold sites ----
rows <- data.table()
for (tx in names(cds_by_tx)[transcript_number:(transcript_number + 70)]) {
  gr <- cds_by_tx[[tx]]
  if (length(gr) == 0) next
  
  # extract CDS sequence in transcript order
  seq <- get_tx_seq(gr)
  
  # genomic positions in transcript order
  tx_pos <- unlist(lapply(seq_along(gr), function(i) {
    rng <- gr[i]
    if (as.character(strand(rng)) == "+") {
      seq(start(rng), end(rng))
    } else {
      seq(end(rng), start(rng))
    }
  }))
  
  # keep only complete codons
  n <- min(length(seq), length(tx_pos))
  n3 <- floor(n / 3) * 3
  if (n3 < 3) next
  
  seq <- subseq(seq, 1, n3)
  tx_pos <- tx_pos[1:n3]
  
  # scan codons
  for (j in seq(1, n3, by = 3)) {
    codon <- subseq(seq, j, j + 2)
    codon_chr <- as.character(codon)
    
    #only 3rd codon positions are 4-fold degenerate
    pos <- 3
    if (!is_fourfold_site(codon_chr, pos)) next
    
    genomic_pos <- tx_pos[j + pos - 1]
    chr <- as.character(seqnames(gr)[1])
  
    result <- data.table(
      chr = chr,
      start = as.integer(genomic_pos - 1),
      end = as.integer(genomic_pos),
      tx = tx,
      codon_chr = codon_chr,
      pos = pos
    )
    
    rows <- rbind(rows, result)
    
  }
}

close(fa)

# ---- write BED ----
setorder(rows, chr, start, end)

fwrite(
  rows[, .(chr, start, end, tx, codon_chr, pos)],
  paste0(output_folder,tx,'.bed'),
  sep = "\t",
  quote = FALSE,
  col.names = FALSE
)
```
