## Supplementary material for "Modified meiosis in the tardigrade *Hypsibius exemplaris* maintains heterozygosity across the genome": File S1: Make_Pedigrees.R

```
library(tidyverse)

setwd("C:/Users/addie/OneDrive - University of North Carolina at Chapel Hill/LAB/DATA/BUSCO_allele_inheritance/BUSCO_allele_inheritance_expt_plotting")

plots_dir <- "C:/Users/addie/OneDrive - University of North Carolina at Chapel Hill/LAB/tardigrade_heterozygosity/He_FIGURE_PLOTS"

###############################################################################
# Read and organize data
###############################################################################

## Technical replicate 1 (majority of data)
raw <- read.csv("20230929_BUSCO_allele_REdigest_genotyping.csv")

# Long format for selected loci
geno_long <- raw %>%
  pivot_longer(
    cols = c(B06, B01, B03),
    names_to = "locus",
    values_to = "genotype"
  ) %>%
  select(Generation, Individual, locus, genotype) %>%
  mutate(
    genotype = str_trim(genotype),
    genotype = na_if(genotype, "")
  )

## Technical replicate 2
tr2_raw <- read.csv("20260107_REdigest_technical_replicate_2.csv")

# pivot to long format (allele presence/absence)
tr2_long <- tr2_raw %>%
  mutate(
    TR2_genotype = str_trim(TR2_genotype),
    TR2_genotype = na_if(TR2_genotype, "")
  ) %>%
  mutate(
    allele1_TR2 = TR2_genotype %in% c("het", "hom1"),
    allele2_TR2 = TR2_genotype %in% c("het", "hom2")
  ) %>%
  select(Individual, Locus, allele1_TR2, allele2_TR2)

# Allele presence/absence for tech rep 1
geno_alleles <- geno_long %>%
  mutate(
    allele1 = genotype %in% c("het", "hom1"),
    allele2 = genotype %in% c("het", "hom2"),
    inconclusive = is.na(genotype) | genotype == "na",
    family = str_extract(Individual, "^[A-Z]"),
    generation = nchar(str_remove(Individual, "^[A-Z]")),
    parent = if_else(generation == 0, NA_character_, str_sub(Individual, 1, -2))
  ) %>%
  filter(family %in% c("A", "B", "C"),
         generation %in% c(0, 1, 2)) %>%
  # join tech rep 2 data
  left_join(
    tr2_long,
    by = c("Individual", "locus" = "Locus")
  ) %>%
  mutate(
    allele1_TR2 = replace_na(allele1_TR2, FALSE),
    allele2_TR2 = replace_na(allele2_TR2, FALSE)
  )

###############################################################################
# Prepare positions
###############################################################################

# All individuals table
all_individuals <- raw %>%
  mutate(
    family = str_extract(Individual, "^[A-Z]"),
    generation = nchar(str_remove(Individual, "^[A-Z]")),
    parent = if_else(generation == 0, NA_character_, str_sub(Individual, 1, -2))
  ) %>%
  select(Individual, family, generation, parent) %>%
  filter(family %in% c("A", "B", "C"),
         generation %in% c(0, 1, 2))

# Determine which individuals to keep:
# keep if any genotype detected OR has children
individuals_to_keep <- geno_alleles %>%
  group_by(Individual) %>%
  summarize(has_data = any(genotype!='na'), .groups = "drop") %>%
  left_join(
    all_individuals %>% mutate(has_children = Individual %in% parent) %>% select(Individual, has_children),
    by = "Individual"
  ) %>%
  filter(has_data | has_children)

# Filter all_individuals to only plotted individuals
positions <- all_individuals %>%
  semi_join(individuals_to_keep, by = "Individual") %>%
  distinct(Individual, family, generation, parent)

# Initialize y for leaves (no children)
positions <- positions %>%
  mutate(has_children = Individual %in% parent) %>%
  group_by(family) %>%
  mutate(
    y = if_else(!has_children, row_number(), NA_real_)
  ) %>%
  ungroup()

# Center parents over children
repeat {
  updated <- FALSE
  
  for (g in sort(unique(positions$generation), decreasing = TRUE)) {
    parents <- positions %>% filter(generation == g - 1) %>% pull(Individual)
    
    for (p in parents) {
      children_y <- positions %>% filter(parent == p) %>% pull(y)
      if (length(children_y) > 0 && any(!is.na(children_y))) {
        new_y <- mean(children_y, na.rm = TRUE)
        if (is.na(positions$y[positions$Individual == p])) {
          positions$y[positions$Individual == p] <- new_y
          updated <- TRUE
        }
      }
    }
  }
  
  if (!updated) break
}

# Small vertical separation within generation to avoid overlap
positions <- positions %>%
  group_by(family, generation) %>%
  mutate(y = y + row_number() * 0.02) %>%
  ungroup()

# Join positions back to geno_alleles
geno_alleles <- geno_alleles %>%
  semi_join(positions, by = "Individual") %>%  # only keep plotted individuals
  left_join(positions %>% select(Individual, y), by = "Individual")

###############################################################################
# Prepare 6-quadrant squares per individual
###############################################################################

cbPalette <- c("#999999", "#56B4E9", "#E69F00", "#CC79A7", "#009E73")

# define colors per locus
locus_colors <- c(
  B01 = "#E69F00",  
  B03 = "#56B4E9",  
  B06 = "#CC79A7"   
)

# total size of individual square (controls overall visual size)
square_size <- 0.2

# factor for scaling width and height independently
y_factor <- 0.52   # larger = smaller height

allele_quadrants <- geno_alleles %>%
  pivot_longer(
    cols = c(allele1, allele2),
    names_to = "allele",
    values_to = "detected_initial"
  ) %>%
  mutate(
    detected_TR2 = if_else(
      allele == "allele1", allele1_TR2, allele2_TR2
    ),
    detected_any = detected_initial | detected_TR2,
    detected_TR2_only = !detected_initial & detected_TR2
  ) %>%
  filter(detected_any | TRUE) %>%  # keep all quadrants (white squares matter)
  mutate(
    locus_num = as.numeric(factor(locus)),
    allele_num = if_else(allele == "allele1", 1, 2),
    x_min_indiv = generation - square_size/2,
    x_max_indiv = generation + square_size/2
  ) %>%
  mutate(
    xmin = x_min_indiv + (allele_num - 1) * (x_max_indiv - x_min_indiv)/2,
    xmax = xmin + (x_max_indiv - x_min_indiv)/2,
    ymin = y - (square_size / y_factor) +
      (3 - locus_num) * (2 * square_size / (3 * y_factor)),
    ymax = ymin + (2 * square_size / (3 * y_factor))
  ) %>%
  mutate(
    fill_state = case_when(
      detected_initial            ~ "initial",
      detected_TR2_only           ~ "TR2_only",
      TRUE                        ~ "none"
    )
  )

# Edges for inheritance lines
edges <- positions %>%
  filter(!is.na(parent)) %>%
  left_join(positions %>% select(Individual, parent_y = y),
            by = c("parent" = "Individual"))

###############################################################################
# Plotting
###############################################################################

# pick the founder (generation 1 or 0 depending on your numbering) per family
family_labels <- positions %>%
  group_by(family) %>%
  filter(generation == min(generation)) %>%  # first generation
  slice(1) %>%                               # in case multiple founders
  ungroup()


ggplot() +
  # edges
  geom_segment(
    data = edges,
    aes(x = generation - 1, y = parent_y,
        xend = generation, yend = y),
    linewidth = 0.4,
    color = "grey60"
  ) +
  # 6-quadrant squares
  geom_rect(
    data = allele_quadrants,
    aes(xmin = xmin, xmax = xmax,
        ymin = ymin, ymax = ymax,
        # use fill = fill_state for black/white, else fill = fill_color to color by locus
        fill = fill_state),
    color = "black",
    linewidth = 0.4
  ) +
  # label the founder of each family
  geom_text(
    data = family_labels,
    aes(x = generation, y = y + 1, label = family),
    size = 4
  ) +
  # aesthetics
  # black/white OR color by locus
  scale_fill_manual(
    name = "Allele detection",
    breaks = c("initial", "TR2_only", "none"),
    values = c(
      "initial"   = "black",
      "TR2_only"  = "grey50",
      "none"      = "white"
    ),
    labels = c(
      "Detected in initial assay",
      "Detected after replication",
      "Not detected"
    )
  ) +
  # scale_fill_manual(name = "Locus",
  #                   values = c(locus_colors, "white" = "white")) +
  facet_wrap(~ family, scales = "fixed", ncol = 3) +
  theme_void() +
  theme(
    panel.spacing = unit(0.5, "lines"),
    legend.position = c(0.99, 0.95),
    legend.justification = c("right", "top"),
    strip.text = element_blank(),
    legend.key.size = unit(0.8, "lines"),
  )

plot_title <- "Pedigrees"
ggsave(filename=paste0(plots_dir,"/",plot_title,".pdf"), width=6, height=5)
ggsave(filename=paste0(plots_dir,"/",plot_title,".png"), width=6, height=5, dpi=300)
```
