## Supplementary material for "Modified meiosis in the tardigrade *Hypsibius exemplaris* maintains heterozygosity across the genome": File S1: OrthoDB_Loci_Heterozygosity_Plotting.R

```
library(nlme)
library(scales)
library(ggplot2)
library(grid)
library(gridExtra)
library(data.table)
library(tidyverse)

###############################################################################
# Specify File locations
###############################################################################

# summary files are downloaded locally before plotting
main_dir = "C:/Users/addie/OneDrive - University of North Carolina at Chapel Hill/LAB/DATA/BUSCO_allele_inheritance/BUSCO_allele_inheritance_expt_plotting"
setwd(main_dir)

plots_dir = "C:/Users/addie/OneDrive - University of North Carolina at Chapel Hill/LAB/tardigrade_heterozygosity/He_FIGURE_PLOTS"

###############################################################################
# Adjust settings for ALL plots
###############################################################################

# theme settings
theme_set(theme_classic())
theme_update(text=element_text(size=10),
             strip.background = element_blank(), # remove box around facet labels
             strip.text = element_text(face = "bold"), # and bold text
             legend.key.size = unit(0.1, "in") # adjust legend sizing
)

cbPalette <- c("#999999", "#56B4E9", "#E69F00", "#CC79A7", "#009E73")

###############################################################################
# Create dataframe manually from Benchling data
###############################################################################

# all loci fully encompass CDS of associated BUSCO gene
# SNP counts include single-nucleotide indels (only one of these is present, in BUSCO 2)

## create dataframe with manually entered data
BUSCO_loci <- data.frame(
  Locus = c("BUSCO 1", "BUSCO 2", "BUSCO 3"),
  Previous_name = c("BUSCO 1", "BUSCO 3", "BUSCO 6"),
  Gene_ID = c("BV898_04098", "BV898_05498", "BV898_09033"),
  Allele1_Length_w_primers = c(961, 1574, 2481),
  Allele1_Length_without_primers = c(917, 1531, 2436),
  Primer_pair_length = c(44, 43, 45),
  Allele2_Length_w_primers = c(961, 1573, 2421),
  Allele2_Length_without_primers = c(917, 1530, 2376),
  CDS_Length_without_non_SNP_Indels = c(734, 1181, 1385),
  SNPs_in_Locus = c(13, 17, 75),
  SNPs_in_CDS = c(8, 6, 36),
  SNPs_not_in_NCBI_ref = c(0, 0, 0),
  non_SNP_Indels = c(0, 0, 3),
  non_SNP_Indels_in_CDS = c(0, 0, 1),
  non_SNP_Indel_Lengths = c(NA, NA, "46 bp; 70 bp; 38 bp"),
  sum_non_SNP_Indel_Lengths = c(0, 0, 154),
  Indel_Notes = c(NA, "one non-CDS single-nucleotide indel", "70bp and 38bp indels are in allele 1; 46bp indel is in allele 2 and in CDS region; indels appear to be in repetitive regions")
)

## calculate length of each locus WITHOUT non-SNP indels (manually)
BUSCO_loci$Locus_Length_without_non_SNP_Indels <- c(
  BUSCO_loci$Allele1_Length_without_primers[1], # BUSCO 1 has no indels
  BUSCO_loci$Allele1_Length_without_primers[2], # BUSCO 2 has no indels
  BUSCO_loci$Allele2_Length_without_primers[3] - 46 # BUSCO 3
)
## also calculate length of each locus WITH non-SNP indels
BUSCO_loci$Locus_Length_with_non_SNP_Indels <- BUSCO_loci$Locus_Length_without_non_SNP_Indels + BUSCO_loci$sum_non_SNP_Indel_Lengths
BUSCO_loci$CDS_Length_with_non_SNP_Indels <- c(
  BUSCO_loci$CDS_Length[1], # BUSCO 1 has no indels
  BUSCO_loci$CDS_Length[2], # BUSCO 2 has no indels
  BUSCO_loci$CDS_Length[3] + 46# BUSCO 3
)
BUSCO_loci$polymorphisms_in_Locus = BUSCO_loci$SNPs_in_Locus + BUSCO_loci$non_SNP_Indels
BUSCO_loci$polymorphisms_in_CDS = BUSCO_loci$SNPs_in_CDS + BUSCO_loci$non_SNP_Indels_in_CDS

head(BUSCO_loci)

###############################################################################
# Organize dataframe for plotting
###############################################################################

## Pivot longer for plotting
BUSCO_loci_longer <- BUSCO_loci %>%
  select(Locus, SNPs_in_Locus, SNPs_in_CDS) %>%
  pivot_longer(cols = c("SNPs_in_Locus", "SNPs_in_CDS"),
               names_to = "Region",
               values_to = "SNP_Count") %>%
  mutate(Region = recode(Region,
                         "SNPs_in_Locus" = "Entire Locus",
                         "SNPs_in_CDS" = "CDS Only"))
# add lengths without non-SNP indels to longer dataframe
BUSCO_loci_longer <- BUSCO_loci_longer %>%
  mutate(Length_without_non_SNP_Indels = ifelse(
    Region == "Entire Locus",
    BUSCO_loci$Locus_Length_without_non_SNP_Indels[match(Locus, BUSCO_loci$Locus)],
    BUSCO_loci$CDS_Length_without_non_SNP_Indels[match(Locus, BUSCO_loci$Locus)]
  )) %>%
  mutate(Length_with_non_SNP_Indels = ifelse(
    Region == "Entire Locus",
    BUSCO_loci$Locus_Length_with_non_SNP_Indels[match(Locus, BUSCO_loci$Locus)],
    BUSCO_loci$CDS_Length_with_non_SNP_Indels[match(Locus, BUSCO_loci$Locus)]
  ))
# add polymorphisms counts (SNPs + non-SNP indels) to longer dataframe
BUSCO_loci_longer <- BUSCO_loci_longer %>%
  mutate(Polymorphism_Count = ifelse(
    Region == "Entire Locus",
    BUSCO_loci$polymorphisms_in_Locus[match(Locus, BUSCO_loci$Locus)],
    BUSCO_loci$polymorphisms_in_CDS[match(Locus, BUSCO_loci$Locus)]
  ))
# calculate SNPs per bp (heterozygosity)
BUSCO_loci_longer <- BUSCO_loci_longer %>%
  mutate(SNPs_per_bp_ignore_indels = SNP_Count / Length_without_non_SNP_Indels,
         SNPs_per_bp_include_indels = SNP_Count / Length_with_non_SNP_Indels,
         Polymorphisms_per_bp_include_indels = Polymorphism_Count / Length_with_non_SNP_Indels)

head(BUSCO_loci_longer)


###############################################################################

# add psuedo-locus that represents genome-wide averages
genome_wide_avg_longer <- data.frame(
  Locus = c("Genome-wide", "Genome-wide"),
  Region = c("Entire Locus", "CDS Only"),
  SNP_Count = c(NA, NA),
  Length_without_non_SNP_Indels = c(NA, NA),
  Length_with_non_SNP_Indels = c(NA, NA),
  # all 3 heterozygosity values below are the same:
  SNPs_per_bp_ignore_indels = c(1.40498414514137/100, 1.00481635759971/100),
  SNPs_per_bp_include_indels = c(1.40498414514137/100, 1.00481635759971/100),
  Polymorphism_Count = c(1.40498414514137/100, 1.00481635759971/100),
  Polymorphisms_per_bp_include_indels = c(1.40498414514137/100, 1.00481635759971/100)
)
head(genome_wide_avg_longer)

# combine with BUSCO loci data
BUSCO_loci_longer <- rbind(BUSCO_loci_longer, genome_wide_avg_longer)

# Re-label BUSCO loci
BUSCO_loci_longer$Locus <- recode(BUSCO_loci_longer$Locus,
                                   "BUSCO 1" = "1 (BV898_04098)",
                                   "BUSCO 2" = "2 (BV898_05498)",
                                   "BUSCO 3" = "3 (BV898_09033)")

###############################################################################
# Make Plots
###############################################################################
## Proportion of SNPs per length (not including large indels)

# v1: as percentage (SNPs per bp * 100 = % of heterozygous bases)

# flag pseudo-locus for distinct appearance in plot
BUSCO_loci_longer$is_genome_wide <- BUSCO_loci_longer$Locus == "Genome-wide"

ggplot(BUSCO_loci_longer, aes(x=Locus,
                              y=SNPs_per_bp_ignore_indels*100,
                              fill=Region,
                              alpha = is_genome_wide)) +
  geom_bar(stat="identity", position=position_dodge(),
           color = "grey20", aes(linetype = is_genome_wide)) +
  scale_fill_manual(values=c("#E69F00", "#CC79A7")) +
  scale_alpha_manual(values = c(`FALSE` = 1, `TRUE` = 0.55), guide = "none") +
  scale_linetype_manual(values = c(`FALSE` = "solid", `TRUE` = "dashed"),
                        guide = "none") +# save plot
  # label bars
  geom_text(
    aes(
      label = ifelse(
        is_genome_wide,
        scales::percent(SNPs_per_bp_ignore_indels, accuracy = 0.01),
        paste0(SNP_Count, " / ", Length_without_non_SNP_Indels)
      )
    ),
    position = position_dodge(width = 0.9),
    vjust = -0.25,
    size = 3
  ) +
  # expand top of plot slightly for bar labels
  scale_y_continuous(expand=expansion(mult=c(0, 0.1))) +
  theme(legend.position = c(0.15,0.75)) +
  labs(x="Locus", y="% of Heterozygous Bases", fill="Region")
plot_title <- 'BUSCO_Loci_Heterozygosity_Percentage_ignore_indels'
ggsave(paste0(plots_dir, "/", plot_title, '.png'), width=5, height=2.5)
ggsave(paste0(plots_dir, "/", plot_title, '.pdf'), width=5, height=2.5)


###############################################################################
## WHat is the mean heterozygosity across all 3 BUSCO loci?
mean_het_ignore_indels <- mean(BUSCO_loci_longer$SNPs_per_bp_ignore_indels[
  BUSCO_loci_longer$Region == "Entire Locus"])
mean_het_ignore_indels_percent <- mean_het_ignore_indels * 100
paste0("Mean PCR Product Heterozygosity: " ,mean_het_ignore_indels_percent, "% (ignoring indels)")

mean_het_CDS_ignore_indels <- mean(
  BUSCO_loci_longer$SNPs_per_bp_ignore_indels[
    BUSCO_loci_longer$Region == "CDS Only"])
mean_het_CDS_ignore_indels_percent <- mean_het_CDS_ignore_indels * 100
paste0("Mean CDS Heterozygosity: " ,mean_het_CDS_ignore_indels_percent, "% (ignoring indels)")
```
