## Supplementary material for "Modified meiosis in the tardigrade *Hypsibius exemplaris* maintains heterozygosity across the genome": File S1: Parse_OrthoDB_Results.ipynb

```
# %% [markdown]
OrthoDB search result for genes that are present in >90% and single-copy in >90% of all metazoa.

# %% [code]
import requests

# %% [code]
# list of orthoDB cluster id's
single_copy_metazoa = ["615596at33208", "273051at33208", "566116at33208", "137383at33208", "450643at33208", "123923at33208", "617020at33208", "386666at33208", "520612at33208", "496070at33208", "526839at33208", "463071at33208", "444231at33208", "413334at33208", "375016at33208", "453687at33208", "461193at33208", "423848at33208", "227087at33208", "648300at33208", "277935at33208", "312570at33208", "541421at33208", "529092at33208", "480700at33208", "581080at33208", "313057at33208", "531281at33208", "353768at33208", "516481at33208", "467106at33208", "642773at33208", "452574at33208", "312577at33208", "354829at33208", "355210at33208", "99987at33208", "274769at33208", "576633at33208", "99614at33208", "615998at33208", "152610at33208", "566916at33208", "548732at33208", "594131at33208", "274064at33208", "419914at33208", "359075at33208", "334331at33208", "389570at33208", "555538at33208", "57158at33208", "334336at33208", "302186at33208", "214788at33208", "367493at33208", "568787at33208", "474302at33208", "480735at33208", "543333at33208", "586719at33208", "581779at33208", "523206at33208", "580851at33208", "123605at33208", "255924at33208", "491333at33208", "400084at33208", "592872at33208", "344977at33208", "257339at33208", "577339at33208", "437653at33208", "540473at33208", "231700at33208", "595858at33208", "157138at33208", "534808at33208", "466602at33208", "224280at33208", "558385at33208", "643954at33208", "432155at33208", "124302at33208", "564230at33208", "503333at33208", "619415at33208", "564234at33208", "271297at33208", "435921at33208", "395147at33208", "465933at33208", "539600at33208", "289212at33208", "592087at33208", "514565at33208", "601768at33208", "435678at33208", "304388at33208", "291505at33208", "255786at33208", "466616at33208", "589901at33208", "467142at33208", "493913at33208", "108764at33208", "592094at33208", "412370at33208", "148067at33208", "565519at33208", "497015at33208", "569294at33208", "566860at33208", "370036at33208", "361100at33208", "244863at33208", "537718at33208", "439335at33208", "532930at33208", "291875at33208", "435939at33208", "440493at33208", "359582at33208", "449056at33208", "215654at33208", "604211at33208", "561248at33208", "540193at33208", "570646at33208", "372257at33208", "237498at33208", "416175at33208", "606048at33208", "522988at33208", "373040at33208", "239597at33208", "365691at33208", "266460at33208", "332870at33208", "591498at33208", "535806at33208", "250484at33208", "414571at33208", "332049at33208", "277291at33208", "435159at33208", "452208at33208", "156355at33208", "437924at33208", "533666at33208", "482691at33208", "189414at33208", "548444at33208", "161355at33208", "519777at33208", "571149at33208", "462897at33208", "270594at33208", "313091at33208", "551831at33208", "581475at33208", "592759at33208", "158845at33208", "564282at33208", "390402at33208", "188827at33208", "636300at33208", "289235at33208", "515528at33208", "571991at33208", "486793at33208", "143619at33208", "585232at33208", "471653at33208", "219856at33208", "318532at33208", "134468at33208", "501788at33208", "337178at33208", "608669at33208", "363634at33208", "436169at33208", "448514at33208", "573534at33208", "379636at33208", "589773at33208", "474387at33208", "509816at33208", "466942at33208", "556510at33208", "617522at33208", "628231at33208", "259581at33208", "117726at33208", "505117at33208", "563358at33208", "581089at33208", "563359at33208", "593097at33208", "302921at33208", "378033at33208", "363851at33208", "371297at33208", "540537at33208", "578582at33208", "124979at33208", "511781at33208", "608869at33208", "206858at33208", "520011at33208", "590221at33208", "518288at33208", "488635at33208", "197549at33208", "406052at33208", "495100at33208", "438169at33208", "338370at33208", "360753at33208", "359771at33208", "92367at33208", "393419at33208", "227979at33208", "534206at33208", "340559at33208", "451242at33208", "631196at33208", "577068at33208", "521343at33208", "351226at33208", "308610at33208", "315393at33208", "469367at33208", "263153at33208", "502584at33208", "447400at33208", "462325at33208", "624945at33208", "207599at33208", "477052at33208", "199603at33208", "510174at33208", "568839at33208", "359324at33208", "272605at33208", "562799at33208", "618128at33208", "508379at33208", "358036at33208", "364694at33208", "557834at33208", "344338at33208", "222489at33208", "367957at33208", "379682at33208", "268397at33208", "143997at33208", "532663at33208", "290029at33208", "323473at33208", "177858at33208", "606958at33208", "488474at33208", "545641at33208", "535524at33208", "529852at33208", "514892at33208", "333439at33208", "612256at33208", "623948at33208", "528608at33208", "539360at33208", "406927at33208", "283554at33208", "344346at33208", "537763at33208", "221288at33208", "478984at33208", "638541at33208", "349627at33208", "247729at33208", "194350at33208", "279992at33208", "504207at33208", "14776at33208", "575469at33208", "381303at33208", "457296at33208", "584333at33208", "395556at33208", "384113at33208", "267439at33208", "266808at33208", "212972at33208", "104525at33208", "556587at33208", "273989at33208", "285146at33208", "45428at33208", "614305at33208", "279631at33208", "203128at33208", "568170at33208", "396237at33208", "618339at33208", "433216at33208", "561878at33208", "387079at33208", "532693at33208", "516216at33208", "554010at33208", "590335at33208", "355739at33208", "401679at33208", "502691at33208", "569812at33208", "446000at33208", "616716at33208", "177718at33208", "422324at33208", "130256at33208", "532728at33208", "625314at33208", "640187at33208", "356972at33208", "387271at33208", "176261at33208", "464204at33208", "179485at33208", "148325at33208", "611979at33208", "603609at33208", "255657at33208", "295556at33208", "324781at33208", "576449at33208", "380583at33208", "578726at33208", "518569at33208", "389166at33208", "563084at33208", "359369at33208", "593401at33208", "211758at33208", "559732at33208", "438453at33208", "372707at33208", "279646at33208", "584373at33208", "391896at33208", "182834at33208", "632472at33208", "554815at33208", "441575at33208", "578734at33208", "464286at33208", "500416at33208", "351151at33208", "539043at33208", "450785at33208", "585656at33208", "446031at33208", "414603at33208", "317470at33208", "538442at33208", "570717at33208", "136652at33208", "465092at33208", "455941at33208", "622172at33208", "527986at33208", "111730at33208", "424392at33208", "545350at33208", "582823at33208", "367551at33208", "419502at33208", "289928at33208", "600462at33208", "327301at33208", "598405at33208", "334272at33208", "524322at33208", "107151at33208", "556925at33208", "597816at33208", "579029at33208", "541231at33208", "114954at33208", "424682at33208", "393291at33208", "206786at33208", "458974at33208", "577929at33208", "538913at33208", "188586at33208", "607638at33208", "585305at33208", "331112at33208", "511386at33208", "602587at33208", "533690at33208", "267612at33208", "517525at33208", "249486at33208", "639252at33208", "471787at33208", "560608at33208", "346056at33208", "569878at33208", "543124at33208", "114070at33208", "490961at33208", "247905at33208", "504292at33208", "235458at33208", "474110at33208", "549156at33208", "368565at33208", "603673at33208", "235411at33208", "490523at33208", "526673at33208", "487121at33208", "465703at33208", "465651at33208", "546305at33208", "531775at33208", "415618at33208", "485003at33208", "541260at33208", "559425at33208", "160378at33208", "510581at33208", "428754at33208", "63697at33208", "431718at33208", "239149at33208", "530431at33208", "305260at33208", "531789at33208", "535268at33208", "605084at33208", "238886at33208", "311758at33208", "440300at33208", "609899at33208", "192641at33208", "231157at33208", "405257at33208", "515640at33208", "454911at33208", "552201at33208", "300303at33208", "587273at33208", "157109at33208", "636423at33208", "313300at33208", "236965at33208", "617667at33208", "360379at33208", "284059at33208", "495605at33208", "315044at33208", "612760at33208", "319328at33208", "572754at33208", "376982at33208", "245482at33208", "642552at33208", "146671at33208", "468631at33208", "505248at33208", "203698at33208", "246241at33208", "164767at33208", "432522at33208", "555026at33208", "333800at33208", "539793at33208", "366124at33208", "395948at33208", "489359at33208", "561417at33208", "449189at33208", "385281at33208", "566551at33208", "167353at33208", "432287at33208", "303117at33208", "468359at33208", "460518at33208", "545063at33208", "311165at33208", "449473at33208", "623444at33208", "196950at33208", "435253at33208", "551620at33208", "611258at33208", "290413at33208", "563679at33208", "600823at33208", "346946at33208", "235430at33208", "294594at33208", "195242at33208", "407472at33208", "529272at33208", "340558at33208", "451608at33208", "551632at33208", "464987at33208", "587958at33208", "344611at33208", "618421at33208", "427789at33208", "500808at33208", "520147at33208", "611279at33208", "572016at33208", "111746at33208", "125779at33208", "424646at33208", "529286at33208", "558897at33208", "401196at33208", "442169at33208", "542217at33208", "585895at33208", "398607at33208", "178343at33208", "166097at33208", "615531at33208", "273367at33208", "365561at33208", "563488at33208", "421732at33208", "302279at33208", "361203at33208", "631767at33208", "184717at33208", "613522at33208", "324249at33208", "464312at33208", "262757at33208", "610541at33208", "226335at33208", "532406at33208", "586310at33208", "478189at33208", "600878at33208", "210614at33208", "410267at33208", "518635at33208", "473106at33208", "511575at33208", "245567at33208", "514724at33208", "380810at33208", "558552at33208", "623995at33208", "227453at33208", "581901at33208", "387117at33208", "332149at33208", "517311at33208", "135294at33208", "567650at33208", "531649at33208", "495311at33208", "465401at33208", "499366at33208", "179527at33208", "504691at33208", "601967at33208", "616155at33208", "502429at33208", "409120at33208", "186208at33208", "436007at33208", "477961at33208", "376375at33208", "459253at33208", "411664at33208", "285902at33208", "364329at33208", "500861at33208", "561970at33208", "517968at33208", "599913at33208", "514740at33208", "535664at33208", "415424at33208", "486743at33208", "205220at33208", "621808at33208", "369197at33208", "109109at33208", "494057at33208", "631354at33208", "428202at33208", "255049at33208", "487873at33208", "531134at33208", "582173at33208", "585078at33208", "533750at33208", "76852at33208", "210538at33208", "613580at33208", "608074at33208", "398202at33208", "486933at33208", "296113at33208", "404988at33208", "131863at33208", "609955at33208", "50756at33208", "508188at33208", "544849at33208", "641073at33208", "547341at33208", "158962at33208", "472352at33208", "389002at33208", "336647at33208", "576938at33208", "333182at33208", "609978at33208", "325183at33208", "591316at33208", "582265at33208", "223310at33208", "600204at33208", "544856at33208", "235149at33208", "451957at33208", "552949at33208", "620753at33208", "566026at33208", "354949at33208", "349064at33208", "423799at33208", "585726at33208", "311838at33208", "452499at33208", "341931at33208", "571944at33208", "565053at33208", "402058at33208", "239016at33208", "132087at33208", "526074at33208", "361265at33208", "153034at33208", "514698at33208", "444785at33208", "352854at33208", "511272at33208", "443305at33208", "319574at33208", "562561at33208", "321332at33208", "184612at33208", "483489at33208", "443928at33208", "582473at33208", "434068at33208", "601886at33208", "553585at33208", "545542at33208", "559501at33208", "493148at33208", "393895at33208", "559883at33208", "342340at33208", "572434at33208", "260717at33208", "411515at33208", "556039at33208", "562918at33208", "366814at33208", "259183at33208", "545481at33208", "486993at33208", "399439at33208", "115878at33208", "637971at33208", "516716at33208", "582951at33208", "519303at33208", "503417at33208", "558933at33208", "631811at33208", "557694at33208", "200997at33208", "534757at33208", "335372at33208", "641257at33208", "618986at33208", "228301at33208", "535367at33208", "434863at33208", "513998at33208", "625769at33208", "618516at33208", "578959at33208", "348449at33208", "590049at33208", "576997at33208", "620546at33208", "417629at33208", "594333at33208", "568691at33208", "498535at33208", "353286at33208", "525478at33208", "515058at33208", "127766at33208", "595404at33208", "491252at33208", "525888at33208", "592612at33208", "531512at33208", "281304at33208", "497925at33208", "548327at33208", "239034at33208", "601242at33208", "585313at33208", "617398at33208", "495802at33208", "443659at33208", "630609at33208", "597958at33208", "169864at33208", "562382at33208", "251577at33208", "189127at33208", "422751at33208", "587399at33208", "544546at33208", "583132at33208", "273543at33208", "364383at33208", "382750at33208", "479155at33208", "591391at33208", "379771at33208", "628247at33208", "267518at33208", "187841at33208", "542676at33208", "500261at33208", "590432at33208", "388256at33208", "274710at33208", "514121at33208", "491289at33208", "323399at33208", "470523at33208", "455512at33208", "445034at33208", "556189at33208", "299250at33208", "442823at33208", "242609at33208", "580062at33208", "397019at33208", "573237at33208", "460044at33208", "522886at33208", "493505at33208", "619326at33208", "515410at33208", "229310at33208", "572463at33208", "516078at33208", "442535at33208", "365063at33208", "273391at33208", "433606at33208", "436098at33208", "556750at33208", "389527at33208", "647692at33208", "485721at33208", "500282at33208", "515425at33208", "559576at33208", "311438at33208", "131349at33208", "643883at33208", "445091at33208", "178537at33208", "453049at33208", "411755at33208", "334764at33208", "501546at33208", "638672at33208", "411757at33208", "344013at33208", "356334at33208", "240437at33208", "268134at33208", "390192at33208", "569303at33208", "602106at33208", "453052at33208", "208219at33208", "584880at33208", "412930at33208", "405493at33208", "633007at33208", "129179at33208", "453056at33208", "526487at33208", "102804at33208", "247133at33208", "475826at33208", "285966at33208", "522515at33208", "440195at33208", "458238at33208", "590470at33208", "468757at33208", "255242at33208", "471608at33208", "561054at33208", "595010at33208", "517404at33208", "578784at33208", "511668at33208", "278227at33208", "257141at33208", "549988at33208", "636070at33208", "625148at33208", "304777at33208", "103854at33208", "281693at33208", "524165at33208", "260752at33208", "414719at33208", "529708at33208", "240962at33208", "75408at33208", "409186at33208", "567750at33208", "539215at33208", "563426at33208", "505718at33208", "587513at33208", "389367at33208", "234290at33208", "519030at33208", "566447at33208", "644665at33208", "523534at33208", "503828at33208", "63149at33208", "634650at33208", "572845at33208", "598944at33208", "405160at33208", "544919at33208", "625077at33208", "368670at33208", "552764at33208", "640349at33208", "269745at33208", "608190at33208", "339032at33208", "553982at33208", "553983at33208", "288202at33208", "539234at33208", "357580at33208", "606776at33208", "587407at33208", "338482at33208", "450349at33208", "274044at33208", "412542at33208", "631107at33208", "553991at33208", "642323at33208", "487987at33208", "409006at33208", "367252at33208", "45934at33208", "271742at33208", "428787at33208"]

# %% [code]
print(len(single_copy_metazoa))

# %% [code]
# fasta = requests.get('https://v101.orthodb.org/pgrest/rpc/fasta?id=615596at33208&species=7227')

# %% [code]
# with open ("orthoDB/615596at33208.fasta", 'wb') as f:
#     f.write(fasta.content)

# %% [markdown]
Loop through list of OrthoDB cluster id's and for each, use OrthoID's API interface (where this url will output a fasta file) to get fasta files for each gene ID with Drosophila melanogaster sequence:

# %% [code]
for i in single_copy_metazoa:
    url = f"https://v101.orthodb.org/pgrest/rpc/fasta?id={i}&species=7227"
    fasta = requests.get(url)
    with open (f"orthoDB/{i}.fasta", 'wb') as f:
        f.write(fasta.content)

# %% [markdown]
Next: go through each fasta file and get rid of all species except for Drosophila melanogaster

- open file as read
- loop through lines
- if line.startswith('>7227_0:0017ef'), keep this and next line. Delete others

# %% [code]
# this works for one file
with open("orthoDB/123923at33208.fasta", 'r') as fin, open("D_melanogaster/123923at33208.fasta", 'w') as fout:
    for line in fin:
        if line.startswith('>7227_0:0017ef'):
            fout.write(line)
            fout.write(next(fin))
            # data[line] = f"{file[line+1]}"
            # print(data)

# %% [code]
# ran this loop on 7/20/22
count = 0
for i in single_copy_metazoa:
    with open(f"orthoDB/{i}.fasta", 'r') as fin, open(f"D_melanogaster/{i}.fasta", 'w') as fout:
        for line in fin:
            if line.startswith('>7227'):
                # print(line)
                fout.write(line)
                fout.write(next(fin))
                count += 1
print(count)

# %% [markdown]
7/20/22

Next we will generate a list of the pub_gene_id's from each of the 873 genes.

Unfortunately, the format of this ID is not consistent. Only FlyBase ID's can be used to easily get DNA sequence.

# %% [code]
import ast

# %% [code]
# goal: make a list of FlyBase IDs for each gene in the D_melanogaster directory
count = 0
geneIDs = []
for i in single_copy_metazoa:
    with open(f"D_melanogaster/{i}.fasta", 'r') as f: # open each file
        for line in f:
            if line.startswith('>7227'):
                l = line[15:] # remove the first 15 characters
                l = ast.literal_eval(l) # convert to a dictionary
                # now we want to make a list of the pub_gene_ids
                geneIDs.append(l['pub_gene_id'])
                count += 1
print(count) # expect 873, same as above
print(geneIDs)

# %% [code]
FlyBaseIDs = []
for i in geneIDs:
    if 'FB' in i:
        FlyBaseIDs.append(i)
print(FlyBaseIDs)
print(len(FlyBaseIDs))
# 158 out of 873 genes include FlyBase IDs. For now, we can use these to go forward.

# %% [code]
import copy

# %% [code]
#clean up the FlyBase IDs
old_FlyBaseIDs = copy.deepcopy(FlyBaseIDs)
FlyBaseIDs = []
for i in old_FlyBaseIDs:
    # remove all characters except for FBpp and following 7 digits
    for c in range(len(i)): # makes c an int
        if i[c] == "F" and i[c+1] == "B" and i[c+2] == "p" and i[c+3] == "p":
            new = i[c:(c+11)]
            FlyBaseIDs.append(new)
print(FlyBaseIDs)

# %% [code]
# randomly choose 15 FlyBase IDs to use for BLAST analysis against Hypsibius
import random
selection = random.sample(FlyBaseIDs, 15)
print(selection)
```
