## Supplementary material for "Modified meiosis in the tardigrade *Hypsibius exemplaris* maintains heterozygosity across the genome": File S1: Plot_Heterozygosity_Data.R

```
library(plyr)
library(nlme)
library(scales)
library(ggplot2)
library(grid)
library(gridExtra)
library(data.table)
library(dplyr)
library(ggbreak)

###############################################################################
# Specify File locations
###############################################################################

# summary files are downloaded locally before plotting
main_dir = "C:/Users/addie/OneDrive - University of North Carolina at Chapel Hill/LAB/tardigrade_heterozygosity/all_refs_heterozygosity_results"
setwd(main_dir)

plots_dir = "C:/Users/addie/OneDrive - University of North Carolina at Chapel Hill/LAB/tardigrade_heterozygosity/He_FIGURE_PLOTS"

###############################################################################
# Adjust settings for ALL plots
###############################################################################

# theme settings
theme_set(theme_classic())
theme_update(text=element_text(size=10),
             strip.background = element_blank(), # remove box around facet labels
             strip.text = element_text(face = "bold"), # and bold text
             legend.key.size = unit(0.1, "in") # adjust legend sizing
)

cbPalette <- c("#999999", "#56B4E9", "#E69F00", "#CC79A7", "#009E73")

###############################################################################
# Read in & Organize Data Files
###############################################################################

### Heterozygosity Summary Data:

## per sample, HiC & NCBI reference genomes, each with & without CDS filter applied
fileName <- "heterozygosity_summary_data.csv"
heterozygosity_summary_all_reps = read.csv(fileName, header=T)
head(heterozygosity_summary_all_reps)
summary(heterozygosity_summary_all_reps)

## four-fold degenerate sites filter, NCBI reference only
fileName <- "four_fold_degenerate_heterozygosity_summary_data.csv"
ffd_heterozygosity_summary_all_reps = read.csv(fileName, header=T)
head(ffd_heterozygosity_summary_all_reps)

# prepare to combine dataframes
ffd_heterozygosity_summary_all_reps$CDS_filter <- FALSE
ffd_heterozygosity_summary_all_reps$four_fold_degenerate_filter <- TRUE
heterozygosity_summary_all_reps$four_fold_degenerate_filter <- FALSE

# combine dataframes
heterozygosity_summary_all_reps <- rbind(heterozygosity_summary_all_reps, ffd_heterozygosity_summary_all_reps)


### Binned Heterozygosity Data: HiC genome only; choose either CDS filtered or unfiltered & bin size
bin_size = 100000
## CDS filtered
fileName <- paste0("tardigrade_HiC_heterozygosity_all_samples_CDS_filter_TRUE_",bin_size/1000,"kb_bins.csv")
CDSonly_heterozygosity_binned_all_reps = read.csv(fileName, header=T)
## CDS unfiltered
fileName <- paste0("tardigrade_HiC_heterozygosity_all_samples_CDS_filter_FALSE_",bin_size/1000,"kb_bins.csv")
heterozygosity_binned_all_reps = read.csv(fileName, header=T)
## Prep for plotting
CDSonly_heterozygosity_binned_all_reps$scaffold = factor(CDSonly_heterozygosity_binned_all_reps$scaffold,
                                      levels = c("HiC_scaffold_1", "HiC_scaffold_2", "HiC_scaffold_3", "HiC_scaffold_4", "HiC_scaffold_5"),
                                      labels = c('chromosome 1', 'chromosome 2', 'chromosome 3', 'chromosome 4', 'chromosome 5')
)
heterozygosity_binned_all_reps$scaffold = factor(heterozygosity_binned_all_reps$scaffold,
                                                         levels = c("HiC_scaffold_1", "HiC_scaffold_2", "HiC_scaffold_3", "HiC_scaffold_4", "HiC_scaffold_5"),
                                                         labels = c('chromosome 1', 'chromosome 2', 'chromosome 3', 'chromosome 4', 'chromosome 5')
)
# convert bin to position by multiplying bin * bin_size
CDSonly_heterozygosity_binned_all_reps$position = CDSonly_heterozygosity_binned_all_reps$bin*bin_size
CDSonly_heterozygosity_binned_all_reps$position_Mb = CDSonly_heterozygosity_binned_all_reps$position/1000000
heterozygosity_binned_all_reps$position = heterozygosity_binned_all_reps$bin*bin_size
heterozygosity_binned_all_reps$position_Mb = heterozygosity_binned_all_reps$position/1000000
## View data
head(CDSonly_heterozygosity_binned_all_reps)
head(heterozygosity_binned_all_reps)


### Telomere BLAST Data: results of BLASTing telomere repeat against HiC genome
telomere_BLAST_data = read.table("He_telomere_repeats_vs_HiC_genome.tsv", header=F)
## Prep for plotting
telomere_BLAST_data <- telomere_BLAST_data %>%
  setnames(old = c("V1", "V2", "V3", "V4", "V5", "V6", "V7", "V8", "V9", "V10", "V11", "V12"),
           new = c("query", "scaffold", "percentage_match", "alignment_length", "N_mismatches", "N_gaps",
                   "query_start", "query_end", "database_start", "database_end", "Evalue", "bitscore")) %>%
  subset(scaffold=="HiC_scaffold_1" | scaffold=="HiC_scaffold_2" | scaffold=="HiC_scaffold_3" | scaffold=="HiC_scaffold_4" | scaffold=="HiC_scaffold_5")
telomere_BLAST_data$scaffold = factor(telomere_BLAST_data$scaffold,
                                      levels = c("HiC_scaffold_1", "HiC_scaffold_2", "HiC_scaffold_3", "HiC_scaffold_4", "HiC_scaffold_5"),
                                      labels = c('chromosome 1', 'chromosome 2', 'chromosome 3', 'chromosome 4', 'chromosome 5')
)
head(telomere_BLAST_data)

###############################################################################
# Summarize Data Across Replicates
###############################################################################

### Summary Data ###
heterozygosity_summary <- heterozygosity_summary_all_reps %>%
  group_by(species, reference, method, CDS_filter, four_fold_degenerate_filter) %>%
  summarise(
    mean_heterozygosity_fraction = mean(heterozygosity_fraction),
    sd_heterozygosity_fraction = sd(heterozygosity_fraction),
    se_heterozygosity_fraction = sd(heterozygosity_fraction)/sqrt(n()),
    grand_mean_coverage = mean(mean_coverage),
    sd_coverage = sd(mean_coverage),
    grand_mean_quality = mean(mean_quality),
    sd_quality = sd(mean_quality),
    .groups = "drop"
  )
head(heterozygosity_summary)

### Binned Data ###
## CDS filtered
CDSonly_heterozygosity_binned <- CDSonly_heterozygosity_binned_all_reps %>%
  group_by(species, reference, method, CDS_filter, scaffold, bin, position, position_Mb) %>%
  summarise(
    mean_heterozygosity_fraction = mean(heterozygosity_fraction),
    sd_heterozygosity_fraction = sd(heterozygosity_fraction),
    mean_minor_allele_freq = mean(mean_minor_allele_freq),
    sd_minor_allele_freq = sd(mean_minor_allele_freq),
    grand_mean_coverage = mean(mean_coverage),
    sd_coverage = sd(mean_coverage),
    grand_mean_quality = mean(mean_quality),
    sd_quality = sd(mean_quality),
    .groups = "drop"
  )
head(CDSonly_heterozygosity_binned)
## no CDS filter
heterozygosity_binned <- heterozygosity_binned_all_reps %>%
  group_by(species, reference, method, CDS_filter, scaffold, bin, position, position_Mb) %>%
  summarise(
    mean_heterozygosity_fraction = mean(heterozygosity_fraction),
    sd_heterozygosity_fraction = sd(heterozygosity_fraction),
    mean_minor_allele_freq = mean(mean_minor_allele_freq),
    sd_minor_allele_freq = sd(mean_minor_allele_freq),
    grand_mean_coverage = mean(mean_coverage),
    sd_coverage = sd(mean_coverage),
    grand_mean_quality = mean(mean_quality),
    sd_quality = sd(mean_quality),
    .groups = "drop"
  )
head(heterozygosity_binned)

###############################################################################
# Make Plots
###############################################################################
## compare 4 single-tardigrade replicates, no CDS filter, both genomes:

plotting_data <- heterozygosity_summary_all_reps %>%
  subset(method == "individual" & CDS_filter == FALSE)
# add pseudo-sample that is mean across 4 reps to plotting_data
pseudo_sample <- heterozygosity_summary %>%
  subset(method == "individual" & CDS_filter == FALSE)
pseudo_sample$replicate <- "Average"
# rename pseudo-sample columns to match plotting_data
colnames(pseudo_sample) <- c("species", "reference", "method", "CDS_filter",
                             "four_fold_degenerate_filter",
                             "heterozygosity_fraction", "heterozygosity_sd",
                             "heterozygosity_se",
                             "mean_coverage", "sd_coverage",
                             "mean_quality", "sd_quality", "replicate")
# combine 
plotting_data$heterozygosity_se <- NA # set se to NA for individual replicates since we are not plotting error bars for them
plotting_data <- rbind(plotting_data, pseudo_sample)
# flag pseudo-sample for distinct appearance in plot
plotting_data$is_combined <- plotting_data$replicate == "Average"
# add unique key for reference, ffd filter combinations
plotting_data$key <- paste0(plotting_data$reference, "_", plotting_data$four_fold_degenerate_filter)


## plot heterozygosity per rep vs average
ggplot(plotting_data, aes(x = factor(replicate),
                          y = heterozygosity_fraction*100,
                          # fill = reference,
                          fill = key,
                          alpha = is_combined)) +
  geom_bar(stat="identity", position=position_dodge(),
           color = "grey20", aes(linetype = is_combined)) + #outline
  # error bars (se)
  geom_errorbar(aes(ymin=heterozygosity_fraction*100 - heterozygosity_se*100,
                    ymax=heterozygosity_fraction*100 + heterozygosity_se*100),
                position=position_dodge(width=0.9), width=0.25, color="grey20") +
  # add labels to bars - above error bars if present
  geom_text(aes(y = heterozygosity_fraction*100 + coalesce(heterozygosity_se, 0)*100,
                label=sprintf("%.2f", heterozygosity_fraction*100)),
            position=position_dodge(width=0.9),
            vjust=-0.25,
            size=2.8) +
  scale_fill_manual(values=c("HiC_FALSE" = "#009E73", "NCBI_FALSE" = "#56B4E9", "NCBI_TRUE" = "#CC79A7"),
                    labels = c("Hi-C assembly", "NCBI assembly", "NCBI, four-fold degenerate sites"))+
  # scale_fill_manual(values=c("HiC" = "#009E73", "NCBI" = "#CC79A7"))+
  scale_alpha_manual(values = c(`FALSE` = 1, `TRUE` = 0.55), guide = "none") +
  scale_linetype_manual(values = c(`FALSE` = "solid", `TRUE` = "dashed"),
                        guide = "none") +
  # expand above labels
  scale_y_continuous(
    expand = expansion(mult = c(NA, 0.05))
  ) +
  labs(x="Single-Tardigrade Replicate", y="Heterozygosity (%)", fill="Reference Genome & Filtering") +
  theme(legend.position = "right")
plot_title <- "het_per_stwgs_replicate_vs_reference_allsites"
ggsave(paste0(plot_title, '.png'), path = plots_dir, width=7, height=3)
ggsave(paste0(plot_title, '.pdf'), path = plots_dir, width=7, height=3)


###############################################################################
# Plot Heterozygosity across the genome
###############################################################################

### Combined replicates ###

## CHOOSE CONDITIONS (except bin size, which is set when reading in data):

# CDS-filtered, mean of 4 sequenced individuals
plotting_data <- CDSonly_heterozygosity_binned %>%
  subset(reference == "HiC" & method == "individual")
summary_data <- heterozygosity_summary %>%
  subset(reference == "HiC" & method == "individual" & CDS_filter == TRUE)
mean_heterozygosity <- summary_data$mean_heterozygosity_fraction*100
paste0("Mean heterozygosity across 4 individuals (CDS only): ", mean_heterozygosity)
plot_title <- paste0("Het_across_genome_mean_of_stwgs_reps_CDSonly_",bin_size/1000,"kbBins")

# OR all sites, mean of 4 sequenced individuals
plotting_data <- heterozygosity_binned %>%
  subset(reference == "HiC" & method == "individual")
summary_data <- heterozygosity_summary %>%
  subset(reference == "HiC" & method == "individual" & CDS_filter == FALSE)
mean_heterozygosity <- summary_data$mean_heterozygosity_fraction*100
paste0("Mean heterozygosity across 4 individuals (all sites): ", mean_heterozygosity)
plot_title <- paste0("Het_across_genome_mean_of_stwgs_reps_allsites_",bin_size/1000,"kbBins")

## Plotting Code
ggplot(plotting_data, aes(x=position_Mb, y=mean_heterozygosity_fraction*100))+
  #geom_line(color='#27AAE1')+
  geom_line(color="grey20")+
  geom_area(fill="#009E73")+
  # geom_smooth(color='grey', alpha=0.8)+
  facet_grid(cols=vars(scaffold), scales='free_x',
             space='free_x')+
  # coord_cartesian(ylim=c(0, 10), # cuts off edge of plot instead of removing data points
  coord_cartesian(ylim=c(0, 6), # cuts off edge of plot instead of removing data points
                  expand=FALSE)+ # make data go to edge of plots
  scale_x_continuous(breaks = c(1,5,10,15,20,25),
                     labels = c('1','5','10','15','20','25'))+
  scale_y_continuous(breaks = c(0, 1, 2, 3, 4, 5, 6),
                     labels = c('   0', '   1', '   2', '   3', '   4', '   5', '   6'))+
  # geom_hline(aes(yintercept=mean_heterozygosity), color='black', linewidth=0.8, linetype="dotted")+
  #geom_hline(aes(yintercept=median_heterozygosity), color='black', linewidth=0.8)+
  labs(x='Genomic Position (Mb)', y='Heterozygosity (%)')+
  theme(panel.spacing = unit(0.3, 'lines')) # reduce space between panels
# Save plot
ggsave(paste0(plot_title, '.png'), path = plots_dir, width=7, height=2)
ggsave(paste0(plot_title, '.pdf'), path = plots_dir, width=7, height=2)


### Separate replicates ###

## all sites, separate stwgs replicates
plotting_data <- heterozygosity_binned_all_reps %>%
  filter(reference == "HiC" & method == "individual")
plot_title <- paste0("Het_across_genome_stwgs_each_rep_allsites_",bin_size/1000,"kbBins")

## Plotting Code
ggplot(plotting_data, aes(x=position_Mb, y=heterozygosity_fraction*100))+
  #geom_line(color='#27AAE1')+
  geom_line(color="grey20")+
  geom_area(fill="#009E73")+
  # geom_smooth(color='grey', alpha=0.8)+
  facet_grid(cols=vars(scaffold), rows = vars(replicate),
             scales='free_x',
             space='free_x')+
  # coord_cartesian(ylim=c(0, 10), # cuts off edge of plot instead of removing data points
  coord_cartesian(ylim=c(0, 6), # cuts off edge of plot instead of removing data points
                  expand=FALSE)+ # make data go to edge of plots
  scale_x_continuous(breaks = c(1,5,10,15,20,25),
                     labels = c('1','5','10','15','20','25'))+
  # geom_hline(aes(yintercept=mean_heterozygosity), color='black', linewidth=0.8, linetype="dotted")+
  #geom_hline(aes(yintercept=median_heterozygosity), color='black', linewidth=0.8)+
  labs(x='Genomic Position (Mb)', y='Heterozygosity (%)')+
  theme(panel.spacing.x = unit(0.3, 'lines'),
        panel.spacing.y = unit(0.6, 'lines'))
# Save plot
ggsave(paste0(plot_title, '.png'), path = plots_dir, width=7, height=5)
ggsave(paste0(plot_title, '.pdf'), path = plots_dir, width=7, height=5)


###############################################################################
# Plot telomere alignments across the genome
###############################################################################

# histogram of count per position, faceted by chromosomes - possibly for figure-making?
ggplot(telomere_BLAST_data)+
  #geom_histogram(aes ( x = database_start), binwidth = 1e+05)+ # matches het plot
  geom_histogram(aes ( x = database_start), binwidth = 1e+06, alpha = 0.5)+ # better aesthetically
  facet_grid(cols=vars(scaffold), scales='free_x',
             space='free_x')+
  coord_cartesian(expand=FALSE)+ # make data go to edge of plots
  scale_x_continuous(breaks = c(1e+06,5e+06,10e+06,15e+06,20e+06,25e+06),
                     labels = c('1','5','10','15','20','25'))+
  labs(x='', y='Telomere Repeats')+
  theme(panel.spacing = unit(0.3, 'lines')) # reduce space between panels
        # axis.line.y = element_blank(),
        # axis.ticks.y = element_blank(),
        # strip.text = element_blank()) # facet labels blank

plot_title <- "telomere_BLAST_histogram_per_position"
ggsave(paste0(plot_title, '.png'), path = plots_dir, width=7, height=2)
ggsave(paste0(plot_title, '.pdf'), path = plots_dir, width=7, height=2)
```
