## Supplementary material for "Modified meiosis in the tardigrade *Hypsibius exemplaris* maintains heterozygosity across the genome": File S1: Plot_High_Impact_Variants_Data.R

```
library(plyr)
library(nlme)
library(scales)
library(ggplot2)
library(grid)
library(gridExtra)
library(data.table)
library(dplyr)
library(ggtext)
library(ggpattern)

###############################################################################
# Specify File locations
###############################################################################

# I download the summary file locally before plotting
main_dir = "C:/Users/addie/OneDrive - University of North Carolina at Chapel Hill/LAB/tardigrade_heterozygosity/NCBI_genome_results"
setwd(main_dir)

plots_dir = "C:/Users/addie/OneDrive - University of North Carolina at Chapel Hill/LAB/tardigrade_heterozygosity/He_FIGURE_PLOTS"

###############################################################################
# Adjust settings for ALL plots
###############################################################################

# theme settings
theme_set(theme_classic())
theme_update(text=element_text(size=10),
             strip.background = element_blank(), # remove box around facet labels
             strip.text = element_text(face = "bold"), # and bold text
             legend.key.size = unit(0.1, "in") # adjust legend sizing
)

cbPalette <- c("#999999", "#56B4E9", "#E69F00", "#CC79A7", "#009E73")

###############################################################################
# Read in & Organize Data File(s)
###############################################################################

## Select high-impact variants file (from Compile_Variant_Summary_Data.R)
fileName <- "select_high_impact_variants.csv"
high_impact_variants = read.csv(fileName, header=T)
head(high_impact_variants)

## Ranked ordered list of genes with high-impact variants based on expression level (from Plot_RNA_Expression_Data.R)
RNAseq_dir <- "C:/Users/addie/OneDrive - University of North Carolina at Chapel Hill/LAB/tardigrade_heterozygosity/rna_expression_analysis_results"
fileName <- paste0(RNAseq_dir, "/high_impact_variant_genes_ranked_by_expression.csv")
high_impact_variant_genes_ranked = read.csv(fileName, header=T)

# ## Protein annotations file(s) (from BLASTp, Uniprot, etc. - not used in final paper figures)
# 
# # fileName <- "../Hypsibius_exemplaris_PredictedProteins.csv"
# fileName <- "../Hypsibius_exemplaris_annotated_genes_with_goslim.csv"
# ProteinAnnotations <- read.csv(fileName, header=T)
# ProteinAnnotations$top_BLASTP_hit <- ifelse(ProteinAnnotations$top_BLASTP_hit == 0, NA, ProteinAnnotations$top_BLASTP_hit)
# ProteinAnnotations$Uniprot_Protein_Annotation <- ifelse(ProteinAnnotations$Uniprot_Protein_Annotation=="Uncharacterized protein", NA, ProteinAnnotations$Uniprot_Protein_Annotation)
# # head(ProteinAnnotations)
# 
# # fileName <- "../H_exemplaris_DNA_Repair_Genes_from_Clark-Hachtel_2024_tables_S3_S4.csv"
# # DNA_Repair_Genes <- read.csv(fileName, header=T)
# # head(DNA_Repair_Genes)

###############################################################################
# Analysis
###############################################################################

### Selection relevant columns & add (optional) protein annotations, TPM rankings ###
mydata <- high_impact_variants %>%
  # left_join(ProteinAnnotations, by = "Gene_ID") %>%
  # left_join(DNA_Repair_Genes %>% select(Gene_ID, Protein, DNA_Repair_Pathway), by = "Gene_ID") %>%
  left_join(high_impact_variant_genes_ranked %>% select(gene_id, mean_TPM, genome_percentile_meanTPM), by = c("Gene_ID" = "gene_id")) %>%
  select(
    Gene_ID,
    Annotation,
    AA_pos,
    # Ensembl_Description,
    # top_BLASTP_hit,
    # Uniprot_Protein_Annotation,
    # Uniprot_subcellular_location,
    # Gene_Ontology,
    # GO_slim
    # Protein,
    # DNA_Repair_Pathway,
    mean_TPM,
    genome_percentile_meanTPM
  )
# head(mydata)

## add column with simplified SnpEff annotation classes
mydata <- mydata %>%
  mutate(
    Annotation_simple = case_when(
      grepl("stop_gained", Annotation) ~ "Stop gained",
      grepl("frameshift_variant", Annotation) ~ "Frameshift",
      grepl("splice_acceptor|splice_donor", Annotation) ~ "Splice site",
      grepl("start_lost|stop_lost", Annotation) ~ "Start/stop lost",
      TRUE ~ "Other high-impact"
    )
  )
head(mydata)

###############################################################################
### Gene counts with variant(s) per annotation category ###
###############################################################################

# prep data for plotting
gene_counts <- mydata %>%
  distinct(Gene_ID, Annotation_simple) %>%
  count(Annotation_simple, name = "n_genes")
variant_counts <- mydata %>%
  count(Annotation_simple, name = "n_variants")
plot_df <- left_join(gene_counts, variant_counts, by = "Annotation_simple")

# plot
ggplot(plot_df, aes(x = reorder(Annotation_simple, n_genes))) +
  # geom_col(aes(y = n_variants), fill = "grey85") +
  geom_col(aes(y = n_genes), fill = "grey40") +
  geom_text(
    aes(label = paste0(n_genes, " (", n_variants, ")"),
        y = n_genes),
    hjust = -0.1,
    size = 3
  ) +
  coord_flip() +
  expand_limits(y = max(plot_df$n_variants) * 1.2) +
  labs(
    x = NULL,
    y = "Number of genes"
    # title = "Genes with predicted high-impact heterozygous variants"
  )

# save
plot_title <- "high_impact_variant_gene_counts_by_annotation"
ggsave(paste0(plot_title, '.png'), path = plots_dir, width=3.5, height=2)
ggsave(paste0(plot_title, '.pdf'), path = plots_dir, width=3.5, height=2)


###############################################################################
### Size changes of early-STOP genes ###
###############################################################################

nonsense <- mydata %>%
  # filter to stop gained variants
  subset(Annotation_simple == "Stop gained") %>%
  # create new columns from AA_pos column
  mutate(
    WT_AA_length = as.integer(sub(".*/", "", AA_pos)),
    nonsense_AA_length = as.integer(sub("/.*", "", AA_pos)),
    trunc_frac = 1 - (nonsense_AA_length / WT_AA_length)
  ) %>%
  # if a gene is in the list twice, filter to shorter nonsense product
  group_by(Gene_ID) %>%
  slice_min(nonsense_AA_length, n = 1, with_ties = FALSE) %>%
  ungroup() %>%
  # make new column that is Gene_ID minus prefix (BV898_)
  mutate(Gene_ID_short = sub("BV898_", "", Gene_ID))
# head(nonsense %>% select(Gene_ID, AA_pos, WT_AA_length, nonsense_AA_length, Uniprot_Protein_Annotation))

## further subset to only genes with significant truncation
nonsense_significant <- nonsense %>%
  filter(trunc_frac >= 0.25)  # keep only proteins truncated >= 25% (change if needed)
# head(nonsense_significant %>% select(Gene_ID, AA_pos, WT_AA_length, nonsense_AA_length, trunc_frac))

## set Gene_ID as a factor with levels sorted by truncation fraction OR mean TPM
nonsense_significant <- nonsense_significant %>%
  # mutate(Gene_ID_short = factor(Gene_ID_short, levels = Gene_ID_short[order(trunc_frac)]))
  mutate(Gene_ID_short = factor(Gene_ID_short, levels = Gene_ID_short[order(mean_TPM)]))

# assuming data is sorted by mean_TPM, find the position where mean_TPM drops below 1 to add a threshold line to the plot
threshold_0_pos <- nonsense_significant %>%
  arrange(mean_TPM) %>%
  mutate(idx = row_number()) %>%
  summarize(pos = max(idx[mean_TPM == 0])) %>%
  pull(pos) + 0.5
threshold_1_pos <- nonsense_significant %>%
  arrange(mean_TPM) %>%
  mutate(idx = row_number()) %>%
  summarize(pos = max(idx[mean_TPM < 1])) %>%
  pull(pos) + 0.5
threshold_10_pos <- nonsense_significant %>%
  arrange(mean_TPM) %>%
  mutate(idx = row_number()) %>%
  summarize(pos = max(idx[mean_TPM < 10])) %>%
  pull(pos) + 0.5
threshold_100_pos <- nonsense_significant %>%
  arrange(mean_TPM) %>%
  mutate(idx = row_number()) %>%
  summarize(pos = max(idx[mean_TPM < 100])) %>%
  pull(pos) + 0.5
y_max <- max(nonsense_significant$WT_AA_length, na.rm = TRUE)

# plot
ggplot(nonsense_significant)+
  # geom_col(aes(x = Gene_ID_short, y = WT_AA_length), fill = "red", width = 0.8)+
  geom_col(aes(x = Gene_ID_short, y = WT_AA_length, fill = mean_TPM), width = 0.8)+
  # geom_col(aes(x = Gene_ID_short, y = nonsense_AA_length), fill = "darkgrey", width = 0.8)+
  geom_col_pattern(
    aes(x = Gene_ID_short, y = nonsense_AA_length),
    pattern = "stripe",
    pattern_fill = "black",
    pattern_angle = 45,
    pattern_density = 0.05,
    pattern_spacing = 0.015,
    fill = NA,
    color = "black",
    linewidth = 0.4,
    width = 0.8
  ) +
  geom_col(aes(x = Gene_ID_short, y = WT_AA_length), color = "black", fill = NA, width = 0.8)+
  # add dotted line(s) to indicate mean_TPM thresholds (computed above)
  geom_vline(xintercept = threshold_1_pos, linetype = "dotted") +  # 1
  # label with mean_TPM
  # geom_text(aes(x = Gene_ID_short, y = WT_AA_length + 50, label = round(mean_TPM, 1)), size = 2.5)+
  coord_flip()+
  # label dotted line(s)
  geom_text(aes(x = threshold_1_pos + 0.7, y = y_max * 0.7),
            label = "mean TPM > 1", hjust = 0, size = 3) +
  scale_fill_gradient(trans = "log10", low = "lightpink", high = "red", na.value = "white",
                      limits = c(0.05, 1001),
                      breaks = c(0.1,1,10,100, 1000),
                      labels = c("0.1", "1", "10", "100", "1000 TPM")) +
  theme(legend.position = c(0.8, 0.12), legend.text = element_text(size = 7)) +
  scale_y_continuous(
    breaks = c(0,500,1000,1500,2000))+
  labs(y = "Nonsense vs Wildtype Protein Length (AA)",
       x = "Locus ID (BV898_#)", fill = "mRNA Abundance")

# save
plot_title <- "nonsense_gene_lengths_significant_truncation_only"
ggsave(paste0(plot_title, '.png'), path = plots_dir, width=3.5, height=4.5)
ggsave(paste0(plot_title, '.pdf'), path = plots_dir, width=3.5, height=4.5)


###############################################################################
### Visualization of frameshift variants ###
###############################################################################

frameshift <- mydata %>%
  # filter to frameshift variants
  subset(Annotation_simple == "Frameshift") %>%
  # create new columns from AA_pos column
  mutate(
    WT_AA_length = as.integer(sub(".*/", "", AA_pos)),
    # nonsense length here means pre-frameshift length; just keeping same labels as above for ease
    nonsense_AA_length = as.integer(sub("/.*", "", AA_pos)),
    trunc_frac = 1 - (nonsense_AA_length / WT_AA_length)
  ) %>%
  # if a gene is in the list twice, filter to shorter nonsense product
  group_by(Gene_ID) %>%
  slice_min(nonsense_AA_length, n = 1, with_ties = FALSE) %>%
  ungroup() %>%
  # make new column that is Gene_ID minus prefix (BV898_)
  mutate(Gene_ID_short = sub("BV898_", "", Gene_ID))
# head(frameshift %>% select(Gene_ID, Gene_ID_short, AA_pos, WT_AA_length, nonsense_AA_length, Uniprot_Protein_Annotation))

## further subset to only genes with significantly early frameshift
frameshift_significant <- frameshift %>%
  filter(trunc_frac >= 0.25)  # keep only proteins with frameshift >= 25% from C-terminus (change if needed)
# head(frameshift_significant %>% select(Gene_ID, AA_pos, WT_AA_length, nonsense_AA_length, trunc_frac))

## set Gene_ID as a factor with levels sorted by truncation fraction OR mean TPM
frameshift_significant <- frameshift_significant %>%
  # mutate(Gene_ID_short = factor(Gene_ID_short, levels = Gene_ID_short[order(trunc_frac)]))
  mutate(Gene_ID_short = factor(Gene_ID_short, levels = Gene_ID_short[order(mean_TPM)]))

# assuming data is sorted by mean_TPM, find the position where mean_TPM drops below 1 to add a threshold line to the plot
threshold_0_pos <- frameshift_significant %>%
  arrange(mean_TPM) %>%
  mutate(idx = row_number()) %>%
  summarize(pos = max(idx[mean_TPM == 0])) %>%
  pull(pos) + 0.5
threshold_1_pos <- frameshift_significant %>%
  arrange(mean_TPM) %>%
  mutate(idx = row_number()) %>%
  summarize(pos = max(idx[mean_TPM < 1])) %>%
  pull(pos) + 0.5
threshold_10_pos <- frameshift_significant %>%
  arrange(mean_TPM) %>%
  mutate(idx = row_number()) %>%
  summarize(pos = max(idx[mean_TPM < 10])) %>%
  pull(pos) + 0.5
threshold_100_pos <- frameshift_significant %>%
  arrange(mean_TPM) %>%
  mutate(idx = row_number()) %>%
  summarize(pos = max(idx[mean_TPM < 100])) %>%
  pull(pos) + 0.5
y_max <- max(frameshift_significant$WT_AA_length, na.rm = TRUE)

# plot
ggplot(frameshift_significant)+
  # geom_col(aes(x = Gene_ID_short, y = WT_AA_length), fill = "blue", width = 0.8)+
  geom_col(aes(x = Gene_ID_short, y = WT_AA_length, fill = mean_TPM), width = 0.8)+
  # geom_col(aes(x = Gene_ID_short, y = nonsense_AA_length), fill = "darkgrey", width = 0.8)+
  geom_col_pattern(
    aes(x = Gene_ID_short, y = nonsense_AA_length),
    pattern = "stripe",
    pattern_fill = "black",
    pattern_angle = 45,
    pattern_density = 0.05,
    pattern_spacing = 0.015,
    fill = NA,
    color = "black",
    linewidth = 0.4,
    width = 0.8
  ) +
  geom_col(aes(x = Gene_ID_short, y = WT_AA_length), color = "black", fill = NA, width = 0.8)+
  # add dotted line(s) to indicate mean_TPM thresholds (computed above)
  geom_vline(xintercept = threshold_1_pos, linetype = "dotted") +  # 1
  # label with mean_TPM
  # geom_text(aes(x = Gene_ID_short, y = WT_AA_length + 50, label = round(mean_TPM, 1)), size = 2.5)+
  coord_flip()+
  # label dotted line(s)
  geom_text(aes(x = threshold_1_pos + 0.7, y = y_max * 0.7),
            label = "mean TPM > 1", hjust = 0, size = 3) +
  scale_y_continuous(
    breaks = c(0,500,1000,1500,2000))+
  scale_fill_gradient(trans = "log10", low = "lightblue", high = "blue", na.value = "white",
                      limits = c(0.05, 1001),
                      breaks = c(0.1,1,10,100, 1000),
                      labels = c("0.1", "1", "10", "100", "1000 TPM")) +
  theme(legend.position = c(0.8, 0.12), legend.text = element_text(size = 7)) +
  labs(y = "Pre-frameshift vs Wildtype Protein Length (AA)",
       x = "Locus ID (BV898_#)", fill = "mRNA Abundance")

# save
plot_title <- "frameshift_gene_lengths_significant_proportion_only"
ggsave(paste0(plot_title, '.png'), path = plots_dir, width=3.5, height=4.5)
ggsave(paste0(plot_title, '.pdf'), path = plots_dir, width=3.5, height=4.5)


# ## save significant nonsense + frameshift genes to file
# significant_truncations <- bind_rows(nonsense_significant, frameshift_significant) %>%
#   arrange(Gene_ID) %>%
#   select(Gene_ID, Annotation, Annotation_simple, AA_pos, WT_AA_length, nonsense_AA_length, trunc_frac, Uniprot_Protein_Annotation)
# # head(significant_truncations)
# # save to file
# write.csv(significant_truncations, "high_impact_variants_significant_truncations.csv", row.names = FALSE)
```
