## Supplementary material for "Modified meiosis in the tardigrade *Hypsibius exemplaris* maintains heterozygosity across the genome": File S1: Plot_RNA_Expression_Data.R

```
library(tidyverse)

###############################################################################
# Read in Summary Files
###############################################################################

# I downloaded files locally before plotting
main_dir = "C:/Users/addie/OneDrive - University of North Carolina at Chapel Hill/LAB/tardigrade_heterozygosity/rna_expression_analysis_results"
setwd(main_dir)

plots_dir = "C:/Users/addie/OneDrive - University of North Carolina at Chapel Hill/LAB/tardigrade_heterozygosity/He_FIGURE_PLOTS"

gene_tpm_ranks_per_sample <- read.csv("gene_tpm_ranks_per_sample.csv")
# head(gene_tpm_ranks_per_sample)
gene_expression_rank_overall <- read.csv("gene_expression_rank_overall.csv")
# head(gene_expression_rank_overall)

fileName <- "select_high_impact_variants.csv"
# genes_with_confident_high_impact_variants <- read.csv("../NCBI_genome_results/20251204_select_high_impact_variants_with_goslim.csv") %>%
genes_with_confident_high_impact_variants <- read.csv(paste0("../NCBI_genome_results/", fileName)) %>%
  select(Gene_ID, Annotation, AA_pos
         # Uniprot_Protein_Annotation, Gene_Ontology, GO_slim
         )
# head(genes_with_confident_high_impact_variants)

# genes_with_significant_truncations <- read.csv("../NCBI_genome_results/high_impact_variants_significant_truncations.csv")

###############################################################################
# Rank-Order Genes with High-Impact Variants by Expression Level
###############################################################################

### Rank-order list of genes with high impact variants by expression level

variant_ids <- unique(genes_with_confident_high_impact_variants$Gene_ID)
n_genome_genes <- nrow(gene_expression_rank_overall)

# Choose the expression metric for ranking (change if you prefer)
metric <- "mean_TPM"    # options: "mean_TPM", "median_TPM", "geo_mean_TPM", "max_TPM"

# Build ranked table of variant genes by the chosen metric
variant_ranked <- gene_expression_rank_overall %>%
  filter(gene_id %in% variant_ids) %>%
  # attach your annotations for context
  left_join(
    genes_with_confident_high_impact_variants %>%
      select(Gene_ID, Annotation,
             # Uniprot_Protein_Annotation, Gene_Ontology, GO_slim,
             AA_pos),
    by = c("gene_id" = "Gene_ID")
  ) %>%
  arrange(desc(.data[[metric]])) %>%
  mutate(
    rank_by_metric = row_number(),
    # genome-wide percentile based on rank_meanTPM if available (otherwise compute from order)
    genome_percentile_meanTPM = if ("rank_meanTPM" %in% names(.)) {
      100 * (1 - (rank_meanTPM - 1) / n_genome_genes)
    } else {
      # fallback percentile based on sorted order within variant set (note: not genome-wide)
      100 * (1 - (rank_by_metric - 1) / n())
    }
  )
head(variant_ranked)

# # add metric to genes with significant truncations
# significant_genes_ranked <- genes_with_significant_truncations %>%
#   left_join(
#     variant_ranked %>%
#       select(gene_id, all_of(metric), genome_percentile_meanTPM),
#     by = c("Gene_ID" = "gene_id")
#   ) %>%
#   arrange(desc(.data[[metric]]))
# head(significant_genes_ranked)

# (Optional) attach average percentile across samples from the per-sample table
variant_avg_percentile <- gene_tpm_ranks_per_sample %>%
  filter(gene_id %in% variant_ids) %>%
  group_by(gene_id) %>%
  summarize(mean_percentile_across_samples = mean(percentile), .groups = "drop")
variant_ranked <- variant_ranked %>%
  left_join(variant_avg_percentile, by = "gene_id")

head(variant_ranked)

# Save the ranked table
out_rank_csv <- "high_impact_variant_genes_ranked_by_expression.csv"
write.csv(variant_ranked, out_rank_csv, row.names = FALSE)
message("[INFO] Saved ranked variant table -> ", out_rank_csv)

###############################################################################
# Plot Aesthetics
###############################################################################

### Adjust settings for ALL plots ###

# theme settings
theme_set(theme_classic())
theme_update(text=element_text(size=10), # IS USUALLY 10
             strip.background = element_blank(), # remove box around facet labels
             strip.text = element_text(face = "bold"), # and bold text
             legend.key.size = unit(0.1, "in") # adjust legend sizing
)

###############################################################################
## Are genes with high-impact variants expressed differently from the rest of the genome?
###############################################################################

eps <- 1e-3

# Histogram (log10 of the chosen metric) for variant genes vs ALL genes
ggplot() +
  # Genome/background distribution
  geom_histogram(
    data = gene_expression_rank_overall,
    aes(x = log10(.data[[metric]] + eps), y = after_stat(density),
        fill = "All genes", color = "All genes", alpha = "All genes"),
    binwidth = 0.2
  ) +
  # Variant set distribution
  geom_histogram(
    data = variant_ranked,
    aes(x = log10(.data[[metric]] + eps), y = after_stat(density),
        fill = "Genes with high-impact variants",
        color = "Genes with high-impact variants",
        alpha = "Genes with high-impact variants"),
    binwidth = 0.2
  ) +
  # adds legend despite using different datasets
  scale_fill_manual(
    values = c(
      "All genes" = "gray70",
      "Genes with high-impact variants" = "#009E73"
    ),
    breaks = c("All genes", "Genes with high-impact variants")  # controls legend order
  ) +
  scale_color_manual(
    values = c(
      "All genes" = "gray70",
      "Genes with high-impact variants" = "#009E73"
    )
  ) +
  scale_alpha_manual(
    values = c(
      "All genes" = 1.0,
      "Genes with high-impact variants" = 0.5
    )
  ) +
  scale_x_continuous(
    breaks = log10(c(0.001, 0.01, 0.1, 1, 10, 100, 1000, 10000)),  # positions on log scale
    labels = c("0", "0.01", "0.1", "1", "10", "100", "1k", "10k")    # what to display
  ) +
  labs(
    # title = paste0("Distribution of variant-containing vs all genes expression (", metric, ")"),
    x = "mRNA Abundance (mean TPM)",
    y = "Proportion of Genes",
    fill = NULL, color = NULL, alpha = NULL
  ) +
  theme(legend.position = c(0.5,0.9))
plot_title <- paste0(metric, "density_distribution_variant_vs_all_genes.png")
ggsave(paste0(plot_title, '.png'), path = plots_dir, width=3.5, height=3)
ggsave(paste0(plot_title, '.pdf'), path = plots_dir, width=3.5, height=3)
```
