## Supplementary material for "Modified meiosis in the tardigrade *Hypsibius exemplaris* maintains heterozygosity across the genome": File S1: README.md

# tardigrade-heterozygosity
Heterozygosity and variant analysis for the tardigrade *Hypsibius exemplaris*, as in preprint: Coke AN, Papell LD, Burch CL, Goldstein B (2026). Modified meiosis in the tardigrade Hypsibius exemplaris maintains heterozygosity across the genome. bioRxiv 2026.03.11.711151 doi.org/10.64898/2026.03.11.711151 [preprint] 

# Code Descriptions

**Identify and analyze heterozygous variants:**

call_filter_and_annotate_variants_pipeline.sh
- Maps short genomic sequencing reads to reference with BWA algorithm
- Calls variants with bcftools
- Filters called variants by depth, quality, and whether they likely represent heterozygous sites
- Annotates variants with SnpEff
- Main outputs: heterozygous_variants.csv and all_variants.csv
- *Usage (per sample):*
   - Single-end reads: ./script.sh <reference.fa> <reads.fastq[.gz]> [snpeff_database snpeff_dir]
   - Paired-end reads: ./script.sh <reference.fa> <reads_1.fastq[.gz]> <reads_2.fastq[.gz]> [snpeff_database snpeff_dir]

identify_synonymous_sites.R
- Identifies four-fold synonymous sites from transcript annotations
- Outputs a .bed file of four-fold generate sites for a subset of transcripts (depending on job submission inputs)
- Combine all sites for downstream use with: $ cat *.bed > all_4_fold_degenerate_sites.bed

Compile_Heterozygosity_Data.R
- Compiles all_variants.csv files generated for multiple samples
- Computes the fraction of heterozygous sites out of total sites post-filtering, both in 100kb bins across putative chromosomes and total across the entire genome
- Calculates heterozygosity at all sites and at four-fold degenerate sites only (the latter requires all_4_fold_degenerate_sites.bed file)
- Main outputs: heterozygosity_summary_data.csv and tardigrade_HiC_heterozygosity_all_samples_CDS_filter_FALSE_100kb_bins.csv

Plot_Heterozygosity_Data.R
- Reads in heterozygosity_summary_data.csv (heterozygosity per sample), tardigrade_HiC_heterozygosity_all_samples_CDS_filter_FALSE_100kb_bins.csv (heterozygosity across the genome), and He_telomere_repeats_vs_HiC_genome.tsv (result of BLAST+ of telomeric repeats against Hi-C genome)
- Makes plots of heterozygosity per sample and across putative chromosomes, as well as telomere BLAST alignment across putative chromosomes

Compile_Variant_Summary_Data.R
- Compiles heterozygous_variants.csv files generated for multiple samples
- Selects SnpEff annotation subfields of interest
- Generates summary file of variants counts per SnpEff annotation category: SnpEff_variant_counts.csv
- Filters variant list to those that are high impact and shared between all three bulk replicates and at least one single-animal replicate for downstream analyses: select_high_impact_variants.csv

Plot_High_Impact_Variants_Data.R
- Reads in select_high_impact_variants.csv
- Makes plots to categorize and analyze select high-impact variants

**Analyze RNA expression of select variants:**

RNA_expression_analysis_pipeline.sh
- Uses Salmon to map and analyze mRNA expression data from raw reads
- Indicate fastq read files and sample names, as well as reference fasta and GFF3, in script
- Outputs are used for analysis by tximport in Compile_RNA_Expression_Data.R

Compile_RNA_Expression_Data.R
- Analyzes mRNA expression data from Salmon outputs using tximport (calculates TPM and other metrics)
- Outputs: gene_tpm_ranks_per_sample.csv, gene_expression_rank_overall.csv, gene_tpm_matrix.csv

Plot_RNA_Expression_Data.R
- Reads in gene_tpm_ranks_per_sample.csv and gene_expression_rank_overall.csv from Compile_RNA_Expression_Data.R
- Reads in select_high_impact_variants.csv from Compile_Variant_Summary_Data.R
- Plots overlaid distribution of RNA expression values to compare select high-impact variants and all genes in genome

allele_specific_analysis_pipeline.sh
- Uses ASEQ to analyze allele-specific expression of transcripts with heterozygous variants of interest

High_Impact_Variants_ASE_Analysis.R
- Statistical analysis of allele-specific expression of genes containing high-impact variants
- Reads in outputs from allele_specific_analysis_pipeline.sh and Compile_Variant_Summary_Data.R

**Analyze results of PCR sequencing and inheritance tracking of select loci:**

parse_OrthoDB_results,ipynb
- custom python script to randomly select loci from OrthoDB results (conserved single-copy and present genes in *Drosophila melanogaster*) for downstream work in *Hypsibius exemplaris*

OrthoDB_Loci_Heterozygosity_Plotting.R
- Used to generate bar plot of heterozygosity per PCR-sequenced locus and compare to genome-wide values

Make_Pedigrees.R
- Plotting code to generate tardigrade family tree pedigrees of allele detection from spreadsheet-input data

Analyze_Heterozygosity_at_Chr_Ends.R
- Analyze Plasmidsaurus Premium PCR sequencing results to determine how many sites out of the total PCR product lengths are heterozygous (aka differ between alleles).

Chr_Ends_Heterozygosity_Plotting.R
- Used to generate bar plot of heterozygosity per PCR-sequenced locus. Reads in output file from Analyze_Heterozygosity_at_Chr_Ends.R

# References for Tools Used

Li, H. Aligning sequence reads, clone sequences and assembly contigs with BWA-MEM. Preprint at https://doi.org/10.48550/arXiv.1303.3997 (2013). 

Danecek, P. et al. Twelve years of SAMtools and BCFtools. Gigascience 10, giab008 (2021). 

Cingolani, P. et al. A program for annotating and predicting the effects of single nucleotide polymorphisms, SnpEff: SNPs in the genome of Drosophila melanogaster strain w1118 ; iso-2; iso-3. Fly 6, 80–92 (2012). 

Patro, R., Duggal, G., Love, M. I., Irizarry, R. A. & Kingsford, C. Salmon provides fast and bias-aware quantification of transcript expression. Nat Methods 14, 417–419 (2017). 

Soneson, C., Love, M. I. & Robinson, M. D. Differential analyses for RNA-seq: transcript-level estimates improve gene-level inferences. Preprint at https://doi.org/10.12688/f1000research.7563.2 (2016). 

Kuznetsov, D. et al. OrthoDB v11: annotation of orthologs in the widest sampling of organismal diversity. Nucleic Acids Research 51, D445–D451 (2023). 

Romanel A, Lago S, Prandi D, Sboner A, Demichelis F. 2015. ASEQ: fast allele-specific studies from next-generation sequencing data. BMC Med Genomics. 8(1):9.
