## Supplementary material for "Modified meiosis in the tardigrade *Hypsibius exemplaris* maintains heterozygosity across the genome": File S1: RNA_expression_analysis_pipeline.sh

```
#!/bin/bash
set -euo pipefail
trap 'echo "Error on line $LINENO: $BASH_COMMAND"; exit 1' ERR

# Load modules
module load gffread
module load salmon

###############################################
# USER SETTINGS
###############################################

# Set working directory
cd /work/users/a/d/adricoke/single_tardigrade_seq_analysis
if [ ! -d rna_expression_analysis_results ]; then
    mkdir rna_expression_analysis_results
fi
cd rna_expression_analysis_results
rna_workdir=$(pwd)

# Input Files: Reference Genome and Gene Annotation
REF=/work/users/a/d/adricoke/single_tardigrade_seq_analysis/refs/Hypsibius_exemplaris/NCBI/GCA_002082055.1/GCA_002082055.1_nHd_3.1_genomic.fna
GFF=/work/users/a/d/adricoke/single_tardigrade_seq_analysis/refs/Hypsibius_exemplaris/NCBI/GCA_002082055.1/genomic.gff

# Input Files: RNA-seq Data
# imporant: assumes paired-end reads in format SAMPLE_1.fastq[.gz] and SAMPLE_2.fastq[.gz]
READS_DIR=/work/users/a/d/adricoke/single_tardigrade_seq_analysis/data/Yoshida2017_RNAseq_reads_10K_Active
declare -a RNASEQ_SAMPLES=("SRR5218239" "SRR5218240" "SRR5218241")
declare -a SAMPLE_NAMES=("Yoshida2017_Active_1" "Yoshida2017_Active_2" "Yoshida2017_Active_3")
# File extension for reads (set to ".fastq" or ".fastq.gz")
READ_EXT=".fastq"   # change to ".fastq.gz" if gzipped

# Salmon settings
THREADS=8
OUTDIR="salmon"
INDEX_DIR="salmon_index"

mkdir -p "${OUTDIR}"

###############################################
# GENERATE TRANSCRIPTOME FASTA FOR SALMON (ONCE)
###############################################

if [[ ! -f "transcripts.fa" ]]; then
  echo "[INFO] Building transcripts.fa from GFF + reference..."
  gffread "${GFF}" -g "${REF}" -w "transcripts.fa"
else
  echo "[INFO] transcripts.fa already exists; skipping."
fi

###############################################
# GENERATE TRANSCRIPT-TO-GENE MAPPING FILE FOR DOWNSTREAM ANALYSIS (ONCE)
###############################################

if [[ ! -f "tx2gene.csv" ]]; then
    echo
    echo "[INFO] Generating tx2gene.csv from GFF..."

    awk -F'\t' '
    $3=="mRNA" || $3=="transcript" {
    tx=""; gene="";
    n=split($9, a, /;/);
    for (i=1;i<=n;i++){
        split(a[i], kv, /=/);
        if (kv[1]=="ID") tx=kv[2];
        if (kv[1]=="Parent") gene=kv[2];
    }
    if (tx!="" && gene!="") print tx","gene;
    }' "${GFF}" \
    | sort -u \
    | (echo "transcript_id,gene_id"; cat) > tx2gene.csv

else
    echo
    echo "[INFO] tx2gene.csv exists; skipping."
fi

###############################################
# BUILD SALMON INDEX FOR QUANTIFICATION (ONCE)
###############################################

# -i: index output directory
# -k: k-mer size (default 31 for reads >=75bp)

if [[ ! -d "${INDEX_DIR}" ]]; then
  echo "[INFO] Building Salmon index in ${INDEX_DIR}..."
  salmon index \
    -t "transcripts.fa" \
    -i "${INDEX_DIR}" \
else
  echo "[INFO] Salmon index ${INDEX_DIR} exists; skipping."
fi

###############################################
# QUANTIFY RNA-SEQ SAMPLES WITH SALMON
###############################################

# -A: automatic library type detection
# -1: reads file 1 (if paired-end)
# -2: reads file 2 (if paired-end)
# OR -r : reads file (if single-end)
# -p: number of threads
# -o: output directory

# PER SAMPLE
for idx in "${!RNASEQ_SAMPLES[@]}"; do
  sample="${RNASEQ_SAMPLES[$idx]}"
  sample_name="${SAMPLE_NAMES[$idx]}"

  # Construct read paths
  reads_1="${READS_DIR}/${sample}_1${READ_EXT}"
  reads_2="${READS_DIR}/${sample}_2${READ_EXT}"

  # Validate inputs
  if [[ ! -f "${reads_1}" ]]; then
    echo "[ERROR] Missing R1: ${reads_1}" >&2
    exit 1
  fi
  if [[ ! -f "${reads_2}" ]]; then
    echo "[ERROR] Missing R2: ${reads_2}" >&2
    exit 1
  fi

  outdir="${OUTDIR}/${sample_name}"
  mkdir -p "${outdir}"

  if [[ -f "${outdir}/quant.sf" ]]; then
    echo "[INFO] ${sample_name}: quant.sf exists; skipping."
    continue
  fi

  echo
  echo "[INFO] Quantifying sample: ${sample_name}"
  salmon quant \
    -i "${INDEX_DIR}" \
    -l A \
    -1 "${reads_1}" \
    -2 "${reads_2}" \
    -p "${THREADS}" \
    -o "${outdir}"
done

###############################################
# END OF PIPELINE
###############################################

echo
echo "[INFO] All quantifications complete."
echo "[INFO] Next step: run R script for tximport + DESeq2."
```
